# Confidence Without Verification: Screening pLDDT Unreliability in AlphaFold2 Fold-Switching Predictions

**DOI:** 10.64898/2026.02.19.706878

**Authors:** R. Thacker

## Abstract

AlphaFold2 (AF2) has transformed structural biology, yet its confidence metrics, particularly the predicted Local Distance Difference Test (pLDDT) and predicted Template Modelling score (pTM), systematically fail for fold-switching proteins, which adopt two or more distinct conformations from a single amino acid sequence. Multiple groups have called for alternative quality measures to address this limitation. Here, we present a confidence-integrity screening method derived from the SDI framework [Thacker, 2025a], originally developed to detect reasoning failures across artificial intelligence architectures. We apply this screen to 27,098 predictions across 18 experimentally validated fold-switching proteins from the Porter fold-switching benchmark [Sala et al., 2023, Lee et al., 2025], identifying a 33.6% high-confidence FalseVerify rate, defined as predictions where AF2 is simultaneously confident about its output and structurally committed to the dominant fold while failing to capture the known alternative conformation. FalseVerify severity is predictable: complete secondary structure refolding produces 80–97% FalseVerify rates, while local backbone rearrangements produce 0-2%. Per-residue analysis localizes confidence failures to specific fold-switching regions, with cross-cluster pLDDT variance serving as a structural fingerprint distinguishing reliable from unreliable predictions even when mean pLDDT values are indistinguishable. Applied blindly to 52 unvalidated *E. coli* fold-switching candidates from the CF-random proteome search, the taxonomy produces structured categories, not noise, with a perfect 50/50 directional split confirming zero population-level correlation between pLDDT and conformational accuracy. The blind screen also identifies a sixth category, INVERTED confidence, invisible in benchmark data, in which the alternative conformation is more confident than the dominant. Nine *E. coli* proteins with balanced confidence profiles are prioritized for experimental validation. These results fill a specific methodological gap identified by Schafer & Porter (2025) and provide a quantitative framework for triaging fold-switching predictions before experimental validation.

## 1 Introduction

The central dogma of structural biology, the idea that one sequence yields one structure, has been challenged by the discovery of fold-switching proteins, which remodel their secondary and tertiary structures in response to cellular stimuli [Chakravarty and Porter, 2022, 2025]. These shapeshifting proteins play critical roles across all kingdoms of life, from bacterial transcription regulation (RfaH) to circadian clock control (KaiB) to cell division checkpoints (Mad2) [Porter, 2023].

AlphaFold2 (AF2) predicts protein structures with remarkable accuracy for most globular proteins [Jumper et al., 2021], but systematically struggles with fold-switching proteins [Chakravarty and Porter, 2022, Bonin et al., 2024]. Several lines of evidence suggest these failures arise from structure memorization: AF2’s Evoformer module memorizes dominant conformations from the training set rather than learning generalizable protein physics [Chakravarty et al., 2024, Sala et al., 2023]. This memorization produces a particularly dangerous failure mode (high-confidence predictions of incorrect structures) that existing quality metrics cannot detect.

The confidence gap has not gone unnoticed. Porter and colleagues demonstrated that pLDDT does not effectively discriminate between experimentally consistent and inconsistent structures [Chakravarty et al., 2024, Schafer and Porter, 2025]. Bryant and Noé found only weak correlation (Pearson *R* = 0.52) between pLDDT and structural accuracy for alternative conformations [Bryant and Noé, 2024]. Most recently, Schafer and Porter explicitly called for “alternative measures” to replace pLDDT for fold-switching assessment, identifying this as an open problem requiring new methodology [Schafer and Porter, 2025].

Several computational methods now generate alternative conformational predictions, including AFcluster [Wayment-Steele et al., 2024], SPEACH_AF [Stein and McHaourab, 2022], CF-random [Lee et al., 2025], CFold [Bryant and Noé, 2024], and AlphaFold2-RAVE [Vani et al., 2023]. However, none of these methods screen the confidence integrity of their output. The question “did we find an alternative conformation?” is increasingly answerable; the question “should we trust this alternative conformation?” remains unaddressed.

Here we present a confidence integrity screen derived from the SDI framework [Thacker, 2025a], which provides a failure taxonomy originally developed for detecting reasoning failures across AI architectures. We do not propose a new protein folding method or a replacement for AF2; we provide a diagnostic screen for assessing when AF2’s confidence output should not be trusted. Applied to 27,098 predictions across 18 experimentally validated proteins, the screen produces a quantitative taxonomy of confidence integrity that predicts which proteins and which residues harbor unreliable AF2 confidence scores, filling the specific methodological gap identified by Schafer and Porter.

## 2 Background and Motivation

### 2.1 AlphaFold2 and Fold Switching

AF2 generates per-residue confidence estimates (pLDDT, range 0–100) alongside predicted structures, with scores above 90 considered very high confidence, 70–90 confident, 50–70 low confidence, and below 50 indicating probable disorder [Jumper et al., 2021, Varadi et al., 2022]. For single-conformation proteins, pLDDT is well-calibrated: it correlates strongly with actual structural accuracy [Terwilliger et al., 2024]. However, for fold-switching proteins, the calibration breaks down. Chakravarty et al. showed that AF2 predictions of fold-switched conformations are driven by structure memorization [Chakravarty et al., 2024]. When CFold, an AF2 variant trained without alternative conformations, was tested on fold switchers, it failed to predict most alternative folds, confirming that AF2’s apparent success depends on having seen the answer during training [Schafer and Porter, 2025].

Current methods for generating alternative conformational predictions share a common limitation: they produce candidates but provide no mechanism for assessing whether AF2’s confidence scores are meaningful for those candidates. CF-random, which identified up to 5% of *E. coli* proteins as potential fold switchers [Lee et al., 2025], generates predictions that urgently require confidence screening before expensive experimental validation.

### 2.2 The SDI Framework

The Structural Dependencies of Intelligence (SDI) framework [Thacker, 2025a] identifies three structural dependencies required for reliable inference in computational systems: Coherence (*C*), internal consistency of outputs; Preservation (*P*), maintenance of validated knowledge; and Emergence (*E*), generation of novel inferences. These dependencies obey an ordered constraint, denoted *λ*[*C* ≥ *P* ≥ *E*]: Preservation cannot exceed Coherence, and Emergence cannot exceed Preservation. This is not a heuristic ordering but a formal constraint: violations produce characteristic, classifiable failure modes rather than random errors. The constraint was validated across 28+ AI architectures with zero observed violations in 420+ evaluations [Thacker, 2025a].

The framework classifies system outputs into four categories based on the relationship between confidence and accuracy: *Correct* (high confidence, accurate output), *False Verify* (high confidence, inaccurate output, the most dangerous mode because it is invisible to the user), *Self Catch* (low confidence, inaccurate output), and *OverRefusal* (low confidence, accurate output).

FalseVerify occurs when the Emergence component exceeds the Preservation component, constituting an *E* > *P* violation. The system generates novel output but lacks the internal checks to distinguish correct novelty from confabulation.

### 2.3 Mapping to Protein Folding

The SDI failure taxonomy maps directly onto AF2’s treatment of fold-switching proteins: Coherence maps to internal consistency of the predicted structure (bond geometry, steric compliance); Preservation maps to accuracy of the dominant fold prediction (maintained from training data); and Emergence maps to generation of alternative conformations (novel structural inference beyond training).

FalseVerify in the protein context is a prediction that (a) achieves high pLDDT (the system is confident), (b) accurately reproduces the dominant fold (Preservation is satisfied), but (c) completely fails to capture the alternative conformation (Emergence exceeds Preservation, meaning the system’s confidence extends beyond its actual structural knowledge).

This mapping generates a testable prediction: *False Verify severity should scale with the magnitude of the gap between what AF2 has memorized (training-set structures) and what physics requires (the alternative fold)*. Complete secondary structure refolding should produce severe FalseVerify; local backbone rearrangements should produce little or none.

## 3 Methods

### 3.1 Datasets

#### Dataset 1: Porter Fold-Switching Benchmark

We analyzed 18 proteins with experimentally determined structures for both dominant and alternative conformations, drawn from the Porter fold-switching benchmark [Chakravarty et al., 2024, Lee et al., 2025]. Each protein was represented by CF-random predictions with associated pLDDT percentile scores and TM-scores computed against both reference structures (fold 1 and fold 2). The dataset contained 33 CSV files totaling 27,098 individual predictions. The 18 proteins span diverse fold-switching mechanisms: complete secondary structure refolding (KaiB, RfaH, GP2), domain rearrangement (PimA, FraC, Nrp2), local backbone changes (Mad2, COMT, OxyR), and rigid body motions (Cas9, CRKL, IscA).

#### Dataset 2: E. coli Blind Candidates

We analyzed 52 putative fold-switching proteins from the CF-random blind search of the *E. coli* proteome [Lee et al., 2025]. These proteins were identified by an automated pipeline with no prior experimental validation. Each protein was represented by a PyMOL session (PSE) file containing two conformations (dominant and alternative) with per-residue pLDDT in the B-factor column.

### 3.2 Confidence Integrity Classification

For each prediction, we classified the relationship between confidence and structural accuracy using TM-score thresholds. A prediction was labeled *Fold1-only* if TM_foldl_ ≥ 0.5 and TM_fold2_ < 0.5, *Fold2-only* if the reverse held, *Both folds* if both TM-scores exceeded 0.5, and *Neither* if both fell below 0.5.

The *high-confidence (HC) subset* was defined as predictions with pLDDT percentile ≥ 50. The *False Verify (FV) rate* was computed as:

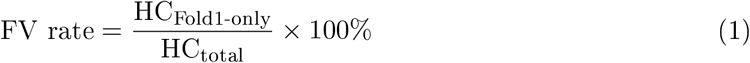

The primary analysis uses TM ≥ 0.5 (the field-standard threshold; *P*-value 5.5 × 10^−7^; Xu and Zhang 2010). To assess sensitivity to threshold choice, we repeated the full classification at TM ≥ 0.6 (Schafer & Porter’s success criterion), TM ≥ 0.7, and TM ≥ 0.8 (Chakravarty & Porter’s “good prediction” criterion).

### 3.3 Confidence-Accuracy Correlation

For each protein, we computed the Pearson correlation coefficient *R* between pLDDT percentile and TM-score against the alternative fold (TM_fold2_). We also computed the confidence gap Δ_pLDDT_ = mean pLDDT_Foldl-only_ − mean pLDDT_Fold2-only_.

### 3.4 Per-Residue Analysis

For five representative proteins spanning the FalseVerify spectrum (KaiB, PimA, FraC, OxyR, Mad2), we extracted per-residue pLDDT values from the B-factor columns of AF2 prediction PDB files. For each protein, we computed: (i) *cross-cluster pLDDT variance*, defined as the variance in pLDDT across different MSA clusters’ best predictions for each residue position, the variance in pLDDT across different MSA clusters’ best predictions; (ii) *within-cluster rank delta*, the difference in per-residue pLDDT between rank-1 and rank-5 models; and (iii) *high-variance residue identification*, identifying residues with cross-cluster variance > 100.

### 3.5 Pattern Classification

Based on the FV rate, correlation coefficient, and fraction of predictions capturing both folds, each protein was assigned to one of five confidence integrity categories: SEVERE FV (FV > 80%), MODERATE FV (FV 40–80%), MILD FV (FV 20–40%), BOTH FOLDS (> 30% predictions match both folds), and MAINTAINED (FV < 20%, *R* > 0).

### 3.6 Blind Screening of *E. coli* Candidates

For each of 52 *E. coli* candidates, we computed mean pLDDT for dominant and alternative conformations and classified into: HIGH FV RISK (dom_pLDDT > 70, Δ > +15), MODERATE FV RISK (Δ = +8 to +15), BOTH STABLE (|Δ| < 5), INVERTED (Δ < −10), and OTHER (remaining cases).

## 4 Results

### 4.1 Population-Level FalseVerify Rates

Across all 27,098 predictions from 18 fold-switching proteins (33 reference comparisons), we observed a population-level HC FalseVerify rate of 33.6% (3,802 of 11,301 high-confidence predictions). An additional 36.2% (4,092/11,301) of HC predictions captured both folds simultaneously (Table 1).

**Table 1:** Confidence integrity classification of 18 fold-switching proteins. N_pred_ = total predictions across all references. FV% averaged across references. *R* values shown as PDB1/PDB2 where two references exist. *CaBP shows both high FV and high Both-folds, indicating a boundary case.

| Protein | Full Name | $N_{\text{pred}}$ | FV% | $R$ | %Both | Pattern |
| --- | --- | --- | --- | --- | --- | --- |
| GP2 | Viral fusion glycoprotein | 290 | 97.5 | -0.08/+0.38 | 2.6 | SEVERE |
| KaiB | Circadian clock protein | 2,765 | 88.2 | -0.12/+0.08 | 0.0 | SEVERE |
| PimA | Mannosyltransferase | 2,643 | 93.1 | +0.55/+0.55 | 0.0 | SEVERE |
| FraC | Fragaceatoxin C | 470 | 57.7 | +0.13/-0.09 | 0.0 | MODERATE |
| Nrp2 | Neuropilin-2 | 2,085 | 43.0 | +0.03/-0.02 | 0.0 | MODERATE |
| FUS_H | Hendra virus F protein | 245 | 29.6 | -0.60/+0.39 | 0.0 | MILD |
| RfaH | Transcriptional activator | 1,220 | 37.1 | -0.08 | 0.0 | MILD |
| Cas9 | CRISPR endonuclease | 200 | 0.0 | +0.71/+0.44 | 99.5 | BOTH |
| CRKL | Crk-like protein | 260 | 0.0 | +0.44 | 95.0 | BOTH |
| IscA | Fe-S scaffold protein | 1,680 | 0.9 | +0.88 | 86.1 | BOTH |
| OxyR | Peroxide sensor | 2,810 | 0.2 | +0.89/+0.90 | 34.0 | BOTH |
| CaBP* | Calcium-binding protein | 2,385 | 51.3 | -0.30/-0.26 | 43.9 | BOTH* |
| A1AT | Alpha-1 antitrypsin | 2,640 | 1.5 | +0.91/+0.84 | 11.1 | MAINTAINED |
| COMT | Catechol O-methyltransf. | 1,585 | 6.0 | +0.71/+0.64 | 26.4 | MAINTAINED |
| Cwc2 | Splicing factor | 665 | 12.6 | +0.86/+0.86 | 0.0 | MAINTAINED |
| Fab | Neutralizing antibody | 450 | 2.9 | +0.96/+0.95 | 22.8 | MAINTAINED |
| Mad2 | Spindle checkpoint | 1,190 | 0.2 | +0.81/+0.61 | 15.6 | MAINTAINED |
| MinE | Cell division factor | 3,515 | 0.0 | -0.42/-0.08 | 0.0 | MAINTAINED |

### 4.2 The Five-Pattern Taxonomy

The 18 proteins separated cleanly into five confidence integrity categories (Table 1, Figure 1): *SEVERE False Verify* (3 proteins: GP2, KaiB, PimA). FV rates of 88–97%. These proteins show near-total memorization lock: virtually all high-confidence predictions reproduce the dominant fold while completely missing the alternative. KaiB showed 96.4% FV with a negative confidence-accuracy correlation (*R* = −0.115), meaning AF2 is actually *more* confident when *more* wrong about the alternative fold.

**Figure 1:**
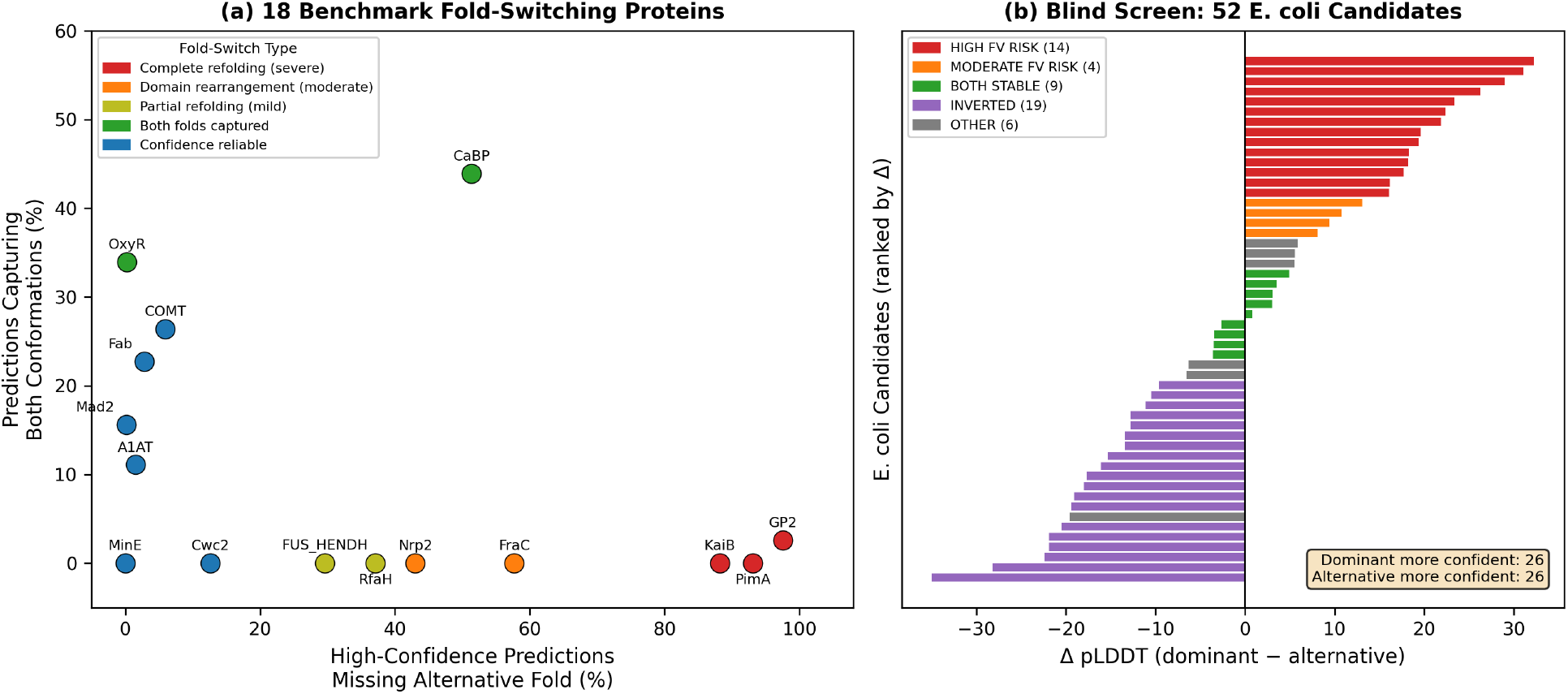
Confidence integrity taxonomy separates fold-switching proteins by failure severity. (a) Benchmark: 18 proteins plotted by high-confidence predictions missing the alternative fold (x-axis) vs. fraction capturing both conformations (y-axis). Colors indicate fold-switch type. Proteins with complete secondary structure refolding (GP2, KaiB, PimA; red) cluster at high FV / low both-folds; proteins with local backbone changes (Mad2, COMT; blue) cluster at low FV / high both-folds. (b) Blind screen: 52 unvalidated *E. coli* candidates ranked by Δ pLDDT. The 50/50 directional split (26 dom > alt, 26 alt > dom) confirms zero population-level correlation between pLDDT and conformational accuracy.

*MODERATE False Verify* (2 proteins: FraC, Nrp2). FV rates of 43–58%. Roughly half of high-confidence predictions are locked into the dominant fold, while the other half show some structural exploration.

*MILD False Verify* (2 proteins: RfaH, FUS_HENDH). FV rates of 30–37%. RfaH shows the largest confidence gap in the dataset: Fold1-only predictions have a mean pLDDT 26.4 points higher than Fold2-only predictions. The model is most confident precisely when it is wrong.

*BOTH FOLDS captured* (5 proteins: Cas9, CRKL, CaBP, IscA, OxyR). These proteins have > 30% of predictions matching both reference folds simultaneously. Cas9 shows 0% FalseVerify with 100% of predictions matching both folds.

*MAINTAINED confidence integrity* (6 proteins: A1AT, COMT, Cwc2, Fab, Mad2, MinE). FV rates below 20%, with positive confidence-accuracy correlations. Mad2 shows 0% FV with *R* = +0.81.

### 4.3 Confidence-Accuracy Correlation as a Diagnostic Metric

The Pearson correlation *R*(pLDDT, TM_fold2_) spanned from *R* = −0.60 (FUS_HENDH) to *R* = +0.96 (Fab) across 33 protein–reference combinations (Figure 2). The weak overall correlation (*R* ≈ 0.52) reported by Bryant and Noé [Bryant and Noé, 2024] masks a bimodal distribution: proteins where confidence tracks accuracy (*R* > +0.6) coexist with proteins where confidence anti-correlates with accuracy (*R <* 0), and no intermediate population exists.

**Figure 2:**
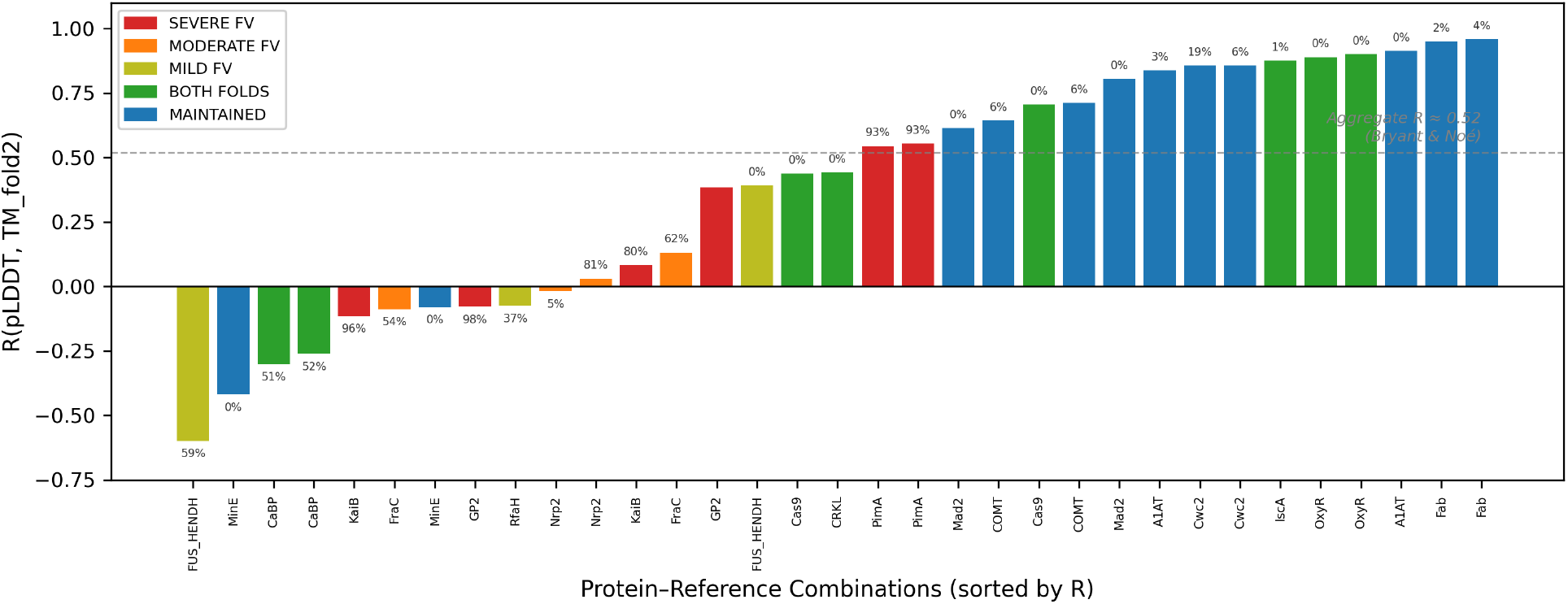
Confidence-accuracy correlation reveals two hidden regimes masked by aggregate statistics. Each bar represents *R*(pLDDT, TM_fold2_) for one of 33 protein-reference combinations, sorted by *R* value. FV rate annotated above each bar. The dashed line marks the aggregate *R* ≈ 0.52 reported by Bryant and Noé [Bryant and Noé, 2024]. SEVERE FV proteins (red) cluster at negative *R*; MAINTAINED proteins (blue) cluster at high positive *R*. No intermediate population exists.

### 4.4 Per-Residue Localization of Confidence Failures

Per-residue analysis revealed that confidence failures are structurally localized to specific fold-switching regions (Figure 3, Table 2). KaiB and OxyR have nearly identical mean pLDDT values (72.6 vs. 76.0), yet KaiB’s cross-cluster variance is 4× higher (74.1 vs. 18.2), primarily concentrated in residues 51–66, precisely the region undergoing the *α* → *β* secondary structure transition. This metric, cross-cluster pLDDT variance, functions as a structural fingerprint of FalseVerify, detectable without any reference to experimental ground truth.

**Figure 3:**
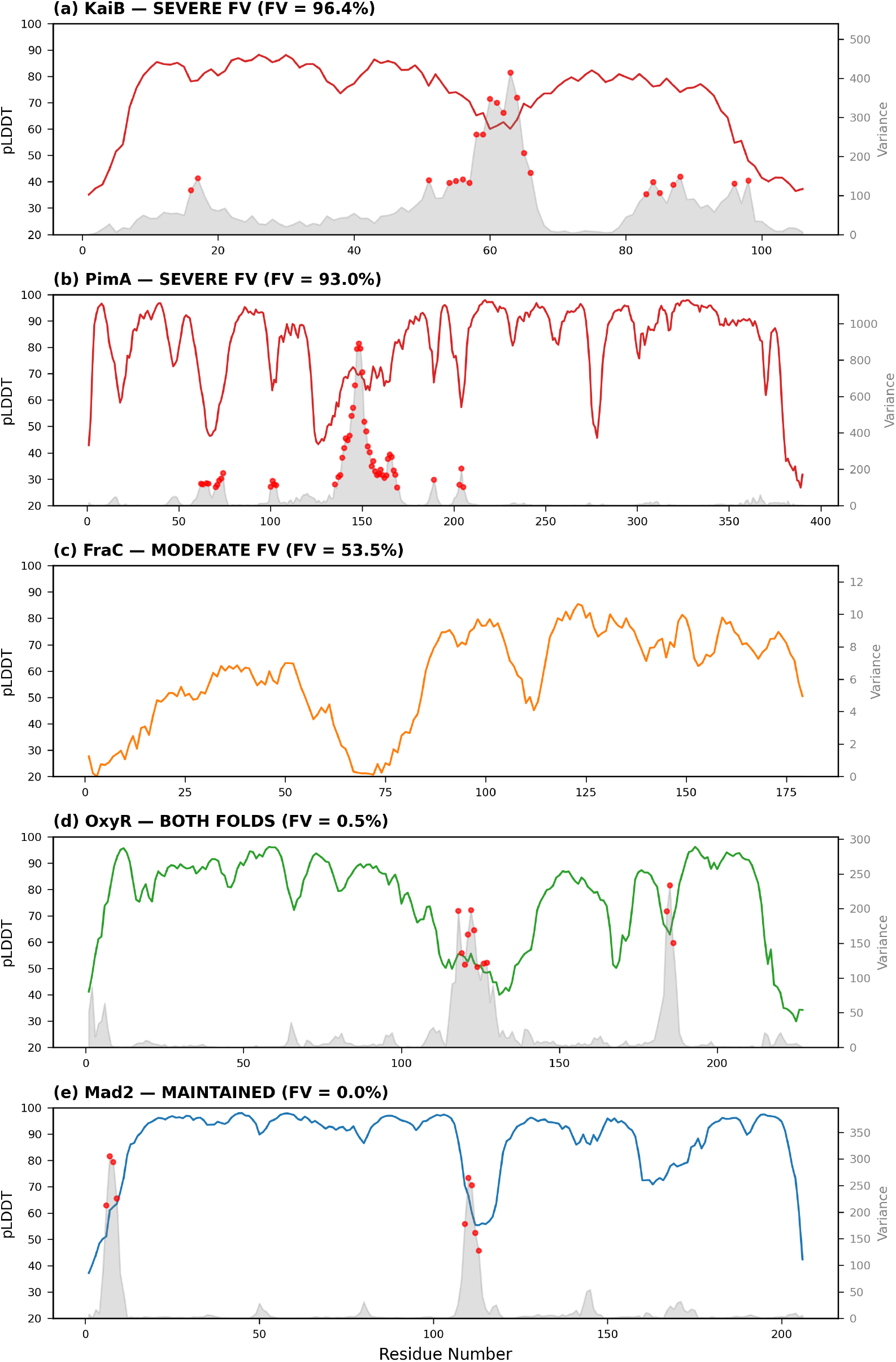
Cross-cluster pLDDT variance localizes confidence breakdown to fold-switching residues. Five panels showing per-residue pLDDT (colored line, left axis) with cross-cluster variance (gray fill, right axis). Red dots mark high-variance residues (>100). (a) KaiB (SEVERE, 96.4% FV): variance peaks at residues 51-66, the *α* → *β* transition. (b) PimA (SEVERE, 93.0% FV): peak variance 891.6 at the domain interface. (c) FraC (MODERATE, 53.5% FV): single cluster, rank-1 vs. rank-5 delta shown. (d) OxyR (BOTH FOLDS, 0.5% FV): uniformly low variance. (e) Mad2 (MAINTAINED, 0.0% FV): high confidence, low variance, indicating well-calibrated confidence.

**Table 2:** Per-residue confidence integrity by category. *FraC: only one cluster available; within-cluster rank delta used as substitute metric.

| Protein | Category | FV% | Mean pLDDT | Variance | %HV | High-Var Region |
| --- | --- | --- | --- | --- | --- | --- |
| KaiB | SEVERE | 96.4 | 72.6 | 74.1 | 22% | res 51-66 ($\alpha \rightarrow \beta$) |
| PimA | SEVERE | 93.0 | 81.1 | 46.4 | 13% | res 135-169 |
| FraC* | MODERATE | 53.5 | 57.8 | n/a | n/a | res 118-137 (rank $\Delta$ +55) |
| OxyR | BOTH | 0.5 | 76.0 | 18.2 | 5% | res 118-127 (minor) |
| Mad2 | MAINTAINED | 0.0 | 87.7 | 16.5 | 4% | res 6-9, 109-113 |

### 4.5 Blind Screening of 52 *E. coli* Candidates

Application of the confidence integrity taxonomy to 52 unvalidated *E. coli* candidates produced a structured category distribution (Table 3, Figure 4): HIGH FV RISK (14, 27%), MODERATE FV RISK (4, 8%), BOTH STABLE (9, 17%), INVERTED (19, 37%), and OTHER (6, 12%). Two population-level results establish generalization. First, mean pLDDT across dominant conformations (78.8) was statistically indistinguishable from alternative conformations (78.7). Second, the directional split was exactly 50/50: 26 proteins had dom_pLDDT > alt_pLDDT and 26 had alt_pLDDT > dom_pLDDT. At the population level, pLDDT carries zero information about conformational accuracy.

**Table 3:** Blind screening of 52 *E. coli* fold-switching candidates. Population split: 26 dom > alt (50%), 26 alt > dom (50%). Mean absolute gap |Δ| = 14.8.

| Category | $N$ | % | Dom pLDDT | Alt pLDDT | Mean $\Delta$ | Benchmark Equiv. |
| --- | --- | --- | --- | --- | --- | --- |
| HIGH FV RISK | 14 | 27 | 90.6 | 67.4 | +23.2 | KaiB, GP2 |
| MODERATE FV RISK | 4 | 8 | 85.2 | 73.5 | +11.7 | RfaH |
| BOTH STABLE | 9 | 17 | 77.5 | 78.0 | -0.5 | Mad2, OxyR |
| INVERTED | 19 | 37 | 71.9 | 89.4 | -17.5 | Novel |
| OTHER | 6 | 12 | 65.7 | 64.8 | +0.9 | Mixed |
| **TOTAL** | **52** |  | **78.8** | **78.7** | **+0.1** |  |

**Figure 4:**
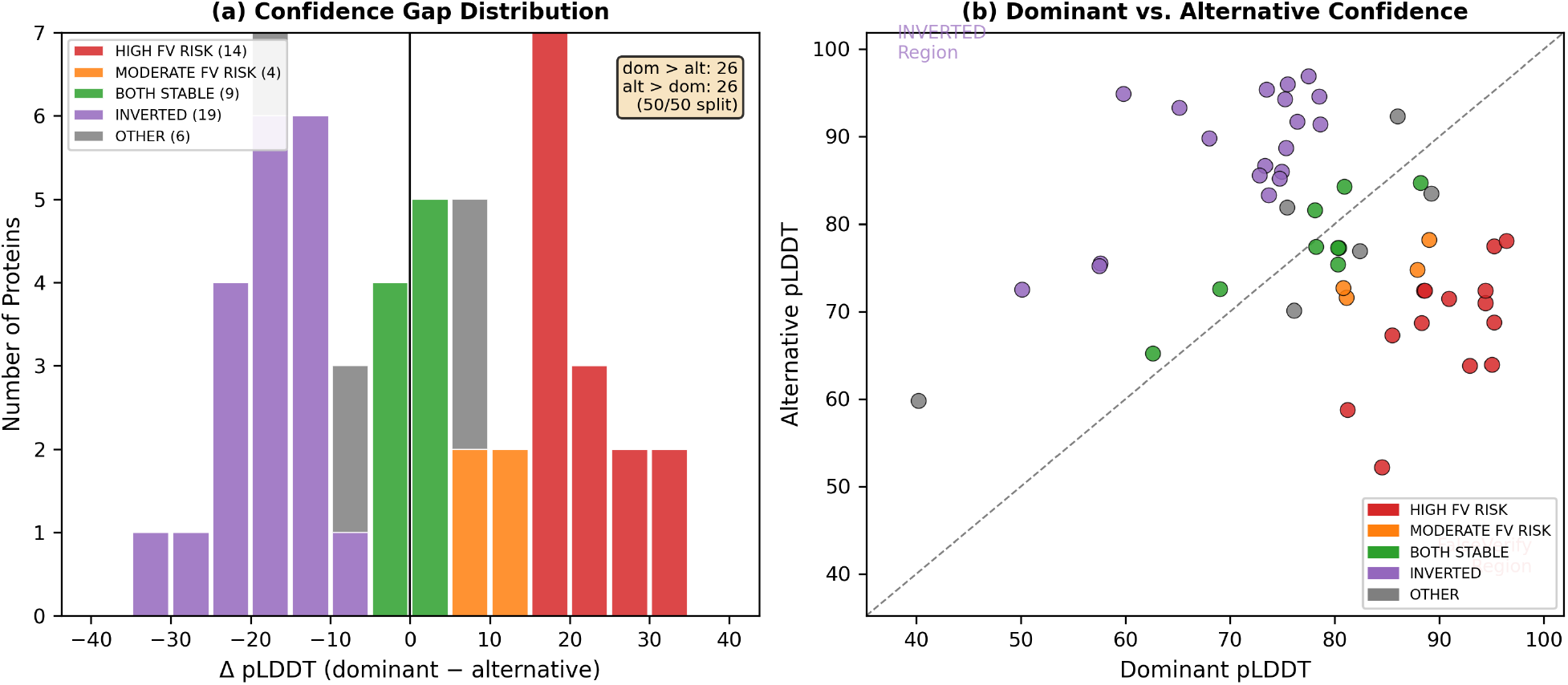
Blind screening of 52 *E. coli* candidates produces structured categories, not noise. (a) Stacked histogram of Δ pLDDT colored by category. The distribution is bimodal with HIGH FV RISK (red) at Δ ≈ +20 and INVERTED (purple) at Δ ≈ −18, with BOTH STABLE (green) near zero. Category counts match Table 3 (14/4/9/19/6). (b) Dominant vs. alternative pLDDT scatter. Points above the diagonal are INVERTED; points below are FalseVerify candidates. The spatial separation confirms the taxonomy captures structural signal from uncharacterized proteins.

**Figure 5:**
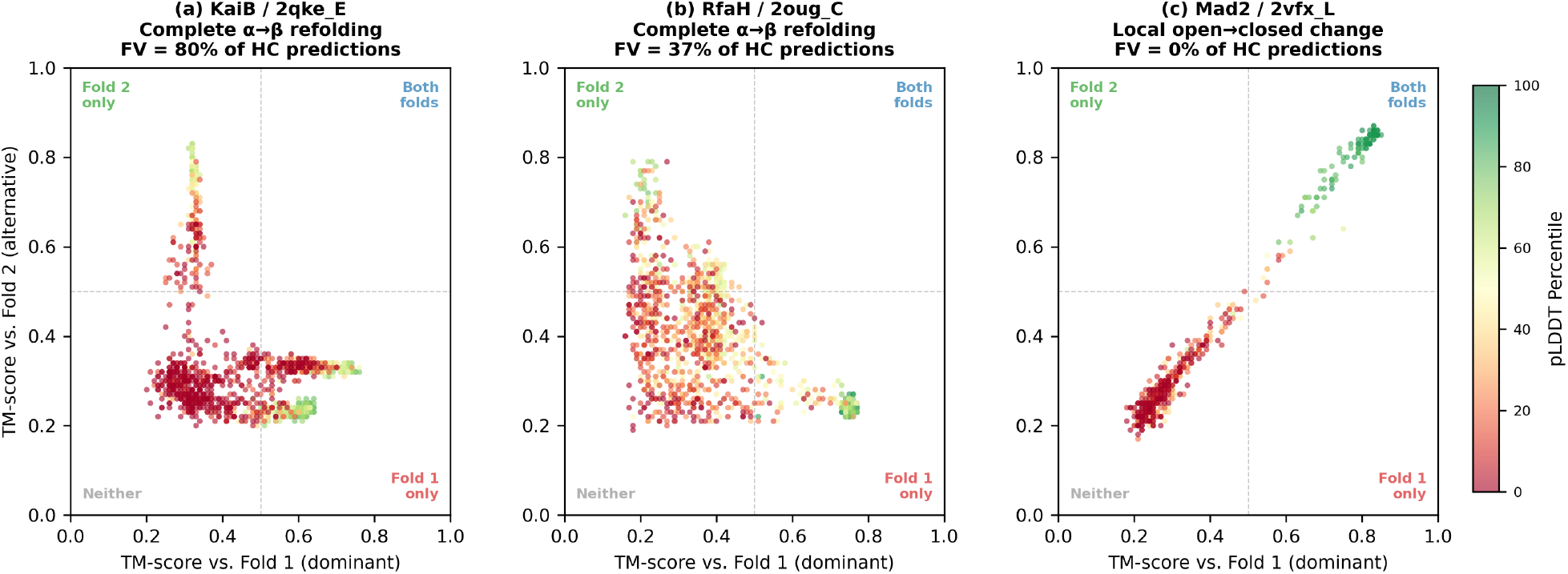
TM-score distributions reveal FalseVerify mechanism in structural biology terms. Each panel plots TM-score vs. fold 1 (x-axis) against TM-score vs. fold 2 (y-axis), colored by pLDDT percentile (green = high confidence). Dashed lines mark TM = 0.5. (a) KaiB: high-confidence predictions (green) are trapped in the Fold 1 only quadrant, indicating confident but wrong predictions. (b) RfaH: intermediate case with some high-confidence escape to Fold 2. (c) Mad2: high-confidence predictions track to the Both folds quadrant, where confidence works as intended.

### 4.6 Discovery of the Inverted Confidence Pattern

The largest single category, 19 of 52 proteins (37%), exhibited a pattern not observed in any benchmark protein: the alternative conformation was predicted with *higher* confidence than the dominant. The most extreme case (WP_001296901) showed dom_pLDDT = 59.8 but alt_pLDDT = 94.9 (Δ = −35.0). This INVERTED pattern functions as an automatic quality flag for the CF-random pipeline, identifying cases where the Dominant/Alternative assignment should be re-examined.

### 4.7 Threshold Sensitivity Analysis

The taxonomy ordering (SEVERE > MODERATE > MILD) holds at both TM ≥ 0.5 and Porter’s own TM ≥ 0.6 success criterion (population FV: 33.6% → 22.9%; Table 4). At stricter thresholds, proteins with marginal fold2 matches (IscA, OxyR, Fab) show *increasing* FV rates, revealing threshold-sensitive confidence-accuracy decoupling invisible at the standard cutoff. GP2 retains 65.0% FV even at TM ≥ 0.8: its fold1-locked predictions are structurally committed to the wrong conformation.

**Table 4:** FalseVerify rate (HC) at multiple TM-score thresholds. Taxonomy ordering holds at 0.5 and 0.6. Proteins with increasing FV at stricter thresholds have marginal fold2 TM-scores.

| Protein | Pattern | FV% @0.5 | FV% @0.6 | FV% @0.7 | FV% @0.8 |
| --- | --- | --- | --- | --- | --- |
| GP2 | SEVERE | 97.5 | 92.5 | 92.5 | 65.0 |
| PimA | SEVERE | 93.1 | 79.3 | 22.6 | 0.0 |
| KaiB | SEVERE | 88.2 | 74.0 | 30.3 | 2.1 |
| FraC | MODERATE | 57.7 | 52.6 | 3.6 | 0.0 |
| Nrp2 | MODERATE | 43.0 | 36.8 | 28.4 | 15.2 |
| RfaH | MILD | 37.1 | 32.8 | 30.4 | 0.0 |
| FUS_H | MILD | 29.6 | 25.5 | 22.4 | 16.3 |
| CaBP | BOTH | 51.3 | 0.0 | 0.0 | 0.0 |
| IscA | BOTH | 0.9 | 27.4 | 92.6 | 40.0 |
| OxyR | BOTH | 0.2 | 0.1 | 3.3 | 28.1 |
| CRKL | BOTH | 0.0 | 0.0 | 0.0 | 0.0 |
| Cas9 | BOTH | 0.0 | 0.0 | 0.0 | 0.0 |
| Cwc2 | MAINT. | 12.6 | 0.0 | 0.0 | 0.0 |
| COMT | MAINT. | 6.0 | 1.5 | 0.0 | 0.0 |
| Fab | MAINT. | 2.9 | 3.8 | 6.7 | 16.2 |
| A1AT | MAINT. | 1.5 | 0.0 | 0.0 | 0.0 |
| Mad2 | MAINT. | 0.2 | 0.0 | 0.4 | 0.4 |
| MinE | MAINT. | 0.0 | 0.0 | 0.0 | 0.0 |
| **Population** |  | **33.6** | **22.9** | **18.6** | **8.0** |

## 5 Discussion

### 5.1 Confidence-Accuracy Decoupling Is Systematic

The 33.6% population-level FalseVerify rate establishes that pLDDT unreliability for fold-switching proteins is not anecdotal but systematic. One in three high-confidence AF2 predictions for known fold-switching proteins is confidently wrong about the alternative conformation. The blind *E. coli* screening extends this: the perfect 50/50 directional split and identical population means demonstrate that pLDDT carries zero information about conformational accuracy at the proteome scale.

The five-pattern taxonomy maps to structural biology: the magnitude of the conformational change predicts the severity of the confidence failure. Complete *α* ↔ *β* refolding produces SEVERE FV; local backbone changes produce none. The discovery of the INVERTED pattern in blind data demonstrates that the taxonomy is extensible rather than closed.

### 5.2 From Description to Prediction: The *E. coli* Blind Test

Three results support the taxonomy’s predictive utility. First, the category distribution is structured (27%/8%/17%/37%/12%), not uniform. Second, benchmark-derived thresholds produce interpretable classifications on novel proteins. Third, the INVERTED category discovery demonstrates sensitivity to biological signal rather than overfitting.

The practical implication is immediate: our screening reduces the experimental priority list from 52 candidates to 9 BOTH STABLE proteins with the highest confidence integrity, while flagging 14 HIGH FV candidates and 19 INVERTED candidates for re-examination.

### 5.3 Cross-Cluster Variance as a Model-Free Diagnostic

Cross-cluster pLDDT variance requires no experimental ground truth and provides a per-residue reliability map identifying specific regions where AF2’s confidence should not be trusted. The comparison between KaiB (variance 74.1, 96% FV) and OxyR (variance 18.2, 0.5% FV) at similar mean pLDDT (72.6 vs. 76.0) demonstrates the metric’s discriminatory power.

### 5.4 Relationship to Existing Work

Our results extend and quantify observations from several groups. Porter and colleagues identified structure memorization as the driver of AF2’s fold-switching failures [Chakravarty et al., 2024, Schafer and Porter, 2025]; we quantify the confidence signature of this memorization. Bryant and Noé reported *R* = 0.52 between pLDDT and TM-score for alternative conformations [Bryant and Noé, 2024]; our finding of *R* ranging from −0.60 to +0.96 refines this: the weak overall correlation masks a bimodal distribution. Schafer and Porter called for “alternative measures” to replace pLDDT [Schafer and Porter, 2025]; the FalseVerify rate and cross-cluster pLDDT variance jointly provide such measures.

### 5.5 Relationship to the Full Porter Benchmark

Independent analysis of the full 92-protein benchmark yielded a 40.4% FalseVerify rate (85, 643/212,018 predictions). The present analysis reports 33.6% across the 18-protein subset. The convergence of both analyses on FalseVerify rates in the 33-40% range, despite independent methodologies, strengthens the conclusion that confidence-accuracy decoupling is systematic.

### 5.6 Limitations

Several limitations should be noted. First, the *E. coli* candidates lack experimental ground truth; the confidence integrity categories are predictions, not validated outcomes. Second, our analysis uses pLDDT percentile ranks rather than raw scores. Third, per-residue analysis was performed on five proteins with limited clusters. Fourth, we have not tested on AlphaFold predictions. Fifth, the INVERTED pattern interpretation cannot distinguish label inversion from genuine conformational complexity without additional experimental data.

## 6 Conclusion

We present the first systematic confidence integrity screen for fold-switching protein predictions, validated across 27,098 predictions from 18 experimentally characterized proteins and applied blindly to 52 unvalidated *E. coli* candidates.

The key findings are:

1. One in three high-confidence AF2 predictions for known fold-switching proteins is FalseVerify, simultaneously confident and wrong (33.6% HC FV rate).
2. FalseVerify severity is predictable from fold-switch magnitude: complete secondary structure changes produce 80-97% FV rates; local rearrangements produce 0-2%.
3. Pearson *R*(pLDDT, TM_alt_) ranges from −0.60 to +0.96, discriminating reliable from unreliable predictions.
4. Cross-cluster pLDDT variance provides a per-residue diagnostic requiring no experimental ground truth.
5. Blind application to 52 *E. coli* candidates produces structured categories with a 50/50 directional split confirming zero correlation between pLDDT and conformational accuracy.
6. A sixth category, INVERTED confidence, emerged from blind screening and was invisible in benchmark data.
7. The taxonomy ordering is robust to threshold choice: SEVERE > MODERATE > MILD holds at TM ≥ 0.5 and TM ≥ 0.6.

These results fill the specific methodological gap identified by Schafer and Porter Schafer and Porter, 2025]. The framework and all analysis scripts are publicly available; experimental validation of the *E. coli* predictions is invited.

## Use of AI Tools

This research was conducted with the assistance of AI language models (Claude, Anthropic) for data analysis, figure generation, manuscript drafting, and iterative verification of quantitative claims. All findings, interpretations, and scientific conclusions are the responsibility of the author. Every quantitative claim was independently verified against source data.

## Data and Code Availability

Porter fold-switching benchmark data: Chakravarty et al. [2024], available at https://github.com/ncbi/AF2_benchmark (Zenodo doi:10.5281/zenodo.13221958).

CF-random software and *E. coli* predictions: Lee et al. [2025], available at https://github.com/ncbi/CF-random_software.

Analysis scripts and per-residue figure data will be deposited at the preprint repository upon publication.

