## supplement_v6 for "Confidence Without Verification: Screening pLDDT Unreliability in AlphaFold2 Fold-Switching Predictions": AlphaFold2.ipynb

```
# %% [markdown]
<img src="https://raw.githubusercontent.com/sokrypton/ColabFold/main/.github/ColabFold_Marv_Logo_Small.png" height="200" align="right" style="height:240px">

##ColabFold v1.5.5: AlphaFold2 using MMseqs2

Easy to use protein structure and complex prediction using [AlphaFold2](https://www.nature.com/articles/s41586-021-03819-2) and [Alphafold2-multimer](https://www.biorxiv.org/content/10.1101/2021.10.04.463034v1). Sequence alignments/templates are generated through [MMseqs2](mmseqs.com) and [HHsearch](https://github.com/soedinglab/hh-suite). For more details, see <a href="#Instructions">bottom</a> of the notebook, checkout the [ColabFold GitHub](https://github.com/sokrypton/ColabFold) and [Nature Protocols](https://www.nature.com/articles/s41596-024-01060-5).

Old versions: [v1.4](https://colab.research.google.com/github/sokrypton/ColabFold/blob/v1.4.0/AlphaFold2.ipynb), [v1.5.1](https://colab.research.google.com/github/sokrypton/ColabFold/blob/v1.5.1/AlphaFold2.ipynb), [v1.5.2](https://colab.research.google.com/github/sokrypton/ColabFold/blob/v1.5.2/AlphaFold2.ipynb), [v1.5.3-patch](https://colab.research.google.com/github/sokrypton/ColabFold/blob/56c72044c7d51a311ca99b953a71e552fdc042e1/AlphaFold2.ipynb)

[Mirdita M, Schütze K, Moriwaki Y, Heo L, Ovchinnikov S, Steinegger M. ColabFold: Making protein folding accessible to all.
*Nature Methods*, 2022](https://www.nature.com/articles/s41592-022-01488-1)

# %% [code]
#@title Input protein sequence(s), then hit `Runtime` -> `Run all`
from google.colab import files
import os
import re
import hashlib
import random

from sys import version_info
python_version = f"{version_info.major}.{version_info.minor}"

def add_hash(x,y):
  return x+"_"+hashlib.sha1(y.encode()).hexdigest()[:5]

query_sequence = 'VGSEVSDKRTCVSLTTQRLPVSRIKTYTITEGSLRAVIFITKRGCRVNSGCCIENSLIVFFKQSKSWKRCEKNNNPAPRFKANMVDSFSWGRVHAQTLNS' #@param {type:"string"}
#@markdown  - Use `:` to specify inter-protein chainbreaks for **modeling complexes** (supports homo- and hetro-oligomers). For example **PI...SK:PI...SK** for a homodimer
jobname = 'XCL1_deep' #@param {type:"string"}
# number of models to use
num_relax = 0 #@param [0, 1, 5] {type:"raw"}
#@markdown - specify how many of the top ranked structures to relax using amber
template_mode = "none" #@param ["none", "pdb100","custom"]
#@markdown - `none` = no template information is used. `pdb100` = detect templates in pdb100 (see [notes](#pdb100)). `custom` - upload and search own templates (PDB or mmCIF format, see [notes](#custom_templates))

use_amber = num_relax > 0

# remove whitespaces
query_sequence = "".join(query_sequence.split())

basejobname = "".join(jobname.split())
basejobname = re.sub(r'\W+', '', basejobname)
jobname = add_hash(basejobname, query_sequence)

# check if directory with jobname exists
def check(folder):
  if os.path.exists(folder):
    return False
  else:
    return True
if not check(jobname):
  n = 0
  while not check(f"{jobname}_{n}"): n += 1
  jobname = f"{jobname}_{n}"

# make directory to save results
os.makedirs(jobname, exist_ok=True)

# save queries
queries_path = os.path.join(jobname, f"{jobname}.csv")
with open(queries_path, "w") as text_file:
  text_file.write(f"id,sequence\n{jobname},{query_sequence}")

if template_mode == "pdb100":
  use_templates = True
  custom_template_path = None
elif template_mode == "custom":
  custom_template_path = os.path.join(jobname,f"template")
  os.makedirs(custom_template_path, exist_ok=True)
  uploaded = files.upload()
  use_templates = True
  for fn in uploaded.keys():
    os.rename(fn,os.path.join(custom_template_path,fn))
else:
  custom_template_path = None
  use_templates = False

print("jobname",jobname)
print("sequence",query_sequence)
print("length",len(query_sequence.replace(":","")))

# %% [code]
#@title Install dependencies
%%time
import os
USE_AMBER = use_amber
USE_TEMPLATES = use_templates
PYTHON_VERSION = python_version

if not os.path.isfile("COLABFOLD_READY"):
  print("installing colabfold...")
  os.system("pip install -q --no-warn-conflicts 'colabfold[alphafold-minus-jax] @ git+https://github.com/sokrypton/ColabFold'")
  if os.environ.get('TPU_NAME', False) != False:
    os.system("pip uninstall -y jax jaxlib")
    os.system("pip install --no-warn-conflicts --upgrade dm-haiku==0.0.10 'jax[cuda12_pip]'==0.3.25 -f https://storage.googleapis.com/jax-releases/jax_cuda_releases.html")
  os.system("ln -s /usr/local/lib/python3.*/dist-packages/colabfold colabfold")
  os.system("ln -s /usr/local/lib/python3.*/dist-packages/alphafold alphafold")
  # hack to fix TF crash
  os.system("rm -f /usr/local/lib/python3.*/dist-packages/tensorflow/core/kernels/libtfkernel_sobol_op.so")
  os.system("touch COLABFOLD_READY")

if USE_AMBER or USE_TEMPLATES:
  if not os.path.isfile("CONDA_READY"):
    print("installing conda...")
    os.system("wget -qnc https://github.com/conda-forge/miniforge/releases/latest/download/Miniforge3-Linux-x86_64.sh")
    os.system("bash Miniforge3-Linux-x86_64.sh -bfp /usr/local")
    os.system("mamba config --set auto_update_conda false")
    os.system("touch CONDA_READY")

if USE_TEMPLATES and not os.path.isfile("HH_READY") and USE_AMBER and not os.path.isfile("AMBER_READY"):
  print("installing hhsuite and amber...")
  os.system(f"mamba install -y -c conda-forge -c bioconda kalign2=2.04 hhsuite=3.3.0 openmm=8.2.0 python='{PYTHON_VERSION}' pdbfixer")
  os.system("touch HH_READY")
  os.system("touch AMBER_READY")
else:
  if USE_TEMPLATES and not os.path.isfile("HH_READY"):
    print("installing hhsuite...")
    os.system(f"mamba install -y -c conda-forge -c bioconda kalign2=2.04 hhsuite=3.3.0 python='{PYTHON_VERSION}'")
    os.system("touch HH_READY")
  if USE_AMBER and not os.path.isfile("AMBER_READY"):
    print("installing amber...")
    os.system(f"mamba install -y -c conda-forge openmm=8.2.0 python='{PYTHON_VERSION}' pdbfixer")
    os.system("touch AMBER_READY")

# %% [code]
#@markdown ### MSA options (custom MSA upload, single sequence, pairing mode)
msa_mode = "mmseqs2_uniref_env" #@param ["mmseqs2_uniref_env", "mmseqs2_uniref","single_sequence","custom"]
pair_mode = "unpaired_paired" #@param ["unpaired_paired","paired","unpaired"] {type:"string"}
#@markdown - "unpaired_paired" = pair sequences from same species + unpaired MSA, "unpaired" = seperate MSA for each chain, "paired" - only use paired sequences.

# decide which a3m to use
if "mmseqs2" in msa_mode:
  a3m_file = os.path.join(jobname,f"{jobname}.a3m")

elif msa_mode == "custom":
  a3m_file = os.path.join(jobname,f"{jobname}.custom.a3m")
  if not os.path.isfile(a3m_file):
    custom_msa_dict = files.upload()
    custom_msa = list(custom_msa_dict.keys())[0]
    header = 0
    import fileinput
    for line in fileinput.FileInput(custom_msa,inplace=1):
      if line.startswith(">"):
         header = header + 1
      if not line.rstrip():
        continue
      if line.startswith(">") == False and header == 1:
         query_sequence = line.rstrip()
      print(line, end='')

    os.rename(custom_msa, a3m_file)
    queries_path=a3m_file
    print(f"moving {custom_msa} to {a3m_file}")

else:
  a3m_file = os.path.join(jobname,f"{jobname}.single_sequence.a3m")
  with open(a3m_file, "w") as text_file:
    text_file.write(">1\n%s" % query_sequence)

# %% [code]
#@markdown ### Advanced settings
model_type = "auto" #@param ["auto", "alphafold2_ptm", "alphafold2_multimer_v1", "alphafold2_multimer_v2", "alphafold2_multimer_v3", "deepfold_v1", "alphafold2"]
#@markdown - if `auto` selected, will use `alphafold2_ptm` for monomer prediction and `alphafold2_multimer_v3` for complex prediction.
#@markdown Any of the mode_types can be used (regardless if input is monomer or complex).
num_recycles = "3" #@param ["auto", "0", "1", "3", "6", "12", "24", "48"]
#@markdown - if `auto` selected, will use `num_recycles=20` if `model_type=alphafold2_multimer_v3`, else `num_recycles=3` .
recycle_early_stop_tolerance = "auto" #@param ["auto", "0.0", "0.5", "1.0"]
#@markdown - if `auto` selected, will use `tol=0.5` if `model_type=alphafold2_multimer_v3` else `tol=0.0`.
relax_max_iterations = 200 #@param [0, 200, 2000] {type:"raw"}
#@markdown - max amber relax iterations, `0` = unlimited (AlphaFold2 default, can take very long)
pairing_strategy = "greedy" #@param ["greedy", "complete"] {type:"string"}
#@markdown - `greedy` = pair any taxonomically matching subsets, `complete` = all sequences have to match in one line.
calc_extra_ptm = False #@param {type:"boolean"}
#@markdown - return pairwise chain iptm/actifptm

#@markdown #### Sample settings
#@markdown -  enable dropouts and increase number of seeds to sample predictions from uncertainty of the model.
#@markdown -  decrease `max_msa` to increase uncertainity
max_msa = "auto" #@param ["auto", "512:1024", "256:512", "64:128", "32:64", "16:32"]
num_seeds = 1 #@param [1,2,4,8,16] {type:"raw"}
use_dropout = False #@param {type:"boolean"}

num_recycles = None if num_recycles == "auto" else int(num_recycles)
recycle_early_stop_tolerance = None if recycle_early_stop_tolerance == "auto" else float(recycle_early_stop_tolerance)
if max_msa == "auto": max_msa = None

#@markdown #### Save settings
save_all = False #@param {type:"boolean"}
save_recycles = False #@param {type:"boolean"}
save_to_google_drive = False #@param {type:"boolean"}
#@markdown -  if the save_to_google_drive option was selected, the result zip will be uploaded to your Google Drive
dpi = 200 #@param {type:"integer"}
#@markdown - set dpi for image resolution

if save_to_google_drive:
  from pydrive2.drive import GoogleDrive
  from pydrive2.auth import GoogleAuth
  from google.colab import auth
  from oauth2client.client import GoogleCredentials
  auth.authenticate_user()
  gauth = GoogleAuth()
  gauth.credentials = GoogleCredentials.get_application_default()
  drive = GoogleDrive(gauth)
  print("You are logged into Google Drive and are good to go!")

#@markdown Don't forget to hit `Runtime` -> `Run all` after updating the form.

# %% [code]
#@title Run Prediction
display_images = True #@param {type:"boolean"}

import sys
import warnings
warnings.simplefilter(action='ignore', category=FutureWarning)
from Bio import BiopythonDeprecationWarning
warnings.simplefilter(action='ignore', category=BiopythonDeprecationWarning)
from pathlib import Path
from colabfold.download import download_alphafold_params, default_data_dir
from colabfold.utils import setup_logging
from colabfold.batch import get_queries, run, set_model_type
from colabfold.plot import plot_msa_v2

import os
import numpy as np
try:
  K80_chk = os.popen('nvidia-smi | grep "Tesla K80" | wc -l').read()
except:
  K80_chk = "0"
  pass
if "1" in K80_chk:
  print("WARNING: found GPU Tesla K80: limited to total length < 1000")
  if "TF_FORCE_UNIFIED_MEMORY" in os.environ:
    del os.environ["TF_FORCE_UNIFIED_MEMORY"]
  if "XLA_PYTHON_CLIENT_MEM_FRACTION" in os.environ:
    del os.environ["XLA_PYTHON_CLIENT_MEM_FRACTION"]

from colabfold.colabfold import plot_protein
from pathlib import Path
import matplotlib.pyplot as plt

# For some reason we need that to get pdbfixer to import
if use_amber and f"/usr/local/lib/python{python_version}/site-packages/" not in sys.path:
    sys.path.insert(0, f"/usr/local/lib/python{python_version}/site-packages/")

def input_features_callback(input_features):
  if display_images:
    plot_msa_v2(input_features)
    plt.show()
    plt.close()

def prediction_callback(protein_obj, length,
                        prediction_result, input_features, mode):
  model_name, relaxed = mode
  if not relaxed:
    if display_images:
      fig = plot_protein(protein_obj, Ls=length, dpi=150)
      plt.show()
      plt.close()

result_dir = jobname
log_filename = os.path.join(jobname,"log.txt")
setup_logging(Path(log_filename))

queries, is_complex = get_queries(queries_path)
model_type = set_model_type(is_complex, model_type)

if "multimer" in model_type and max_msa is not None:
  use_cluster_profile = False
else:
  use_cluster_profile = True

download_alphafold_params(model_type, Path("."))
results = run(
    queries=queries,
    result_dir=result_dir,
    use_templates=use_templates,
    custom_template_path=custom_template_path,
    num_relax=num_relax,
    msa_mode=msa_mode,
    model_type=model_type,
    num_models=5,
    num_recycles=num_recycles,
    relax_max_iterations=relax_max_iterations,
    recycle_early_stop_tolerance=recycle_early_stop_tolerance,
    num_seeds=num_seeds,
    use_dropout=use_dropout,
    model_order=[1,2,3,4,5],
    is_complex=is_complex,
    data_dir=Path("."),
    keep_existing_results=False,
    rank_by="auto",
    pair_mode=pair_mode,
    pairing_strategy=pairing_strategy,
    stop_at_score=float(100),
    prediction_callback=prediction_callback,
    dpi=dpi,
    zip_results=False,
    save_all=save_all,
    max_msa=max_msa,
    use_cluster_profile=use_cluster_profile,
    input_features_callback=input_features_callback,
    save_recycles=save_recycles,
    user_agent="colabfold/google-colab-main",
    calc_extra_ptm=calc_extra_ptm,
)
results_zip = f"{jobname}.result.zip"
os.system(f"zip -r {results_zip} {jobname}")

# %% [code]
#@title Display 3D structure {run: "auto"}
import py3Dmol
import glob
import matplotlib.pyplot as plt
from colabfold.colabfold import plot_plddt_legend
from colabfold.colabfold import pymol_color_list, alphabet_list
rank_num = 1 #@param ["1", "2", "3", "4", "5"] {type:"raw"}
color = "lDDT" #@param ["chain", "lDDT", "rainbow"]
show_sidechains = False #@param {type:"boolean"}
show_mainchains = False #@param {type:"boolean"}

tag = results["rank"][0][rank_num - 1]
jobname_prefix = ".custom" if msa_mode == "custom" else ""
pdb_filename = f"{jobname}/{jobname}{jobname_prefix}_unrelaxed_{tag}.pdb"
pdb_file = glob.glob(pdb_filename)

def show_pdb(rank_num=1, show_sidechains=False, show_mainchains=False, color="lDDT"):
  model_name = f"rank_{rank_num}"
  view = py3Dmol.view(js='https://3dmol.org/build/3Dmol.js',)
  view.addModel(open(pdb_file[0],'r').read(),'pdb')

  if color == "lDDT":
    view.setStyle({'cartoon': {'colorscheme': {'prop':'b','gradient': 'roygb','min':50,'max':90}}})
  elif color == "rainbow":
    view.setStyle({'cartoon': {'color':'spectrum'}})
  elif color == "chain":
    chains = len(queries[0][1]) + 1 if is_complex else 1
    for n,chain,color in zip(range(chains),alphabet_list,pymol_color_list):
       view.setStyle({'chain':chain},{'cartoon': {'color':color}})

  if show_sidechains:
    BB = ['C','O','N']
    view.addStyle({'and':[{'resn':["GLY","PRO"],'invert':True},{'atom':BB,'invert':True}]},
                        {'stick':{'colorscheme':f"WhiteCarbon",'radius':0.3}})
    view.addStyle({'and':[{'resn':"GLY"},{'atom':'CA'}]},
                        {'sphere':{'colorscheme':f"WhiteCarbon",'radius':0.3}})
    view.addStyle({'and':[{'resn':"PRO"},{'atom':['C','O'],'invert':True}]},
                        {'stick':{'colorscheme':f"WhiteCarbon",'radius':0.3}})
  if show_mainchains:
    BB = ['C','O','N','CA']
    view.addStyle({'atom':BB},{'stick':{'colorscheme':f"WhiteCarbon",'radius':0.3}})

  view.zoomTo()
  return view

show_pdb(rank_num, show_sidechains, show_mainchains, color).show()
if color == "lDDT":
  plot_plddt_legend().show()

# %% [code]
#@title Plots {run: "auto"}
from IPython.display import display, HTML
import base64
from html import escape

# see: https://stackoverflow.com/a/53688522
def image_to_data_url(filename):
  ext = filename.split('.')[-1]
  prefix = f'data:image/{ext};base64,'
  with open(filename, 'rb') as f:
    img = f.read()
  return prefix + base64.b64encode(img).decode('utf-8')

pae = ""
pae_file = os.path.join(jobname,f"{jobname}{jobname_prefix}_pae.png")
if os.path.isfile(pae_file):
    pae = image_to_data_url(pae_file)
cov = image_to_data_url(os.path.join(jobname,f"{jobname}{jobname_prefix}_coverage.png"))
plddt = image_to_data_url(os.path.join(jobname,f"{jobname}{jobname_prefix}_plddt.png"))
display(HTML(f"""
<style>
  img {{
    float:left;
  }}
  .full {{
    max-width:100%;
  }}
  .half {{
    max-width:50%;
  }}
  @media (max-width:640px) {{
    .half {{
      max-width:100%;
    }}
  }}
</style>
<div style="max-width:90%; padding:2em;">
  <h1>Plots for {escape(jobname)}</h1>
  { '<!--' if pae == '' else '' }<img src="{pae}" class="full" />{ '-->' if pae == '' else '' }
  <img src="{cov}" class="half" />
  <img src="{plddt}" class="half" />
</div>
"""))

# %% [code]
#@title Package and download results
#@markdown If you are having issues downloading the result archive, try disabling your adblocker and run this cell again. If that fails click on the little folder icon to the left, navigate to file: `jobname.result.zip`, right-click and select \"Download\" (see [screenshot](https://pbs.twimg.com/media/E6wRW2lWUAEOuoe?format=jpg&name=small)).

if msa_mode == "custom":
  print("Don't forget to cite your custom MSA generation method.")

files.download(f"{jobname}.result.zip")

if save_to_google_drive == True and drive:
  uploaded = drive.CreateFile({'title': f"{jobname}.result.zip"})
  uploaded.SetContentFile(f"{jobname}.result.zip")
  uploaded.Upload()
  print(f"Uploaded {jobname}.result.zip to Google Drive with ID {uploaded.get('id')}")

# %% [markdown]
# Instructions <a name="Instructions"></a>
For detailed instructions, tips and tricks, see recently published paper at [Nature Protocols](https://www.nature.com/articles/s41596-024-01060-5)

**Quick start**
1. Paste your protein sequence(s) in the input field.
2. Press "Runtime" -> "Run all".
3. The pipeline consists of 5 steps. The currently running step is indicated by a circle with a stop sign next to it.

**Result zip file contents**

1. PDB formatted structures sorted by avg. pLDDT and complexes are sorted by pTMscore. (unrelaxed and relaxed if `use_amber` is enabled).
2. Plots of the model quality.
3. Plots of the MSA coverage.
4. Parameter log file.
5. A3M formatted input MSA.
6. A `predicted_aligned_error_v1.json` using [AlphaFold-DB's format](https://alphafold.ebi.ac.uk/faq#faq-7) and a `scores.json` for each model which contains an array (list of lists) for PAE, a list with the average pLDDT and the pTMscore.
7. BibTeX file with citations for all used tools and databases.

At the end of the job a download modal box will pop up with a `jobname.result.zip` file. Additionally, if the `save_to_google_drive` option was selected, the `jobname.result.zip` will be uploaded to your Google Drive.

**MSA generation for complexes**

For the complex prediction we use unpaired and paired MSAs. Unpaired MSA is generated the same way as for the protein structures prediction by searching the UniRef100 and environmental sequences three iterations each.

The paired MSA is generated by searching the UniRef100 database and pairing the best hits sharing the same NCBI taxonomic identifier (=species or sub-species). We only pair sequences if all of the query sequences are present for the respective taxonomic identifier.

**Using a custom MSA as input**

To predict the structure with a custom MSA (A3M formatted): (1) Change the `msa_mode`: to "custom", (2) Wait for an upload box to appear at the end of the "MSA options ..." box. Upload your A3M. The first fasta entry of the A3M must be the query sequence without gaps.

It is also possilbe to provide custom MSAs for complex predictions. Read more about the format [here](https://github.com/sokrypton/ColabFold/issues/76).

As an alternative for MSA generation the [HHblits Toolkit server](https://toolkit.tuebingen.mpg.de/tools/hhblits) can be used. After submitting your query, click "Query Template MSA" -> "Download Full A3M". Download the A3M file and upload it in this notebook.

**PDB100** <a name="pdb100"></a>

As of 23/06/08, we have transitioned from using the PDB70 to a 100% clustered PDB, the PDB100. The construction methodology of PDB100 differs from that of PDB70.

The PDB70 was constructed by running each PDB70 representative sequence through [HHblits](https://github.com/soedinglab/hh-suite) against the [Uniclust30](https://uniclust.mmseqs.com/). On the other hand, the PDB100 is built by searching each PDB100 representative structure with [Foldseek](https://github.com/steineggerlab/foldseek) against the [AlphaFold Database](https://alphafold.ebi.ac.uk).

To maintain compatibility with older Notebook versions and local installations, the generated files and API responses will continue to be named "PDB70", even though we're now using the PDB100.

**Using custom templates** <a name="custom_templates"></a>

To predict the structure with a custom template (PDB or mmCIF formatted): (1) change the `template_mode` to "custom" in the execute cell and (2) wait for an upload box to appear at the end of the "Input Protein" box. Select and upload your templates (multiple choices are possible).

* Templates must follow the four letter PDB naming with lower case letters.

* Templates in mmCIF format must contain `_entity_poly_seq`. An error is thrown if this field is not present. The field `_pdbx_audit_revision_history.revision_date` is automatically generated if it is not present.

* Templates in PDB format are automatically converted to the mmCIF format. `_entity_poly_seq` and `_pdbx_audit_revision_history.revision_date` are automatically generated.

If you encounter problems, please report them to this [issue](https://github.com/sokrypton/ColabFold/issues/177).

**Comparison to the full AlphaFold2 and AlphaFold2 Colab**

This notebook replaces the homology detection and MSA pairing of AlphaFold2 with MMseqs2. For a comparison against the [AlphaFold2 Colab](https://colab.research.google.com/github/deepmind/alphafold/blob/main/notebooks/AlphaFold.ipynb) and the full [AlphaFold2](https://github.com/deepmind/alphafold) system read our [paper](https://www.nature.com/articles/s41592-022-01488-1).

**Troubleshooting**
* Check that the runtime type is set to GPU at "Runtime" -> "Change runtime type".
* Try to restart the session "Runtime" -> "Factory reset runtime".
* Check your input sequence.

**Known issues**
* Google Colab assigns different types of GPUs with varying amount of memory. Some might not have enough memory to predict the structure for a long sequence.
* Your browser can block the pop-up for downloading the result file. You can choose the `save_to_google_drive` option to upload to Google Drive instead or manually download the result file: Click on the little folder icon to the left, navigate to file: `jobname.result.zip`, right-click and select \"Download\" (see [screenshot](https://pbs.twimg.com/media/E6wRW2lWUAEOuoe?format=jpg&name=small)).

**Limitations**
* Computing resources: Our MMseqs2 API can handle ~20-50k requests per day.
* MSAs: MMseqs2 is very precise and sensitive but might find less hits compared to HHblits/HMMer searched against BFD or MGnify.
* We recommend to additionally use the full [AlphaFold2 pipeline](https://github.com/deepmind/alphafold).

**Description of the plots**
*   **Number of sequences per position** - We want to see at least 30 sequences per position, for best performance, ideally 100 sequences.
*   **Predicted lDDT per position** - model confidence (out of 100) at each position. The higher the better.
*   **Predicted Alignment Error** - For homooligomers, this could be a useful metric to assess how confident the model is about the interface. The lower the better.

**Bugs**
- If you encounter any bugs, please report the issue to https://github.com/sokrypton/ColabFold/issues

**License**

The source code of ColabFold is licensed under [MIT](https://raw.githubusercontent.com/sokrypton/ColabFold/main/LICENSE). Additionally, this notebook uses the AlphaFold2 source code and its parameters licensed under [Apache 2.0](https://raw.githubusercontent.com/deepmind/alphafold/main/LICENSE) and [CC BY 4.0](https://creativecommons.org/licenses/by-sa/4.0/) respectively. Read more about the AlphaFold license [here](https://github.com/deepmind/alphafold).

**Acknowledgments**
- We thank the AlphaFold team for developing an excellent model and open sourcing the software.

- [KOBIC](https://kobic.re.kr) and [Söding Lab](https://www.mpinat.mpg.de/soeding) for providing the computational resources for the MMseqs2 MSA server.

- Richard Evans for helping to benchmark the ColabFold's Alphafold-multimer support.

- [David Koes](https://github.com/dkoes) for his awesome [py3Dmol](https://3dmol.csb.pitt.edu/) plugin, without whom these notebooks would be quite boring!

- Do-Yoon Kim for creating the ColabFold logo.

- A colab by Sergey Ovchinnikov ([@sokrypton](https://twitter.com/sokrypton)), Milot Mirdita ([@milot_mirdita](https://twitter.com/milot_mirdita)) and Martin Steinegger ([@thesteinegger](https://twitter.com/thesteinegger)).
```
