## supplement_v6 for "Confidence Without Verification: Screening pLDDT Unreliability in AlphaFold2 Fold-Switching Predictions": README.md

# Supplemental Code and Data

**Paper:** Confidence Without Verification: Why AlphaFold2's Learned Metrics Fail for Fold-Switching Proteins

**Author:** R. Thacker, Independent Researcher, Vancouver Island, BC, Canada

**Framework:** SDI (Structural Dependencies of Intelligence), Canadian Patent Application CA 3294110

**Source Data:** Porter Lab CF-random repository (doi:10.5281/zenodo.15596182)

---

## Directory Structure

```
supplement/
├── README.md                  (this file)
├── scripts/                   (analysis code)
│   ├── core/                  (scripts that produced paper results)
│   └── exploratory/           (diagnostic and development scripts)
├── data/                      (derived results and intermediate data)
├── reports/                   (analysis summaries)
├── notebooks/                 (ColabFold prediction notebook)
└── replication/               (infrastructure for replication packaging)
```

## Core Analysis Scripts

These scripts produced the results reported in the paper.

| Script | Paper Section | Purpose |
|--------|--------------|---------|
| `multi_threshold_fv.py` | Table 4, Fig 4 | Multi-threshold FalseVerify analysis across 18 proteins at TM thresholds 0.5, 0.6, 0.7, 0.8 |
| `run_analysis.py` | Methods §3 | Main entry point: orchestrates population-level and per-residue SDI analysis |
| `audit_trail.py` | Methods §3 | Epistemic constraint layer — tags every claim as VERIFIED, DERIVED, ASSUMED, or FLAGGED |
| `tier1_analysis.py` | Methods §3.1 | Tier 1: classifies FSR residues using internal metrics only (pLDDT, PAE) |
| `tier2_analysis.py` | Methods §3.2, Table 2 | Tier 2: structural SDI typing with Kabsch alignment against experimental references |
| `population_typing.py` | Table 1, Fig 1 | Population-level SDI classification (FalseVerify, SelfCatch, Correct, OverRefusal) |
| `extract_and_screen.py` | Table 3 | E. coli blind screening: extracts PSE files, computes per-residue pLDDT, flags FV risk |
| `sdi_deviation_tracer.py` | Results §4 | Full per-protein SDI deviation tracing across Porter benchmark |

## Exploratory and Diagnostic Scripts

These supported development and verification but are not required to reproduce paper results.

| Script | Purpose |
|--------|---------|
| `blindfold_analysis.py` | Early CPE filter prototype applied to 5 benchmark cases |
| `vpc_porter_analysis.py` | V_PC (cross-cluster variance) analysis pipeline |
| `vpc_porter_complete.py` | V_PC analysis using aggregated .npy arrays |
| `verify_bccipa.py` | Verification of BCCIPα TM-score claims against raw data |
| `verify_sa1.py` | Verification of Sa1 TM-score claims against raw data |
| `extract_pse.py` | PSE extraction via PyMOL (requires PyMOL installation) |
| `extract_pse_v2.py` | PSE extraction without PyMOL dependency |
| `probe_pse.py` – `probe_pse_v4.py` | Iterative PSE format reverse-engineering (development trail) |
| `find_mapping.py` | Protein-to-index mapping in Porter data arrays |
| `analyze_npy.py` | Inspection of aggregated .npy TM-score files |
| `diagnose_data.py` | Directory structure diagnostics |
| `search_broader.py` | File discovery across BlindFold directory tree |

## Data Files

### Paper Results
| File | Paper Section | Description |
|------|--------------|-------------|
| `multi_threshold_fv_results.csv` | Table 4 | Full multi-threshold results for all 18 proteins × 4 thresholds |
| `multi_threshold_fv_table.txt` | Table 4 | Formatted table as printed in paper |
| `full_18protein_sdi_results.csv` | Table 1 | Complete SDI classification for 18 benchmark proteins |
| `afcluster_sdi_results.csv` | Table 1 | AFcluster-level SDI results |
| `perresidue_figure_data.csv` | Fig 3, Table 2 | Per-residue pLDDT and variance data for figure generation |
| `ecoli_sdi_screening.csv` | Table 3 | 52 E. coli candidate screening results |

### Intermediate Data
| File | Description |
|------|-------------|
| `vpc_results.csv` | V_PC cross-cluster variance scores |
| `blindfold_v0_1_flags.csv` | Early BLINDFOLD prototype flags |
| `blindfold_v0_1_examples_overconfident_strict.csv` | Overconfident prediction examples |
| `blindfold_v0_2_predicted_rankings.csv` | BLINDFOLD v0.2 rankings |
| `blindfold_v0_2_top9_predictions.csv` | Top 9 prioritized candidates |
| `tmscores_top5.txt` | Top-5 model TM-scores (fold1 + fold2) |
| `tmscores_a_top5.txt` | Top-5 model TM-scores (alternate reference) |
| `tmscores_unrelaxed.txt` | Full unrelaxed TM-scores |
| `tmscores_a_unrelaxed.txt` | Full unrelaxed TM-scores (alternate) |
| `trace_index_2oug_C_0_014.csv` | RfaH trace index for run 0_014 |
| `trace_metrics_2oug_C_0_014.json` | RfaH confidence metrics for run 0_014 |
| `XCL1_trace_index.csv` | XCL1 prediction trace index |
| `extraction_summary.csv` | PSE extraction summary for E. coli set |

## Reports

| File | Description |
|------|-------------|
| `cross_protein_sdi_report.md` | Cross-protein analysis confirming five-pattern taxonomy |
| `ecoli_sdi_screening_report.md` | Full E. coli screening report with prioritized candidates |
| `XCL1_TRACE_SUMMARY.md` | XCL1 case study — template-free vs template-guided failure modes |

## Notebook

| File | Description |
|------|-------------|
| `AlphaFold2.ipynb` | ColabFold notebook used for structure predictions |

## Requirements

- Python 3.8+
- NumPy
- Standard library (csv, json, os, math, struct, pickle, glob)
- Optional: pandas (for vpc_porter_complete.py), PyMOL (for extract_pse.py)

## Reproducing Paper Results

### Table 1 (Benchmark Classification)
Source data: Porter Lab CF-random repository `Data/Fold-switch_hits-AFcluster/`
```
python run_analysis.py population --whole-tm tmscores_all.npy --fsr-tm tmscores_fs_all.npy
```

### Table 4 (Multi-Threshold Analysis)
```
python multi_threshold_fv.py
```
Reads 18 protein directories from CF-random AFcluster data. Update `BASE_DIR` in script to local path.

### Table 3 (E. coli Screening)
```
python extract_and_screen.py
```
Reads 52 PSE files from CF-random `Data/E_coli/`. Update `base` path in script.

### Per-Residue Analysis (Table 2, Figure 3)
```
python run_analysis.py per-residue \
    --pred prediction.pdb \
    --ref1 fold1.pdb --ref1-label "2oug_C alpha" \
    --ref2 fold2.pdb --ref2-label "6c6s_D beta" \
    --scores scores.json \
    --fsr 115-162
```

## Notes

- All scripts contain hardcoded Windows paths from the original analysis environment. Update `BASE_DIR` / `base` / `data_dir` variables to your local data paths.
- The `probe_pse_v*.py` series documents the iterative development of PSE file parsing without PyMOL — retained for transparency.
- The `audit_trail.py` module implements the SDI-UE epistemic constraint layer (λ[C ≥ P ≥ E]). Every analysis function that imports it tags claims by verification status.

## License

Research use permitted. Contact author for commercial licensing.
SDI framework: Canadian Patent Application CA 3294110.

## Citation

Thacker, R. (2026). Confidence Without Verification: Why AlphaFold2's Learned Metrics Fail for Fold-Switching Proteins. Preprint v6.
