## supplement_v6 for "Confidence Without Verification: Screening pLDDT Unreliability in AlphaFold2 Fold-Switching Predictions": export_supplement.py

```
"""Export supplement tables.

Expected outputs:
- supplement/trace_index.csv (one row per trace with key metrics)
- supplement/summary_tables.csv (case-level aggregates)
"""
def main():
    raise NotImplementedError("Implement after metrics/ schemas are finalized.")

if __name__ == "__main__":
    main()
```
