## supplement_v6 for "Confidence Without Verification: Screening pLDDT Unreliability in AlphaFold2 Fold-Switching Predictions": make_manifest.py

```
import os, json, hashlib, argparse

def sha256_file(path, chunk=1<<20):
    h = hashlib.sha256()
    with open(path, "rb") as f:
        while True:
            b = f.read(chunk)
            if not b:
                break
            h.update(b)
    return h.hexdigest()

def main(raw_dir):
    manifest = {"raw_dir": raw_dir, "files": []}
    for root, _, files in os.walk(raw_dir):
        for name in files:
            if name == ".keep":
                continue
            p = os.path.join(root, name)
            rel = os.path.relpath(p, raw_dir)
            manifest["files"].append({
                "path": rel,
                "bytes": os.path.getsize(p),
                "sha256": sha256_file(p),
            })
    manifest["files"].sort(key=lambda x: x["path"])
    out = os.path.join(raw_dir, "manifest.json")
    with open(out, "w", encoding="utf-8") as f:
        json.dump(manifest, f, indent=2)
    print(f"Wrote {out} with {len(manifest['files'])} entries.")

if __name__ == "__main__":
    ap = argparse.ArgumentParser()
    ap.add_argument("--raw_dir", default="data/raw")
    args = ap.parse_args()
    main(args.raw_dir)
```
