## supplement_v6 for "Confidence Without Verification: Screening pLDDT Unreliability in AlphaFold2 Fold-Switching Predictions": ecoli_sdi_screening_report.md

# E. coli Fold-Switching Candidate Screening: SDI/CPE Analysis

## Summary

52 putative fold-switching proteins identified by CF-random blind search (Porter et al. 2025) were screened for confidence integrity using SDI framework analysis. Each PSE file contains two conformations — "Dominant" (standard AF2) and an alternative found by CF-random — with per-residue pLDDT scores.

**Key finding**: The confidence-accuracy decoupling discovered in RfaH/KaiB generalizes to the full E. coli candidate set, but reveals a **fourth pattern** not seen in the benchmark: inverted confidence, where the alternative conformation is MORE confident than the dominant.

## Population Statistics

| Metric | Value |
|--------|-------|
| Total candidates screened | 52 |
| Mean dominant pLDDT | 78.8 |
| Mean alternative pLDDT | 78.7 |
| Mean absolute confidence gap | 14.8 |
| Candidates with significant gap (|Δ| > 5) | 43/52 (83%) |

## SDI Category Breakdown

| Category | Count | % | CPE Regime | Interpretation |
|----------|-------|---|------------|----------------|
| HIGH FalseVerify risk | 14 | 27% | E>P | Dominant memorized, alternative unreliable |
| MODERATE FalseVerify risk | 4 | 8% | E>P (mild) | Some memorization bias |
| Both stable | 9 | 17% | C≥P≥E | Best wet-lab candidates |
| INVERTED confidence | 19 | 37% | P>E? | Alternative MORE confident — see below |
| Low confidence | 1 | 2% | Uncertain | Insufficient signal |
| Mild / Mixed | 5 | 10% | Variable | Requires per-residue analysis |

## The Four Patterns

### Pattern 1: HIGH FalseVerify (14 proteins, 27%)

Dominant conformation has pLDDT > 70 with Δ > +15 over alternative. This is the RfaH/KaiB signature: AF2 is very confident about the dominant fold but the alternative shows dramatically lower confidence, suggesting the alternative prediction may be unreliable memorization artifact.

**These should be treated with CAUTION before wet-lab investment.**

| WP ID | Res | Dom pLDDT | Alt pLDDT | Δ | Max Δ | % Differ |
|-------|-----|-----------|-----------|---|-------|----------|
| WP_001282181 | 121 | 84.5 | 52.2 | +32.3 | 61.0 | 97% |
| WP_000847304 | 109 | 95.0 | 63.9 | +31.1 | 52.6 | 100% |
| WP_000024392 | 83 | 92.9 | 63.8 | +29.0 | 52.8 | 99% |
| WP_000617148 | 92 | 95.2 | 68.8 | +26.3 | 60.5 | 100% |
| WP_000015473 | 121 | 94.4 | 71.0 | +23.4 | 58.3 | 95% |
| WP_001295442 | 232 | 81.2 | 58.8 | +22.4 | 53.7 | 76% |
| WP_000994516 | 62 | 94.4 | 72.4 | +21.9 | 41.3 | 97% |
| WP_001272149 | 183 | 88.3 | 68.7 | +19.6 | 45.0 | 79% |
| WP_000064148 | 77 | 90.9 | 71.5 | +19.4 | 31.3 | 90% |
| WP_001119863 | 126 | 96.4 | 78.1 | +18.3 | 51.1 | 71% |
| WP_000705622 | 183 | 85.5 | 67.3 | +18.2 | 54.6 | 82% |
| WP_000807125 | 119 | 95.2 | 77.5 | +17.7 | 53.0 | 67% |
| WP_001280953 | 124 | 88.6 | 72.4 | +16.2 | 39.2 | 79% |
| WP_000841554 | 45 | 88.5 | 72.4 | +16.1 | 36.1 | 53% |

### Pattern 2: INVERTED Confidence (19 proteins, 37%)

The alternative conformation has HIGHER pLDDT than the dominant — the opposite of FalseVerify. This is a novel finding not observed in the benchmark proteins.

**Interpretation**: CF-random's "blind mode" labels the first conformation as "Dominant" based on clustering, but in these cases the alternative may actually be the more physically plausible structure. The dominant label may be an artifact of the clustering algorithm, not a ground truth.

**Top inverted cases** (alternative pLDDT > 90):

| WP ID | Dom pLDDT | Alt pLDDT | Δ |
|-------|-----------|-----------|---|
| WP_001296901 | 59.8 | 94.9 | -35.0 |
| WP_001070563 | 65.1 | 93.3 | -28.2 |
| WP_000323025 | 73.5 | 95.4 | -21.9 |
| WP_001272856 | 75.5 | 96.0 | -20.5 |
| WP_001270286 | 77.5 | 96.9 | -19.4 |
| WP_000190655 | 75.2 | 94.3 | -19.1 |
| WP_001279084 | 78.5 | 94.6 | -16.1 |

### Pattern 3: Both Conformations Stable (9 proteins, 17%)

Both dominant and alternative show similar pLDDT (|Δ| < 5). This matches the Mad2 pattern where both conformations are captured with comparable confidence. **These are the strongest candidates for genuine fold switching** and should be prioritized for experimental validation.

| WP ID | Res | Dom pLDDT | Alt pLDDT | Δ | % Differ |
|-------|-----|-----------|-----------|---|----------|
| WP_001241339 | 128 | 78.2 | 77.4 | +0.8 | 35% |
| WP_001002059 | 124 | 62.6 | 65.2 | -2.6 | 38% |
| WP_001281772 | 78 | 80.3 | 77.3 | +3.0 | 32% |
| WP_000955366 | 49 | 80.4 | 77.3 | +3.1 | 61% |
| WP_001151233 | 137 | 80.9 | 84.3 | -3.4 | 42% |
| WP_000881326 | 205 | 78.1 | 81.6 | -3.5 | 13% |
| WP_001129553 | 176 | 88.2 | 84.7 | +3.5 | 7% |
| WP_000803992 | 87 | 69.0 | 72.6 | -3.6 | 51% |
| WP_000648420 | 78 | 80.3 | 75.4 | +4.9 | 10% |

**Note**: WP_000955366 is unique — it has **7 objects** (6 alternative conformations), suggesting a highly polymorphic protein. WP_001129553 has the tightest gap (Δ=+3.5) with the highest average pLDDT across both conformations (88.2/84.7), making it a particularly strong candidate.

### Pattern 4: Low Confidence (1 protein)

WP_000134927 has dom_pLDDT=40.2 — below the reliability threshold. Neither conformation is trustworthy.

---

## CPE Framework Validation

The three-pattern taxonomy from the benchmark (RfaH, KaiB, Mad2) extends to this population with a fourth pattern:

| Benchmark Pattern | E. coli Equivalent | CPE Regime | N |
|-------------------|--------------------|------------|---|
| RfaH/KaiB (FalseVerify) | High FV risk | E>P | 14+4 |
| Mad2 (both folds) | Both stable | C≥P≥E | 9 |
| — (not seen) | Inverted confidence | P>E or label error | 19 |

The **perfect 50/50 split** (26 proteins each direction) in the confidence gap is itself informative: it means pLDDT is essentially UNCORRELATED with which conformation is "correct" in the fold-switching context. This is the population-level signature of systematic confidence-accuracy decoupling.

## Recommendation to Porter Lab

**Priority 1 — Wet lab (9 proteins)**: The "both stable" candidates (WP_001129553, WP_001241339, WP_001281772, WP_000955366, WP_001151233, WP_000881326, WP_001002059, WP_000803992, WP_000648420) have the strongest confidence integrity signatures and should be prioritized for experimental validation.

**Priority 2 — Re-examine (19 proteins)**: The inverted cases may reflect labeling artifacts in CF-random's blind mode clustering. Cross-referencing with ACE coevolutionary data and Foldseek assignments would clarify whether the "Dominant" or "Alternative" label is correct.

**Priority 3 — Caution (14 proteins)**: The high FV risk candidates have the classic memorization signature. The alternative conformations predicted by CF-random may not be reliable for these proteins, even though they passed the ACE/Foldseek filters.

---

*Analysis: R. Thacker, SDI Framework / CPE Constraint Hierarchy*
*Data: Porter Lab CF-random repository (doi:10.5281/zenodo.15596182)*
*52 E. coli putative fold-switching proteins, 104 PDB conformations analyzed*
*February 2026*
