## supplement_v6 for "Confidence Without Verification: Screening pLDDT Unreliability in AlphaFold2 Fold-Switching Predictions": XCL1_TRACE_SUMMARY.md

# XCL1 Trace Acquisition Summary
## Date: 2026-01-05

## Raw Data Collected

| Condition | MSA Depth | Artifacts |
|-----------|-----------|-----------|
| Deep | 562 seqs | XCL1_deep_54f38_result.zip |
| U100-008 | 101 seqs | XCL1_U100008_54f38_0_result.zip |
| U10-008 | 11 seqs | XCL1_U10008_54f38_0_result.zip |

## Confidence Metrics (Rank 1)

| Condition | MSA Depth | pLDDT | pTM | Verdict |
|-----------|-----------|-------|-----|---------|
| **Deep** | 562 seqs | 40.3 | 0.300 | FAIL |
| **U100** | 101 seqs | 38.7 | 0.220 | FAIL |
| **U10** | 11 seqs | 33.6 | 0.160 | FAIL |

## Key Finding

**XCL1 is a ROBUST FAILURE for template-free AlphaFold2:**

1. Confidence stays uniformly LOW across all MSA depths (pLDDT 33-40)
2. No fold-switching behavior observed (unlike RfaH)
3. AF2 correctly signals uncertainty via low pLDDT and pTM

## Comparison to RfaH

| Protein | Deep pLDDT | Deep TM(fold1) | U10 pLDDT | U10 TM(fold2) | Behavior |
|---------|------------|----------------|-----------|---------------|----------|
| **RfaH** | ~80 | 0.74 (α) | ~80 | 0.75 (β) | **Fold-switches** |
| **XCL1** | ~40 | ? | ~34 | ? | **Fails uniformly** |

## Critical Distinction: Template-Free vs AF2Rank

| Method | XCL1 pLDDT | Interpretation |
|--------|------------|----------------|
| **AF2Rank** (template-guided) | 88.9 | High confidence WITH template |
| **ColabFold** (template-free) | 40.3 | Low confidence WITHOUT template |

The AF2Rank "calibration gap" (pLDDT 88.9 vs TM 0.62) reflects template-guided scoring.
Our template-free predictions show AF2 correctly recognizes uncertainty.

## Implications for BLINDFOLD

XCL1 represents a **different failure mode** than expected:

- **Expected:** High confidence, wrong fold (miscalibration)
- **Observed:** Low confidence, no fold predicted (calibrated uncertainty)

This suggests AF2's internal confidence (pLDDT, pTM) may be more reliable than 
AF2Rank's template-preference scores for detecting genuinely difficult targets.

## Next Steps

1. Compute TM-scores against reference structures (requires TM-align/USalign)
2. Compare template-free vs template-guided confidence patterns
3. Update BLINDFOLD protocol to distinguish these failure modes

## Artifacts

- `XCL1_trace_index.csv` - Consolidated metrics for all 15 predictions
- Raw result zips contain PDBs, score JSONs, and PAE matrices
