## supplement_v6 for "Confidence Without Verification: Screening pLDDT Unreliability in AlphaFold2 Fold-Switching Predictions": audit_trail.py

```
"""
SDI-UE Audit Trail: Epistemic Constraint Layer

Every claim produced by this tool is tagged with its epistemic status:
  VERIFIED  — directly measured from input data
  DERIVED   — computed from verified inputs (Derivation ≠ Verification)
  ASSUMED   — depends on an unverifiable assumption (flagged)
  FLAGGED   — potential E>P violation detected

The audit trail is not optional. It is the tool's primary output.
The numbers are secondary to knowing which numbers you can trust.

Reference: SDI v1.1 §4 (violation taxonomy), UE v1.9 §7.4 (Transmission Gap)
"""

from dataclasses import dataclass, field
from typing import List, Optional
from datetime import datetime
import json


@dataclass
class AuditEntry:
    """Single auditable claim."""
    claim: str
    status: str  # VERIFIED | DERIVED | ASSUMED | FLAGGED
    verification_path: str  # How this was established
    depends_on: List[str] = field(default_factory=list)  # Other claims this relies on
    flag_reason: Optional[str] = None  # If ASSUMED or FLAGGED, why

    def to_dict(self):
        d = {
            'claim': self.claim,
            'status': self.status,
            'verification_path': self.verification_path,
        }
        if self.depends_on:
            d['depends_on'] = self.depends_on
        if self.flag_reason:
            d['flag_reason'] = self.flag_reason
        return d


class AuditTrail:
    """
    Accumulates auditable claims throughout an analysis run.

    Usage:
        audit = AuditTrail()
        audit.verified("Total predictions = 212018", "len(tmscores_fs_all.npy)")
        audit.derived("FalseVerify rate = 40.4%", "85643/212018", ["Total predictions = 212018"])
        audit.assumed("Array pairing", "First N entries assumed paired", "Different array lengths")
        audit.report()
    """

    def __init__(self):
        self.entries: List[AuditEntry] = []
        self.start_time = datetime.now()

    def verified(self, claim: str, verification_path: str):
        """Claim directly measured from input data."""
        self.entries.append(AuditEntry(
            claim=claim,
            status="VERIFIED",
            verification_path=verification_path
        ))

    def derived(self, claim: str, verification_path: str, depends_on: List[str] = None):
        """Claim computed from verified inputs. Derivation ≠ Verification."""
        self.entries.append(AuditEntry(
            claim=claim,
            status="DERIVED",
            verification_path=verification_path,
            depends_on=depends_on or []
        ))

    def assumed(self, claim: str, verification_path: str, flag_reason: str):
        """Claim depends on unverifiable assumption."""
        self.entries.append(AuditEntry(
            claim=claim,
            status="ASSUMED",
            verification_path=verification_path,
            flag_reason=flag_reason
        ))

    def flagged(self, claim: str, verification_path: str, flag_reason: str):
        """Potential E>P violation detected."""
        self.entries.append(AuditEntry(
            claim=claim,
            status="FLAGGED",
            verification_path=verification_path,
            flag_reason=flag_reason
        ))

    def has_flags(self) -> bool:
        """Are there any ASSUMED or FLAGGED entries?"""
        return any(e.status in ("ASSUMED", "FLAGGED") for e in self.entries)

    def get_flags(self) -> List[AuditEntry]:
        """Return only ASSUMED and FLAGGED entries."""
        return [e for e in self.entries if e.status in ("ASSUMED", "FLAGGED")]

    def report(self) -> str:
        """Human-readable audit report."""
        lines = []
        lines.append("=" * 70)
        lines.append("SDI-UE AUDIT TRAIL")
        lines.append(f"Generated: {datetime.now().isoformat()}")
        lines.append(f"Framework: λ[C ≥ P ≥ E]")
        lines.append(f"Constraint: No claim may exceed what preservation can verify")
        lines.append("=" * 70)
        lines.append("")

        # Group by status
        for status in ["VERIFIED", "DERIVED", "ASSUMED", "FLAGGED"]:
            group = [e for e in self.entries if e.status == status]
            if not group:
                continue
            lines.append(f"[{status}] ({len(group)} claims)")
            lines.append("-" * 50)
            for e in group:
                lines.append(f"  • {e.claim}")
                lines.append(f"    Path: {e.verification_path}")
                if e.depends_on:
                    lines.append(f"    Depends on: {'; '.join(e.depends_on)}")
                if e.flag_reason:
                    lines.append(f"    ⚠ FLAG: {e.flag_reason}")
            lines.append("")

        # Summary
        flags = self.get_flags()
        if flags:
            lines.append("=" * 70)
            lines.append(f"⚠ {len(flags)} OPEN FLAG(S) — claims require external confirmation")
            lines.append("=" * 70)
            for f in flags:
                lines.append(f"  • [{f.status}] {f.claim}")
                lines.append(f"    {f.flag_reason}")
        else:
            lines.append("✓ No open flags. All claims verified or derived from verified inputs.")

        lines.append("")
        return "\n".join(lines)

    def to_json(self) -> str:
        """Machine-readable audit trail."""
        return json.dumps({
            'framework': 'SDI-UE λ[C ≥ P ≥ E]',
            'generated': datetime.now().isoformat(),
            'total_claims': len(self.entries),
            'verified': len([e for e in self.entries if e.status == "VERIFIED"]),
            'derived': len([e for e in self.entries if e.status == "DERIVED"]),
            'assumed': len([e for e in self.entries if e.status == "ASSUMED"]),
            'flagged': len([e for e in self.entries if e.status == "FLAGGED"]),
            'entries': [e.to_dict() for e in self.entries]
        }, indent=2)
```
