## supplement_v6 for "Confidence Without Verification: Screening pLDDT Unreliability in AlphaFold2 Fold-Switching Predictions": extract_and_screen.py

```
"""Extract and analyze all 52 E. coli PSE files for SDI screening."""
import sys, types, pickle, glob, os, struct

class SafeUnpickler(pickle.Unpickler):
    def find_class(self, module, name):
        safe = {'builtins','collections','collections.abc','copy_reg','copyreg','_codecs','codecs'}
        for s in safe:
            if module == s or module.startswith(s + '.'):
                try: return super().find_class(module, name)
                except: pass
        def make(*a, **kw):
            o = object.__new__(S)
            o._mc = f'{module}.{name}'; o._a = a
            return o
        class S:
            _mc = f'{module}.{name}'
            def __init__(s, *a, **kw): s._a = a
            def __setstate__(s, st):
                if isinstance(st, dict): s.__dict__.update(st)
                else: s._state = st
            def __reduce__(s): return (make, s._a)
        S.__name__ = name
        return S

def extract_pse(pse_file):
    """Extract objects from a PSE file. Returns list of (name, atoms, coords_per_state)."""
    with open(pse_file, 'rb') as f:
        data = SafeUnpickler(f, encoding='latin1').load()
    
    results = []
    names = data.get('names', [])
    
    for entry in names:
        if entry is None or not isinstance(entry, list) or len(entry) < 6:
            continue
        
        obj_name = entry[0]
        mol_data = entry[5]
        
        if not isinstance(mol_data, list) or len(mol_data) < 8:
            continue
        
        mol_type = mol_data[1]  # 1 = molecule
        if mol_type != 1:
            continue
        
        n_atoms = mol_data[2]
        atom_list = mol_data[7]  # atom info
        coord_sets_container = mol_data[4]  # coordinate sets
        
        if not isinstance(atom_list, list) or not atom_list:
            continue
        
        # Extract atom info: resn, chain, resi, atom_name, b_factor, element
        atoms = []
        for ai in atom_list:
            if not isinstance(ai, (list, tuple)) or len(ai) < 15:
                continue
            atoms.append({
                'resi': ai[3],        # residue number (string)
                'resn': ai[5],        # residue name
                'name': ai[6],        # atom name
                'chain': ai[1],       # chain
                'b': ai[14],          # B-factor = pLDDT
                'elem': ai[7] if len(ai) > 7 else ai[6][0],  # element
            })
        
        # Extract coordinates for each state
        states_coords = []
        if isinstance(coord_sets_container, list):
            for cs in coord_sets_container:
                if cs is None:
                    continue
                # cs is a list, cs[0] should have coordinates
                if isinstance(cs, list) and len(cs) > 0:
                    coord_data = cs[0]
                    if isinstance(coord_data, bytes):
                        nf = len(coord_data) // 4
                        coords = struct.unpack(f'{nf}f', coord_data)
                        states_coords.append(coords)
                    elif isinstance(coord_data, (list, tuple)):
                        states_coords.append(coord_data)
        
        results.append({
            'name': obj_name,
            'n_atoms': len(atoms),
            'atoms': atoms,
            'n_states': len(states_coords),
            'coords': states_coords
        })
    
    return results

def write_pdb(filename, atoms, coords):
    """Write a PDB file from atom info and coordinates."""
    with open(filename, 'w') as f:
        for i, atom in enumerate(atoms):
            if i * 3 + 2 >= len(coords):
                break
            x = coords[i*3]
            y = coords[i*3 + 1]
            z = coords[i*3 + 2]
            
            name = atom['name']
            resn = atom['resn'][:3]
            chain = atom['chain'][:1] if atom['chain'] else 'A'
            resi = atom['resi']
            b = atom['b']
            elem = atom['elem'][:2] if atom['elem'] else name[0]
            
            # Format atom name (4 chars, left-justified if 4 chars, else space-prefixed)
            if len(name) < 4:
                name_fmt = f' {name:<3s}'
            else:
                name_fmt = f'{name:<4s}'
            
            try:
                resi_int = int(resi)
            except:
                resi_int = i + 1
            
            f.write(f'ATOM  {i+1:5d} {name_fmt}{resn:>3s} {chain:1s}{resi_int:4d}    '
                    f'{x:8.3f}{y:8.3f}{z:8.3f}{1.0:6.2f}{b:6.2f}          {elem:>2s}\n')
        f.write('END\n')

def get_ca_plddt(atoms):
    """Get per-residue pLDDT from CA atoms."""
    ca_plddt = []
    for a in atoms:
        if a['name'] == 'CA':
            ca_plddt.append(a['b'])
    return ca_plddt

# =========================================================
# MAIN
# =========================================================
base = r'C:\Users\robin\Desktop\MAGIC-Protein-Fold\Phase 2 - Porter\data\cf_random_repo\CF-random_software\Data\E_coli'
os.chdir(base)
os.makedirs('extracted_pdbs', exist_ok=True)

all_results = []

for pse_file in sorted(glob.glob('*.pse')):
    wp = pse_file.replace('.pse', '')
    
    try:
        objects = extract_pse(pse_file)
    except Exception as e:
        print(f'{wp}: ERROR - {e}')
        all_results.append({'wp': wp, 'error': str(e)})
        continue
    
    if not objects:
        print(f'{wp}: No molecular objects found')
        all_results.append({'wp': wp, 'error': 'no objects'})
        continue
    
    # Identify dominant vs alternative
    dominant = None
    alternatives = []
    
    for obj in objects:
        if obj['name'].lower() == 'dominant':
            dominant = obj
        else:
            alternatives.append(obj)
    
    if dominant is None and objects:
        dominant = objects[0]
        alternatives = objects[1:]
    
    # Extract pLDDT for dominant
    dom_plddt = get_ca_plddt(dominant['atoms'])
    dom_mean = sum(dom_plddt) / len(dom_plddt) if dom_plddt else 0
    dom_n_res = len(dom_plddt)
    
    # Write dominant PDB
    if dominant['coords']:
        write_pdb(f'extracted_pdbs/{wp}_dominant.pdb', dominant['atoms'], dominant['coords'][0])
    
    # Process alternatives
    alt_info = []
    for alt in alternatives:
        alt_plddt = get_ca_plddt(alt['atoms'])
        alt_mean = sum(alt_plddt) / len(alt_plddt) if alt_plddt else 0
        
        if alt['coords']:
            write_pdb(f'extracted_pdbs/{wp}_{alt["name"]}.pdb', alt['atoms'], alt['coords'][0])
        
        # Compare per-residue pLDDT between dominant and alternative
        # (only for shared residues by position)
        min_len = min(len(dom_plddt), len(alt_plddt))
        if min_len > 0:
            dom_sub = dom_plddt[:min_len]
            alt_sub = alt_plddt[:min_len]
            
            # Regions where alternative has LOWER pLDDT
            drops = [dom_sub[i] - alt_sub[i] for i in range(min_len)]
            mean_drop = sum(drops) / len(drops)
            max_drop = max(drops)
            
            # Regions where alternative has HIGHER pLDDT
            gains = [alt_sub[i] - dom_sub[i] for i in range(min_len)]
            mean_gain = sum(g for g in gains if g > 0) / max(1, sum(1 for g in gains if g > 0))
            
            # Count residues with significant pLDDT difference (>10)
            n_sig_diff = sum(1 for d in drops if abs(d) > 10)
            pct_sig_diff = n_sig_diff / min_len * 100
        else:
            mean_drop = 0
            max_drop = 0
            mean_gain = 0
            n_sig_diff = 0
            pct_sig_diff = 0
        
        alt_info.append({
            'name': alt['name'],
            'n_res': len(alt_plddt),
            'mean_plddt': alt_mean,
            'mean_drop': mean_drop,
            'max_drop': max_drop,
            'mean_gain': mean_gain,
            'n_sig_diff': n_sig_diff,
            'pct_sig_diff': pct_sig_diff,
        })
    
    result = {
        'wp': wp,
        'n_objects': len(objects),
        'dom_residues': dom_n_res,
        'dom_mean_plddt': dom_mean,
        'alternatives': alt_info,
    }
    all_results.append(result)
    
    # Print summary
    alt_str = ''
    if alt_info:
        a = alt_info[0]
        alt_str = (f'  alt_pLDDT={a["mean_plddt"]:.1f}  '
                   f'Δ={a["mean_drop"]:+.1f}  '
                   f'maxΔ={a["max_drop"]:.1f}  '
                   f'{a["pct_sig_diff"]:.0f}% residues differ >10')
    
    print(f'{wp}: {len(objects)} obj, {dom_n_res} res, dom_pLDDT={dom_mean:.1f}{alt_str}')

# =========================================================
# SUMMARY TABLE
# =========================================================
print('\n' + '='*100)
print('E. COLI SDI SCREENING SUMMARY')
print('='*100)
print(f'\n{"WP_ID":<18} {"Res":>4} {"Dom_pLDDT":>10} {"Alt_pLDDT":>10} {"MeanΔ":>7} {"MaxΔ":>6} {"%Diff>10":>8} {"Flag":<20}')
print('-'*100)

flagged_high = []
flagged_low = []

for r in all_results:
    if 'error' in r:
        print(f'{r["wp"]:<18} ERROR: {r["error"]}')
        continue
    
    wp = r['wp']
    dom_p = r['dom_mean_plddt']
    n_res = r['dom_residues']
    
    if not r['alternatives']:
        print(f'{wp:<18} {n_res:>4} {dom_p:>10.1f} {"N/A":>10} {"N/A":>7} {"N/A":>6} {"N/A":>8} DOMINANT ONLY')
        continue
    
    a = r['alternatives'][0]
    alt_p = a['mean_plddt']
    md = a['mean_drop']
    mx = a['max_drop']
    pct = a['pct_sig_diff']
    
    # SDI FalseVerify flag logic:
    # HIGH RISK: dominant pLDDT high, alternative much lower → classic FalseVerify
    # LOW RISK: both similar pLDDT → Mad2-like, both conformations plausible
    if dom_p > 70 and md > 15:
        flag = '*** HIGH FV RISK'
        flagged_high.append(wp)
    elif dom_p > 70 and md > 8:
        flag = '** MODERATE FV RISK'
        flagged_high.append(wp)
    elif dom_p > 60 and md > 5:
        flag = '* MILD FV RISK'
    elif abs(md) < 5 and dom_p > 60:
        flag = 'OK (both stable)'
        flagged_low.append(wp)
    elif dom_p < 50:
        flag = 'LOW CONF (both)'
    else:
        flag = 'MIXED'
    
    print(f'{wp:<18} {n_res:>4} {dom_p:>10.1f} {alt_p:>10.1f} {md:>+7.1f} {mx:>6.1f} {pct:>7.0f}% {flag}')

print('\n' + '='*100)
print(f'HIGH/MODERATE FV RISK: {len(flagged_high)} proteins')
for wp in flagged_high:
    print(f'  {wp}')
print(f'\nLOW FV RISK (both conformations stable): {len(flagged_low)} proteins')
for wp in flagged_low:
    print(f'  {wp}')
print(f'\nTotal processed: {len(all_results)}')
print(f'Total PDBs extracted: {len(glob.glob("extracted_pdbs/*.pdb"))}')

# Save detailed CSV
with open('ecoli_sdi_screening.csv', 'w') as f:
    f.write('wp_id,n_objects,n_residues,dom_mean_plddt,alt_name,alt_mean_plddt,mean_delta,max_delta,pct_diff_gt10,flag\n')
    for r in all_results:
        if 'error' in r:
            f.write(f'{r["wp"]},ERROR,,,,,,,,{r["error"]}\n')
            continue
        wp = r['wp']
        dom_p = r['dom_mean_plddt']
        n_res = r['dom_residues']
        n_obj = r['n_objects']
        if r['alternatives']:
            a = r['alternatives'][0]
            md = a['mean_drop']
            if dom_p > 70 and md > 15: flag = 'HIGH_FV'
            elif dom_p > 70 and md > 8: flag = 'MODERATE_FV'
            elif dom_p > 60 and md > 5: flag = 'MILD_FV'
            elif abs(md) < 5 and dom_p > 60: flag = 'OK'
            elif dom_p < 50: flag = 'LOW_CONF'
            else: flag = 'MIXED'
            f.write(f'{wp},{n_obj},{n_res},{dom_p:.1f},{a["name"]},{a["mean_plddt"]:.1f},{md:.1f},{a["max_drop"]:.1f},{a["pct_sig_diff"]:.1f},{flag}\n')
        else:
            f.write(f'{wp},{n_obj},{n_res},{dom_p:.1f},,,,,,DOMINANT_ONLY\n')

print('Saved: ecoli_sdi_screening.csv')
```
