## supplement_v6 for "Confidence Without Verification: Screening pLDDT Unreliability in AlphaFold2 Fold-Switching Predictions": multi_threshold_fv.py

```
#!/usr/bin/env python3
"""
Multi-Threshold FalseVerify Analysis
=====================================
Reads all 33 AFcluster CSVs and computes FV rates at TM thresholds 0.5, 0.6, 0.7, 0.8.

Usage:
    python multi_threshold_fv.py

Output:
    multi_threshold_fv_results.csv — full results for all proteins at all thresholds
    multi_threshold_fv_table.txt  — formatted table for the paper

Author: Robin (PI) + Claude (research assistant)
"""

import csv
import os
import math

# ============================================================
# CONFIGURATION — adjust this path to your local data
# ============================================================
BASE_DIR = r'C:\Users\robin\Desktop\MAGIC-Protein-Fold\Phase 2 - Porter\data\cf_random_repo\CF-random_software\Data\Fold-switch_hits-AFcluster'

TM_THRESHOLDS = [0.5, 0.6, 0.7, 0.8]

# Protein display names mapped to folder names
# (folder names match the AFcluster directory structure)
PROTEIN_MAP = {
    'AIAT': 'A1AT',
    'CaBP': 'CaBP',
    'Cas9': 'Cas9',
    'COMT': 'COMT',
    'CRKL': 'CRKL',
    'Cwc2': 'Cwc2',
    'Fab': 'Fab',
    'FraC': 'FraC',
    'FUS_HENDH': 'FUS_HENDH',
    'GP2': 'GP2',
    'IscA': 'IscA',
    'KaiB': 'KaiB',
    'Mad2': 'Mad2',
    'MinE': 'MinE',
    'Nrp2': 'Nrp2',
    'OxyR': 'OxyR',
    'PimA': 'PimA',
    'RfaH': 'RfaH',
}

# SDI pattern from verified Table 1 (for ordering)
SDI_PATTERN = {
    'GP2': 'SEVERE FV',
    'KaiB': 'SEVERE FV',
    'PimA': 'SEVERE FV',
    'FraC': 'MODERATE FV',
    'Nrp2': 'MODERATE FV',
    'FUS_HENDH': 'MILD FV',
    'RfaH': 'MILD FV',
    'CaBP': 'BOTH FOLDS',
    'Cas9': 'BOTH FOLDS',
    'CRKL': 'BOTH FOLDS',
    'IscA': 'BOTH FOLDS',
    'OxyR': 'BOTH FOLDS',
    'A1AT': 'MAINTAINED',
    'COMT': 'MAINTAINED',
    'Cwc2': 'MAINTAINED',
    'Fab': 'MAINTAINED',
    'Mad2': 'MAINTAINED',
    'MinE': 'MAINTAINED',
}

PATTERN_ORDER = ['SEVERE FV', 'MODERATE FV', 'MILD FV', 'BOTH FOLDS', 'MAINTAINED']


def analyze_csv(filepath, tm_threshold=0.5):
    """
    Read a single AFcluster CSV and classify predictions.
    
    CSV format (no header): prediction_id, pLDDT_percentile, TM_fold1, TM_fold2
    
    Returns dict with counts and FV rate.
    """
    rows = []
    with open(filepath, 'r') as f:
        reader = csv.reader(f)
        for r in reader:
            if len(r) < 4:
                continue
            try:
                plddt_pct = float(r[1])
                tm1 = float(r[2])
                tm2 = float(r[3])
                rows.append((r[0].strip(), plddt_pct, tm1, tm2))
            except ValueError:
                continue
    
    n = len(rows)
    if n == 0:
        return None
    
    f1only = sum(1 for _, _, t1, t2 in rows if t1 >= tm_threshold and t2 < tm_threshold)
    f2only = sum(1 for _, _, t1, t2 in rows if t1 < tm_threshold and t2 >= tm_threshold)
    both = sum(1 for _, _, t1, t2 in rows if t1 >= tm_threshold and t2 >= tm_threshold)
    neither = sum(1 for _, _, t1, t2 in rows if t1 < tm_threshold and t2 < tm_threshold)
    
    # High-confidence = pLDDT percentile >= 50
    hc_rows = [(p, pl, t1, t2) for p, pl, t1, t2 in rows if pl >= 50.0]
    hc_n = len(hc_rows)
    hc_f1only = sum(1 for _, _, t1, t2 in hc_rows if t1 >= tm_threshold and t2 < tm_threshold)
    
    fv_rate = (hc_f1only / hc_n * 100) if hc_n > 0 else float('nan')
    
    return {
        'n': n,
        'f1only': f1only, 'f2only': f2only, 'both': both, 'neither': neither,
        'hc_n': hc_n, 'hc_f1only': hc_f1only, 'fv_rate': fv_rate,
    }


def main():
    print("=" * 80)
    print("Multi-Threshold FalseVerify Analysis")
    print("=" * 80)
    print(f"Base directory: {BASE_DIR}")
    print(f"TM thresholds: {TM_THRESHOLDS}")
    print()
    
    if not os.path.isdir(BASE_DIR):
        print(f"ERROR: Directory not found: {BASE_DIR}")
        print("Please update BASE_DIR in the script.")
        return
    
    # Collect all results
    all_results = []  # (protein, ref, tm_thresh, result_dict)
    
    for folder_name, protein_name in sorted(PROTEIN_MAP.items()):
        folder_path = os.path.join(BASE_DIR, folder_name)
        if not os.path.isdir(folder_path):
            print(f"WARNING: Folder not found: {folder_name}")
            continue
        
        # Find all _tmscores_fs_all.csv files
        csv_files = [f for f in os.listdir(folder_path) if f.endswith('_tmscores_fs_all.csv')]
        
        for csv_file in sorted(csv_files):
            ref = csv_file.replace('_tmscores_fs_all.csv', '')
            filepath = os.path.join(folder_path, csv_file)
            
            for tm_thresh in TM_THRESHOLDS:
                result = analyze_csv(filepath, tm_thresh)
                if result:
                    result['protein'] = protein_name
                    result['ref'] = ref
                    result['tm_threshold'] = tm_thresh
                    all_results.append(result)
    
    if not all_results:
        print("No results generated. Check file paths.")
        return
    
    # Count unique protein-ref combinations
    unique_combos = set((r['protein'], r['ref']) for r in all_results)
    print(f"Analyzed {len(unique_combos)} protein-reference combinations")
    print(f"Total analyses: {len(all_results)} ({len(unique_combos)} × {len(TM_THRESHOLDS)} thresholds)")
    
    # ============================================================
    # OUTPUT 1: Full CSV
    # ============================================================
    output_csv = 'multi_threshold_fv_results.csv'
    with open(output_csv, 'w', newline='') as f:
        writer = csv.writer(f)
        writer.writerow(['protein', 'ref', 'tm_threshold', 'n', 'f1only', 'f2only', 
                         'both', 'neither', 'hc_n', 'hc_f1only', 'fv_rate_hc'])
        for r in all_results:
            writer.writerow([
                r['protein'], r['ref'], r['tm_threshold'], r['n'],
                r['f1only'], r['f2only'], r['both'], r['neither'],
                r['hc_n'], r['hc_f1only'],
                f"{r['fv_rate']:.1f}" if not math.isnan(r['fv_rate']) else 'nan'
            ])
    print(f"\nFull results saved to: {output_csv}")
    
    # ============================================================
    # OUTPUT 2: Paper table — per-protein average FV% at each threshold
    # ============================================================
    
    # Group by protein, average FV% across references at each threshold
    from collections import defaultdict
    protein_data = defaultdict(lambda: defaultdict(list))
    
    for r in all_results:
        if not math.isnan(r['fv_rate']):
            protein_data[r['protein']][r['tm_threshold']].append(r['fv_rate'])
    
    # Sort by SDI pattern then by FV% at TM=0.5
    def sort_key(protein):
        pattern = SDI_PATTERN.get(protein, 'UNKNOWN')
        order = PATTERN_ORDER.index(pattern) if pattern in PATTERN_ORDER else 99
        fv_05 = sum(protein_data[protein].get(0.5, [0])) / max(1, len(protein_data[protein].get(0.5, [1])))
        return (order, -fv_05)
    
    sorted_proteins = sorted(protein_data.keys(), key=sort_key)
    
    # Print formatted table
    print("\n" + "=" * 80)
    print("TABLE: FalseVerify Rate (HC) at Multiple TM-Score Thresholds")
    print("=" * 80)
    
    header = f"{'Protein':<12} {'Pattern':<14}"
    for tm in TM_THRESHOLDS:
        header += f" {'FV% @'+str(tm):>10}"
    print(header)
    print("-" * 70)
    
    table_lines = []
    current_pattern = None
    
    for protein in sorted_proteins:
        pattern = SDI_PATTERN.get(protein, '?')
        
        if pattern != current_pattern:
            if current_pattern is not None:
                table_lines.append("")  # blank line between pattern groups
            current_pattern = pattern
        
        line = f"{protein:<12} {pattern:<14}"
        row_data = {'protein': protein, 'pattern': pattern}
        
        for tm in TM_THRESHOLDS:
            fv_rates = protein_data[protein].get(tm, [])
            if fv_rates:
                avg_fv = sum(fv_rates) / len(fv_rates)
                line += f" {avg_fv:>9.1f}%"
                row_data[f'fv_{tm}'] = avg_fv
            else:
                line += f" {'n/a':>10}"
                row_data[f'fv_{tm}'] = None
        
        table_lines.append(line)
    
    for tl in table_lines:
        print(tl)
    
    # Population totals
    print("-" * 70)
    pop_line = f"{'TOTAL':<12} {'':14}"
    for tm in TM_THRESHOLDS:
        total_hc = sum(r['hc_n'] for r in all_results if r['tm_threshold'] == tm)
        total_hc_fv = sum(r['hc_f1only'] for r in all_results if r['tm_threshold'] == tm)
        pop_fv = (total_hc_fv / total_hc * 100) if total_hc > 0 else 0
        pop_line += f" {pop_fv:>9.1f}%"
    print(pop_line)
    
    # Save table to text file
    table_file = 'multi_threshold_fv_table.txt'
    with open(table_file, 'w') as f:
        f.write("TABLE: FalseVerify Rate (HC) at Multiple TM-Score Thresholds\n")
        f.write("=" * 70 + "\n")
        f.write(header + "\n")
        f.write("-" * 70 + "\n")
        for tl in table_lines:
            f.write(tl + "\n")
        f.write("-" * 70 + "\n")
        f.write(pop_line + "\n")
    print(f"\nTable saved to: {table_file}")
    
    # ============================================================
    # KEY FINDING: Does the taxonomy ordering hold?
    # ============================================================
    print("\n" + "=" * 80)
    print("TAXONOMY INVARIANCE CHECK")
    print("=" * 80)
    
    for tm in TM_THRESHOLDS:
        print(f"\n  At TM >= {tm}:")
        
        pattern_means = {}
        for pattern in PATTERN_ORDER:
            proteins_in_pattern = [p for p in sorted_proteins if SDI_PATTERN.get(p) == pattern]
            fv_vals = []
            for p in proteins_in_pattern:
                fv_vals.extend(protein_data[p].get(tm, []))
            if fv_vals:
                pattern_means[pattern] = sum(fv_vals) / len(fv_vals)
            else:
                pattern_means[pattern] = 0.0
            print(f"    {pattern:<14}: mean FV = {pattern_means[pattern]:>6.1f}%")
        
        # Check ordering: SEVERE > MODERATE > MILD > BOTH ~ MAINTAINED
        severe = pattern_means.get('SEVERE FV', 0)
        moderate = pattern_means.get('MODERATE FV', 0)
        mild = pattern_means.get('MILD FV', 0)
        both_f = pattern_means.get('BOTH FOLDS', 0)
        maintained = pattern_means.get('MAINTAINED', 0)
        
        ordering_holds = severe >= moderate >= mild
        print(f"    Ordering SEVERE >= MODERATE >= MILD: {'YES' if ordering_holds else 'NO'}")


if __name__ == '__main__':
    main()
```
