## supplement_v6 for "Confidence Without Verification: Screening pLDDT Unreliability in AlphaFold2 Fold-Switching Predictions": population_typing.py

```
"""
Population-Level SDI Typing

Classifies AlphaFold predictions into SDI integrity categories
using whole-protein and FSR-specific TM-scores.

SDI Categories:
  FalseVerify     — whole TM ≥ threshold, FSR TM < threshold
                    (model confident, fold-switching region wrong)
  SelfCatch       — whole TM < threshold, FSR TM < threshold
                    (model uncertain, and indeed wrong)
  OverRefusal     — whole TM < threshold, FSR TM ≥ threshold
                    (model uncertain, but FSR actually ok)
  CorrectConfident — whole TM ≥ threshold, FSR TM ≥ threshold
                    (model confident, and correct)

Input: two .npy arrays (whole-protein TM-scores, FSR TM-scores)
Output: counts, rates, IR, and audit trail
"""

import numpy as np
from audit_trail import AuditTrail


def run_population_typing(
    whole_tm_path: str,
    fsr_tm_path: str,
    threshold: float = 0.5,
    audit: AuditTrail = None
) -> dict:
    """
    Classify predictions into SDI categories.

    Args:
        whole_tm_path: Path to whole-protein TM-score .npy file
        fsr_tm_path: Path to FSR-specific TM-score .npy file
        threshold: TM-score threshold for confident/uncertain (default 0.5)
        audit: AuditTrail instance (created if None)

    Returns:
        dict with counts, rates, IR, and audit trail
    """
    if audit is None:
        audit = AuditTrail()

    # Load arrays
    whole_tm = np.load(whole_tm_path)
    fsr_tm = np.load(fsr_tm_path)

    audit.verified(
        f"whole_tm array: shape={whole_tm.shape}, dtype={whole_tm.dtype}",
        f"np.load({whole_tm_path})"
    )
    audit.verified(
        f"fsr_tm array: shape={fsr_tm.shape}, dtype={fsr_tm.dtype}",
        f"np.load({fsr_tm_path})"
    )

    # Check array lengths
    if whole_tm.shape != fsr_tm.shape:
        n = min(len(whole_tm), len(fsr_tm))
        audit.assumed(
            f"Array pairing: using first {n} entries of each array",
            f"min(len(whole_tm)={len(whole_tm)}, len(fsr_tm)={len(fsr_tm)})",
            f"Arrays have different lengths ({len(whole_tm)} vs {len(fsr_tm)}). "
            f"Pairing assumption requires confirmation from data source. "
            f"Difference: {abs(len(whole_tm) - len(fsr_tm))} entries."
        )
        whole_tm = whole_tm[:n]
        fsr_tm = fsr_tm[:n]
    else:
        n = len(whole_tm)
        audit.verified(
            f"Array pairing: both arrays length {n}",
            "Identical shapes"
        )

    # Check for NaN / negative
    nan_whole = int(np.sum(np.isnan(whole_tm)))
    nan_fsr = int(np.sum(np.isnan(fsr_tm)))
    neg_whole = int(np.sum(whole_tm < 0))
    neg_fsr = int(np.sum(fsr_tm < 0))

    if nan_whole > 0 or nan_fsr > 0:
        audit.flagged(
            f"NaN values detected: whole_tm={nan_whole}, fsr_tm={nan_fsr}",
            "np.isnan() check",
            "NaN values will be excluded from classification"
        )
    if neg_whole > 0 or neg_fsr > 0:
        audit.flagged(
            f"Negative TM-scores: whole_tm={neg_whole}, fsr_tm={neg_fsr}",
            "negative value check",
            "Negative TM-scores are unusual and may indicate data issues"
        )

    # Classification
    false_verify = (whole_tm >= threshold) & (fsr_tm < threshold)
    self_catch = (whole_tm < threshold) & (fsr_tm < threshold)
    over_refusal = (whole_tm < threshold) & (fsr_tm >= threshold)
    correct_confident = (whole_tm >= threshold) & (fsr_tm >= threshold)

    fv = int(np.sum(false_verify))
    sc = int(np.sum(self_catch))
    orr = int(np.sum(over_refusal))
    cc = int(np.sum(correct_confident))

    # Sanity check
    assert fv + sc + orr + cc == n, f"Classification error: {fv}+{sc}+{orr}+{cc} != {n}"

    audit.derived(
        f"FalseVerify = {fv} ({fv/n*100:.1f}%)",
        f"(whole_tm >= {threshold}) & (fsr_tm < {threshold})",
        [f"whole_tm array: shape={whole_tm.shape}",
         f"fsr_tm array: shape={fsr_tm.shape}"]
    )
    audit.derived(
        f"SelfCatch = {sc} ({sc/n*100:.1f}%)",
        f"(whole_tm < {threshold}) & (fsr_tm < {threshold})"
    )
    audit.derived(
        f"OverRefusal = {orr} ({orr/n*100:.1f}%)",
        f"(whole_tm < {threshold}) & (fsr_tm >= {threshold})"
    )
    audit.derived(
        f"CorrectConfident = {cc} ({cc/n*100:.1f}%)",
        f"(whole_tm >= {threshold}) & (fsr_tm >= {threshold})"
    )

    # IR = SC / (SC + FV)
    ir = sc / (sc + fv) if (sc + fv) > 0 else 1.0
    audit.derived(
        f"IR = SC/(SC+FV) = {sc}/({sc}+{fv}) = {ir:.4f}",
        "SDI Internalization Rate formula"
    )

    # Threshold sensitivity (check ±0.05)
    for alt_t in [threshold - 0.05, threshold + 0.05]:
        alt_fv = int(np.sum((whole_tm >= alt_t) & (fsr_tm < alt_t)))
        alt_sc = int(np.sum((whole_tm < alt_t) & (fsr_tm < alt_t)))
        alt_ir = alt_sc / (alt_sc + alt_fv) if (alt_sc + alt_fv) > 0 else 1.0
        audit.derived(
            f"Sensitivity check (threshold={alt_t:.2f}): FV={alt_fv} ({alt_fv/n*100:.1f}%), IR={alt_ir:.4f}",
            f"Same classification with threshold={alt_t:.2f}"
        )

    return {
        'n': n,
        'threshold': threshold,
        'false_verify': fv,
        'self_catch': sc,
        'over_refusal': orr,
        'correct_confident': cc,
        'fv_rate': fv / n,
        'ir': ir,
        'audit': audit
    }


def print_results(results: dict):
    """Print population-level results."""
    n = results['n']
    t = results['threshold']
    fv = results['false_verify']
    sc = results['self_catch']
    orr = results['over_refusal']
    cc = results['correct_confident']
    ir = results['ir']

    print()
    print("=" * 70)
    print("POPULATION-LEVEL SDI TYPING")
    print(f"  Framework: λ[C ≥ P ≥ E]")
    print(f"  Threshold: TM-score ≥ {t}")
    print(f"  Total predictions: {n:,}")
    print("=" * 70)
    print()
    print(f"  FalseVerify      (whole≥{t}, FSR<{t}):  {fv:>8,}  ({fv/n*100:.1f}%)")
    print(f"  SelfCatch        (whole<{t}, FSR<{t}):  {sc:>8,}  ({sc/n*100:.1f}%)")
    print(f"  OverRefusal      (whole<{t}, FSR≥{t}):  {orr:>8,}  ({orr/n*100:.1f}%)")
    print(f"  CorrectConfident (whole≥{t}, FSR≥{t}):  {cc:>8,}  ({cc/n*100:.1f}%)")
    print()
    print(f"  IR = SC/(SC+FV) = {ir:.4f}")
    print()


if __name__ == "__main__":
    import sys
    if len(sys.argv) < 3:
        print("Usage: python population_typing.py <whole_tm.npy> <fsr_tm.npy> [threshold]")
        sys.exit(1)

    whole_path = sys.argv[1]
    fsr_path = sys.argv[2]
    threshold = float(sys.argv[3]) if len(sys.argv) > 3 else 0.5

    results = run_population_typing(whole_path, fsr_path, threshold)
    print_results(results)
    print(results['audit'].report())
```
