## supplement_v6 for "Confidence Without Verification: Screening pLDDT Unreliability in AlphaFold2 Fold-Switching Predictions": run_analysis.py

```
#!/usr/bin/env python3
"""
SDI-UE Protein Fold Integrity Analyzer
=======================================

Detects confident-but-wrong predictions in AlphaFold fold-switching
benchmarks using the SDI framework (Structural Dependencies of Intelligence).

Framework: λ[C ≥ P ≥ E]
  C (Coherence):   Internal consistency of all claims
  P (Preservation): What can be verified against evidence
  E (Emergence):    Novel claims generated by this analysis

Constraint: No claim may exceed what preservation can verify.
Every output includes an audit trail documenting verification status.

Usage:
  # Population level (requires pre-computed TM-score arrays)
  python run_analysis.py population \\
      --whole-tm tmscores_all.npy \\
      --fsr-tm tmscores_fs_all.npy

  # Per-residue level (requires PDBs + scores)
  python run_analysis.py per-residue \\
      --pred prediction.pdb \\
      --ref1 fold1.pdb --ref1-label "2oug_C alpha" \\
      --ref2 fold2.pdb --ref2-label "6c6s_D beta" \\
      --scores scores.json \\
      --fsr 115-162

  # Full analysis (both)
  python run_analysis.py full \\
      --whole-tm tmscores_all.npy \\
      --fsr-tm tmscores_fs_all.npy \\
      --pred prediction.pdb \\
      --ref1 fold1.pdb --ref1-label "2oug_C alpha" \\
      --ref2 fold2.pdb --ref2-label "6c6s_D beta" \\
      --scores scores.json \\
      --fsr 115-162

Reference:
  SDI v1.1: Canadian Patent Application CA 3294110
  UE v1.9: Unbounded Emergence — operational constraints
  R. Thacker, Vancouver Island University

License: Research use. Contact author for commercial licensing.
"""

import argparse
import json
import os
import sys
from datetime import datetime

from audit_trail import AuditTrail
from population_typing import run_population_typing, print_results as print_pop
from tier1_analysis import run_tier1, parse_fsr_range, print_results as print_t1
from tier2_analysis import run_tier2, print_results as print_t2


def main():
    parser = argparse.ArgumentParser(
        description="SDI-UE Protein Fold Integrity Analyzer",
        formatter_class=argparse.RawDescriptionHelpFormatter,
        epilog="""
Examples:
  python run_analysis.py population --whole-tm tmscores_all.npy --fsr-tm tmscores_fs_all.npy
  python run_analysis.py per-residue --pred pred.pdb --ref1 fold1.pdb --ref2 fold2.pdb --scores scores.json --fsr 115-162
  python run_analysis.py full --whole-tm tm_all.npy --fsr-tm tm_fsr.npy --pred pred.pdb --ref1 f1.pdb --ref2 f2.pdb --scores s.json --fsr 115-162
        """
    )

    parser.add_argument('mode', choices=['population', 'per-residue', 'full'],
                        help="Analysis mode")

    # Population args
    parser.add_argument('--whole-tm', help="Whole-protein TM-score .npy file")
    parser.add_argument('--fsr-tm', help="FSR-specific TM-score .npy file")
    parser.add_argument('--tm-threshold', type=float, default=0.5,
                        help="TM-score threshold (default: 0.5)")

    # Per-residue args
    parser.add_argument('--pred', help="ColabFold prediction PDB")
    parser.add_argument('--ref1', help="Experimental reference PDB, fold 1")
    parser.add_argument('--ref2', help="Experimental reference PDB, fold 2")
    parser.add_argument('--ref1-label', default="fold1", help="Label for fold 1")
    parser.add_argument('--ref2-label', default="fold2", help="Label for fold 2")
    parser.add_argument('--scores', help="ColabFold scores JSON")
    parser.add_argument('--fsr', help="FSR residue range (e.g., 115-162)")

    # Thresholds
    parser.add_argument('--plddt-threshold', type=float, default=70.0,
                        help="pLDDT confident threshold (default: 70.0)")
    parser.add_argument('--pae-threshold', type=float, default=10.0,
                        help="PAE confident threshold (default: 10.0 Å)")
    parser.add_argument('--dist-threshold', type=float, default=3.8,
                        help="CA distance wrong-fold threshold (default: 3.8 Å)")
    parser.add_argument('--pae-scope', choices=['fsr_internal', 'all_columns'],
                        default='fsr_internal',
                        help="PAE computation scope (default: fsr_internal)")

    # Output
    parser.add_argument('--output-dir', default='results',
                        help="Output directory (default: results)")
    parser.add_argument('--quiet', action='store_true',
                        help="Suppress printed output")

    args = parser.parse_args()

    # Validate required args
    if args.mode in ('population', 'full'):
        if not args.whole_tm or not args.fsr_tm:
            parser.error("Population mode requires --whole-tm and --fsr-tm")

    if args.mode in ('per-residue', 'full'):
        missing = []
        if not args.pred: missing.append('--pred')
        if not args.ref1: missing.append('--ref1')
        if not args.ref2: missing.append('--ref2')
        if not args.scores: missing.append('--scores')
        if not args.fsr: missing.append('--fsr')
        if missing:
            parser.error(f"Per-residue mode requires: {', '.join(missing)}")

    # Create output directory
    os.makedirs(args.output_dir, exist_ok=True)

    # Shared audit trail
    audit = AuditTrail()

    # Record run configuration
    audit.verified(
        f"Analysis mode: {args.mode}",
        "CLI argument"
    )
    audit.verified(
        f"Thresholds: TM={args.tm_threshold}, pLDDT={args.plddt_threshold}, "
        f"PAE={args.pae_threshold}, dist={args.dist_threshold}",
        "CLI arguments (defaults or user-specified)"
    )

    results = {}
    timestamp = datetime.now().strftime("%Y%m%d_%H%M%S")

    # === POPULATION LEVEL ===
    if args.mode in ('population', 'full'):
        if not args.quiet:
            print("\n" + "=" * 70)
            print("RUNNING: Population-Level SDI Typing")
            print("=" * 70)

        pop_results = run_population_typing(
            args.whole_tm, args.fsr_tm, args.tm_threshold, audit
        )

        if not args.quiet:
            print_pop(pop_results)

        results['population'] = {
            'n': pop_results['n'],
            'threshold': pop_results['threshold'],
            'false_verify': pop_results['false_verify'],
            'self_catch': pop_results['self_catch'],
            'over_refusal': pop_results['over_refusal'],
            'correct_confident': pop_results['correct_confident'],
            'fv_rate': pop_results['fv_rate'],
            'ir': pop_results['ir']
        }

    # === PER-RESIDUE LEVEL ===
    if args.mode in ('per-residue', 'full'):
        fsr_residues = parse_fsr_range(args.fsr)

        if not args.quiet:
            print("\n" + "=" * 70)
            print("RUNNING: Tier 1 (Internal Metrics Only)")
            print("=" * 70)

        t1_results = run_tier1(
            args.scores, fsr_residues,
            args.pae_threshold, args.plddt_threshold, args.pae_scope, audit
        )

        if not args.quiet:
            print_t1(t1_results)

        if not args.quiet:
            print("\n" + "=" * 70)
            print("RUNNING: Tier 2 (Structural SDI Typing)")
            print("=" * 70)

        t2_results = run_tier2(
            args.pred, args.ref1, args.ref2, args.scores, fsr_residues,
            args.ref1_label, args.ref2_label,
            args.dist_threshold, args.pae_threshold, args.plddt_threshold,
            args.pae_scope, audit
        )

        if not args.quiet:
            print_t2(t2_results)

        results['tier1'] = {
            'n_residues': t1_results['n_residues'],
            'n_confident': t1_results['n_confident'],
            'n_uncertain': t1_results['n_uncertain'],
            'ptm': t1_results['ptm'],
            'thresholds': {
                'plddt': args.plddt_threshold,
                'pae': args.pae_threshold,
                'pae_scope': args.pae_scope
            }
        }

        results['tier2'] = {
            'n_residues': t2_results['n_residues'],
            'predicted_fold': t2_results['predicted_fold'],
            'alternate_fold': t2_results['alternate_fold'],
            'counts': t2_results['counts'],
            'fv_rate': t2_results['fv_rate'],
            'ir': t2_results['ir'],
            'tier1_detects_fv': t2_results['tier1_detects_fv'],
            'tier1_misses_fv': t2_results['tier1_misses_fv'],
            'thresholds': {
                'dist': args.dist_threshold,
                'plddt': args.plddt_threshold,
                'pae': args.pae_threshold,
                'pae_scope': args.pae_scope
            },
            'residues': t2_results['residues']
        }

    # === SAVE RESULTS ===
    results_file = os.path.join(args.output_dir, f"sdi_results_{timestamp}.json")
    with open(results_file, 'w') as f:
        json.dump({
            'framework': 'SDI-UE λ[C ≥ P ≥ E]',
            'version': '1.0',
            'timestamp': datetime.now().isoformat(),
            'configuration': {
                'mode': args.mode,
                'tm_threshold': args.tm_threshold,
                'plddt_threshold': args.plddt_threshold,
                'pae_threshold': args.pae_threshold,
                'dist_threshold': args.dist_threshold,
                'pae_scope': args.pae_scope
            },
            'results': results
        }, f, indent=2, default=str)

    # === SAVE AUDIT TRAIL ===
    audit_text_file = os.path.join(args.output_dir, f"audit_trail_{timestamp}.txt")
    with open(audit_text_file, 'w') as f:
        f.write(audit.report())

    audit_json_file = os.path.join(args.output_dir, f"audit_trail_{timestamp}.json")
    with open(audit_json_file, 'w') as f:
        f.write(audit.to_json())

    if not args.quiet:
        print("\n" + audit.report())
        print(f"\n[FILES]")
        print(f"  Results:     {results_file}")
        print(f"  Audit (txt): {audit_text_file}")
        print(f"  Audit (json): {audit_json_file}")


if __name__ == "__main__":
    main()
```
