## supplement_v6 for "Confidence Without Verification: Screening pLDDT Unreliability in AlphaFold2 Fold-Switching Predictions": sdi_deviation_tracer.py

```
#!/usr/bin/env python3
"""
SDI Deviation Tracer — MAGIC Phase 2
=====================================
Applies SDI (Structural Dependencies of Intelligence) typing to AlphaFold2
predictions on Dr. Lauren Porter's fold-switching protein benchmark.

Detects per-residue E>P violations: where model confidence (pLDDT) exceeds
verifiable structural accuracy (CA distance to experimental reference).

Two-tier analysis:
  Tier 1: PAE-based (no experimental reference required)
  Tier 2: Coordinate-based (requires experimental reference)

Input: Porter AF2 benchmark dataset
Output: Per-protein SDI deviation traces + aggregate statistics

Author: Robin (PI) + Claude (research assistant)
Patent: Canadian Patent Application 3294110 (CPE/SDI methods)
"""

import json
import os
import sys
import csv
import numpy as np
from pathlib import Path
from typing import Dict, List, Tuple, Optional, NamedTuple
from dataclasses import dataclass, field


# ============================================================================
# DATA STRUCTURES
# ============================================================================

@dataclass
class SDIResult:
    """Per-residue SDI typing result"""
    residue_num: int
    residue_name: str
    plddt: float
    ca_distance: float       # Å, from Tier 2 (NaN if Tier 1 only)
    cross_pae: float         # mean PAE to other domain (Tier 1)
    local_pae: float         # mean PAE to ±5 neighbors
    sdi_type: str            # FalseVerify, SelfCatch, OverRefusal, Correct
    in_fsr: bool             # whether residue is in fold-switching region

@dataclass
class ProteinSDITrace:
    """Complete SDI trace for one protein"""
    pdb1: str                # ground state PDB ID
    pdb2: str                # excited state PDB ID
    fsr_range: str           # e.g., "115-162"
    n_residues: int
    n_fsr_residues: int
    tier: int                # 1 or 2
    residues: List[SDIResult]
    
    # Aggregate metrics (whole protein)
    false_verify_count: int = 0
    self_catch_count: int = 0
    over_refusal_count: int = 0
    correct_confident_count: int = 0
    ir_whole: float = 0.0
    
    # Aggregate metrics (FSR only)
    fsr_false_verify: int = 0
    fsr_self_catch: int = 0
    fsr_over_refusal: int = 0
    fsr_correct_confident: int = 0
    ir_fsr: float = 0.0
    
    # Global scores
    mean_plddt_whole: float = 0.0
    mean_plddt_fsr: float = 0.0
    ptm: float = 0.0
    nonfsr_rmsd: float = 0.0
    fsr_rmsd: float = 0.0


# ============================================================================
# PDB PARSING
# ============================================================================

def parse_ca_atoms(pdb_path: str) -> Tuple[np.ndarray, np.ndarray, List[int], List[str]]:
    """
    Extract CA atom coordinates, B-factors, residue numbers and names from PDB.
    
    Returns:
        coords: (N, 3) array of CA coordinates
        bfactors: (N,) array of B-factors (pLDDT in ColabFold PDBs)
        resnums: list of residue numbers
        resnames: list of 3-letter residue names
    """
    coords, bfactors, resnums, resnames = [], [], [], []
    with open(pdb_path) as f:
        for line in f:
            if line.startswith('ATOM') and line[12:16].strip() == 'CA':
                coords.append([
                    float(line[30:38]),
                    float(line[38:46]),
                    float(line[46:54])
                ])
                bfactors.append(float(line[60:66]))
                resnums.append(int(line[22:26]))
                resnames.append(line[17:20].strip())
    return np.array(coords), np.array(bfactors), resnums, resnames


def get_sequence(resnames: List[str]) -> str:
    """Convert 3-letter residue names to 1-letter sequence"""
    aa3to1 = {
        'ALA':'A','ARG':'R','ASN':'N','ASP':'D','CYS':'C','GLN':'Q','GLU':'E',
        'GLY':'G','HIS':'H','ILE':'I','LEU':'L','LYS':'K','MET':'M','PHE':'F',
        'PRO':'P','SER':'S','THR':'T','TRP':'W','TYR':'Y','VAL':'V'
    }
    return ''.join(aa3to1.get(r, 'X') for r in resnames)


# ============================================================================
# STRUCTURAL ALIGNMENT
# ============================================================================

def kabsch_align(P: np.ndarray, Q: np.ndarray) -> Tuple[np.ndarray, np.ndarray, np.ndarray]:
    """
    Kabsch algorithm: find optimal rotation to align P onto Q.
    
    Args:
        P: (N, 3) coordinates to be rotated
        Q: (N, 3) target coordinates
    
    Returns:
        R: (3, 3) rotation matrix
        p_mean: centroid of P
        q_mean: centroid of Q
    """
    p_mean = P.mean(axis=0)
    q_mean = Q.mean(axis=0)
    H = (P - p_mean).T @ (Q - q_mean)
    U, S, Vt = np.linalg.svd(H)
    d = np.linalg.det(Vt.T @ U.T)
    sign = np.eye(3)
    sign[2, 2] = np.sign(d)
    R = Vt.T @ sign @ U.T
    return R, p_mean, q_mean


def superpose_and_measure(
    pred_coords: np.ndarray,
    ref_coords: np.ndarray,
    align_mask: np.ndarray
) -> Tuple[np.ndarray, float, float]:
    """
    Superpose prediction onto reference using subset of residues (align_mask),
    then measure per-residue CA distance for ALL residues.
    
    Args:
        pred_coords: (N, 3) predicted coordinates
        ref_coords: (N, 3) reference coordinates  
        align_mask: (N,) boolean mask — True for residues used in alignment
    
    Returns:
        distances: (N,) per-residue CA distances after superposition
        align_rmsd: RMSD of alignment region
        total_rmsd: RMSD of all residues
    """
    R, p_mean, q_mean = kabsch_align(
        pred_coords[align_mask],
        ref_coords[align_mask]
    )
    pred_sup = ((pred_coords - p_mean) @ R.T) + q_mean
    distances = np.sqrt(np.sum((pred_sup - ref_coords)**2, axis=1))
    align_rmsd = np.sqrt(np.mean(distances[align_mask]**2))
    total_rmsd = np.sqrt(np.mean(distances**2))
    return distances, align_rmsd, total_rmsd


# ============================================================================
# SEQUENCE ALIGNMENT (for offset detection)
# ============================================================================

def find_sequence_offset(pred_names: List[str], ref_names: List[str]) -> int:
    """
    Find the offset between prediction and reference sequences.
    Handles initiator methionine and other N-terminal differences.
    
    Returns:
        offset: number of residues to skip at start of prediction
                to align with reference. pred[offset:] aligns to ref[:]
    """
    pred_seq = get_sequence(pred_names)
    ref_seq = get_sequence(ref_names)
    
    best_offset = 0
    best_matches = 0
    max_check = min(10, len(pred_seq) - len(ref_seq) + 1)
    
    for offset in range(max(0, -5), max_check + 1):
        if offset >= 0:
            p = pred_seq[offset:offset + len(ref_seq)]
            r = ref_seq
        else:
            p = pred_seq
            r = ref_seq[-offset:-offset + len(pred_seq)]
        
        compare_len = min(len(p), len(r))
        if compare_len == 0:
            continue
        matches = sum(1 for a, b in zip(p[:compare_len], r[:compare_len]) if a == b)
        if matches > best_matches:
            best_matches = matches
            best_offset = offset
    
    return best_offset


# ============================================================================
# FSR RANGE PARSING
# ============================================================================

def parse_fsr_ranges(range_file: str) -> Dict[Tuple[str, str], dict]:
    """Parse fold-switching region definitions from range_fs_pairs_all.txt"""
    pairs = {}
    with open(range_file) as f:
        for line in f:
            line = line.strip().rstrip(',')
            if line.startswith('#') or not line:
                continue
            parts = [p.strip() for p in line.split(',')]
            pdb1 = parts[0]
            pdb2 = parts[1]
            pairs[(pdb1, pdb2)] = {
                'range1': parts[2],
                'range2': parts[3],
                'pred1': parts[4] if len(parts) > 4 else parts[2],
                'pred2': parts[5] if len(parts) > 5 else parts[3],
            }
    return pairs


def parse_range_string(range_str: str) -> Tuple[int, int]:
    """Parse '115-162' into (115, 162)"""
    parts = range_str.split('-')
    return int(parts[0]), int(parts[1])


# ============================================================================
# SDI TYPING
# ============================================================================

def classify_residue(
    plddt: float,
    error_metric: float,
    plddt_threshold: float = 70.0,
    error_threshold: float = 3.8
) -> str:
    """
    Classify a residue into SDI type.
    
    Args:
        plddt: per-residue pLDDT score
        error_metric: CA distance (Tier 2) or cross-domain PAE (Tier 1)
        plddt_threshold: confidence threshold (default 70)
        error_threshold: error threshold (default 3.8 Å for Tier 2, 10.0 for Tier 1)
    
    Returns:
        SDI type string
    """
    confident = plddt >= plddt_threshold
    wrong = error_metric > error_threshold
    
    if confident and wrong:
        return 'FalseVerify'
    elif not confident and wrong:
        return 'SelfCatch'
    elif not confident and not wrong:
        return 'OverRefusal'
    else:
        return 'Correct'


def compute_ir(self_catch: int, false_verify: int) -> float:
    """Compute Internalization Rate"""
    total = self_catch + false_verify
    return self_catch / total if total > 0 else float('nan')


# ============================================================================
# TIER 2: COORDINATE-BASED ANALYSIS
# ============================================================================

def run_tier2(
    pred_pdb: str,
    ref_pdb: str,
    score_json: str,
    fsr_start: int,
    fsr_end: int,
    pdb1: str,
    pdb2: str,
    plddt_threshold: float = 70.0,
    dist_threshold: float = 3.8
) -> ProteinSDITrace:
    """
    Run Tier 2 (coordinate-based) SDI analysis on a single protein.
    
    Args:
        pred_pdb: path to ColabFold relaxed prediction PDB
        ref_pdb: path to experimental reference PDB
        score_json: path to ColabFold scores JSON (for PAE data)
        fsr_start: first residue of FSR (1-indexed, prediction numbering)
        fsr_end: last residue of FSR (1-indexed, prediction numbering)
        pdb1: ground state PDB ID
        pdb2: excited state PDB ID
        plddt_threshold: confidence cutoff
        dist_threshold: structural error cutoff (Å)
    
    Returns:
        ProteinSDITrace with complete results
    """
    # Load structures
    pred_c, pred_b, pred_r, pred_n = parse_ca_atoms(pred_pdb)
    ref_c, ref_b, ref_r, ref_n = parse_ca_atoms(ref_pdb)
    
    # Detect and apply sequence offset
    offset = find_sequence_offset(pred_n, ref_n)
    n_common = min(len(pred_r) - offset, len(ref_r))
    
    pred_c_a = pred_c[offset:offset + n_common]
    pred_b_a = pred_b[offset:offset + n_common]
    pred_r_a = pred_r[offset:offset + n_common]
    pred_n_a = pred_n[offset:offset + n_common]
    ref_c_a = ref_c[:n_common]
    ref_r_a = ref_r[:n_common]
    ref_n_a = ref_n[:n_common]
    
    # Verify sequence match
    pred_seq = get_sequence(pred_n_a)
    ref_seq = get_sequence(ref_n_a)
    if pred_seq != ref_seq:
        print(f"WARNING: Sequence mismatch for {pdb1}/{pdb2} after offset {offset}")
        print(f"  Pred: {pred_seq[:50]}...")
        print(f"  Ref:  {ref_seq[:50]}...")
    
    # Define FSR mask (adjusted for offset)
    # pred_r_a contains prediction residue numbers after offset
    fsr_mask = np.array([(r >= fsr_start and r <= min(fsr_end, pred_r_a[-1])) 
                         for r in pred_r_a])
    non_fsr_mask = ~fsr_mask
    
    # Load PAE data if available
    pae_data = None
    ptm = 0.0
    if score_json and os.path.exists(score_json):
        with open(score_json) as f:
            jdata = json.load(f)
        pae_data = np.array(jdata.get('pae', []))
        ptm = jdata.get('ptm', 0.0)
    
    # Superpose on non-FSR region, measure all
    if non_fsr_mask.sum() < 3:
        print(f"WARNING: Too few non-FSR residues for alignment in {pdb1}")
        distances = np.full(n_common, np.nan)
        nonfsr_rmsd = np.nan
        fsr_rmsd = np.nan
    else:
        distances, nonfsr_rmsd, _ = superpose_and_measure(
            pred_c_a, ref_c_a, non_fsr_mask
        )
        fsr_rmsd = np.sqrt(np.mean(distances[fsr_mask]**2)) if fsr_mask.sum() > 0 else np.nan
    
    # Build per-residue results
    residues = []
    for i in range(n_common):
        pred_res = pred_r_a[i]
        in_fsr = fsr_mask[i]
        
        # Cross-domain PAE (if available)
        cross_pae = np.nan
        local_pae = np.nan
        if pae_data is not None and len(pae_data) > pred_res - 1:
            idx = pred_res - 1  # 0-indexed into full pae matrix
            if idx < len(pae_data):
                # Cross-PAE: mean to non-FSR residues
                nonfsr_pae_idx = [j for j in range(len(pae_data)) 
                                  if not (j + 1 >= fsr_start and j + 1 <= fsr_end)]
                if nonfsr_pae_idx:
                    cross_pae = pae_data[idx, nonfsr_pae_idx].mean()
                # Local PAE: ±5 neighbors
                local_idx = [j for j in range(max(0, idx-5), min(len(pae_data), idx+6)) if j != idx]
                if local_idx:
                    local_pae = pae_data[idx, local_idx].mean()
        
        sdi_type = classify_residue(
            pred_b_a[i], distances[i], plddt_threshold, dist_threshold
        )
        
        residues.append(SDIResult(
            residue_num=pred_res,
            residue_name=pred_n_a[i],
            plddt=pred_b_a[i],
            ca_distance=distances[i],
            cross_pae=cross_pae,
            local_pae=local_pae,
            sdi_type=sdi_type,
            in_fsr=in_fsr
        ))
    
    # Aggregate
    fv = sum(1 for r in residues if r.sdi_type == 'FalseVerify')
    sc = sum(1 for r in residues if r.sdi_type == 'SelfCatch')
    orf = sum(1 for r in residues if r.sdi_type == 'OverRefusal')
    cc = sum(1 for r in residues if r.sdi_type == 'Correct')
    
    fsr_res = [r for r in residues if r.in_fsr]
    fv_f = sum(1 for r in fsr_res if r.sdi_type == 'FalseVerify')
    sc_f = sum(1 for r in fsr_res if r.sdi_type == 'SelfCatch')
    orf_f = sum(1 for r in fsr_res if r.sdi_type == 'OverRefusal')
    cc_f = sum(1 for r in fsr_res if r.sdi_type == 'Correct')
    
    plddt_all = np.array([r.plddt for r in residues])
    plddt_fsr = np.array([r.plddt for r in fsr_res]) if fsr_res else np.array([])
    
    return ProteinSDITrace(
        pdb1=pdb1,
        pdb2=pdb2,
        fsr_range=f"{fsr_start}-{fsr_end}",
        n_residues=n_common,
        n_fsr_residues=len(fsr_res),
        tier=2,
        residues=residues,
        false_verify_count=fv,
        self_catch_count=sc,
        over_refusal_count=orf,
        correct_confident_count=cc,
        ir_whole=compute_ir(sc, fv),
        fsr_false_verify=fv_f,
        fsr_self_catch=sc_f,
        fsr_over_refusal=orf_f,
        fsr_correct_confident=cc_f,
        ir_fsr=compute_ir(sc_f, fv_f),
        mean_plddt_whole=float(plddt_all.mean()),
        mean_plddt_fsr=float(plddt_fsr.mean()) if len(plddt_fsr) > 0 else 0.0,
        ptm=ptm,
        nonfsr_rmsd=nonfsr_rmsd,
        fsr_rmsd=fsr_rmsd
    )


# ============================================================================
# TIER 1: PAE-BASED ANALYSIS (no experimental reference)
# ============================================================================

def run_tier1(
    score_json: str,
    fsr_start: int,
    fsr_end: int,
    pdb1: str,
    pdb2: str,
    plddt_threshold: float = 70.0,
    pae_threshold: float = 10.0
) -> ProteinSDITrace:
    """
    Run Tier 1 (PAE-based) SDI analysis. No experimental reference needed.
    Uses cross-domain PAE as proxy for structural accuracy.
    
    WARNING: Tier 1 MISSES confident-wrong predictions where both pLDDT
    and PAE agree on the wrong structure. Use Tier 2 when experimental
    references are available.
    """
    with open(score_json) as f:
        jdata = json.load(f)
    
    plddt = np.array(jdata['plddt'])
    pae = np.array(jdata['pae'])
    ptm = jdata.get('ptm', 0.0)
    n = len(plddt)
    
    fsr_idx = [i for i in range(n) if (i + 1) >= fsr_start and (i + 1) <= fsr_end]
    non_fsr_idx = [i for i in range(n) if not ((i + 1) >= fsr_start and (i + 1) <= fsr_end)]
    
    residues = []
    for i in range(n):
        in_fsr = (i + 1) >= fsr_start and (i + 1) <= fsr_end
        
        # Cross-domain PAE
        if non_fsr_idx:
            cross_pae = pae[i, non_fsr_idx].mean()
        else:
            cross_pae = pae[i].mean()
        
        # Local PAE
        local_idx = [j for j in range(max(0, i-5), min(n, i+6)) if j != i]
        local_pae = pae[i, local_idx].mean() if local_idx else 0.0
        
        sdi_type = classify_residue(
            plddt[i], cross_pae, plddt_threshold, pae_threshold
        )
        
        residues.append(SDIResult(
            residue_num=i + 1,
            residue_name='',  # not available from JSON
            plddt=float(plddt[i]),
            ca_distance=float('nan'),
            cross_pae=float(cross_pae),
            local_pae=float(local_pae),
            sdi_type=sdi_type,
            in_fsr=in_fsr
        ))
    
    # Aggregate (same as Tier 2)
    fv = sum(1 for r in residues if r.sdi_type == 'FalseVerify')
    sc = sum(1 for r in residues if r.sdi_type == 'SelfCatch')
    orf = sum(1 for r in residues if r.sdi_type == 'OverRefusal')
    cc = sum(1 for r in residues if r.sdi_type == 'Correct')
    
    fsr_res = [r for r in residues if r.in_fsr]
    fv_f = sum(1 for r in fsr_res if r.sdi_type == 'FalseVerify')
    sc_f = sum(1 for r in fsr_res if r.sdi_type == 'SelfCatch')
    orf_f = sum(1 for r in fsr_res if r.sdi_type == 'OverRefusal')
    cc_f = sum(1 for r in fsr_res if r.sdi_type == 'Correct')
    
    return ProteinSDITrace(
        pdb1=pdb1, pdb2=pdb2,
        fsr_range=f"{fsr_start}-{fsr_end}",
        n_residues=n,
        n_fsr_residues=len(fsr_res),
        tier=1,
        residues=residues,
        false_verify_count=fv, self_catch_count=sc,
        over_refusal_count=orf, correct_confident_count=cc,
        ir_whole=compute_ir(sc, fv),
        fsr_false_verify=fv_f, fsr_self_catch=sc_f,
        fsr_over_refusal=orf_f, fsr_correct_confident=cc_f,
        ir_fsr=compute_ir(sc_f, fv_f),
        mean_plddt_whole=float(plddt.mean()),
        mean_plddt_fsr=float(plddt[fsr_idx].mean()) if fsr_idx else 0.0,
        ptm=ptm,
        nonfsr_rmsd=float('nan'),
        fsr_rmsd=float('nan')
    )


# ============================================================================
# OUTPUT FORMATTING
# ============================================================================

def print_trace(trace: ProteinSDITrace, verbose: bool = True):
    """Print SDI trace to console"""
    print(f"\n{'='*70}")
    print(f"SDI Deviation Trace: {trace.pdb1} <-> {trace.pdb2}")
    print(f"Tier: {trace.tier}, FSR: {trace.fsr_range}")
    print(f"{'='*70}")
    
    print(f"\nGlobal: pTM={trace.ptm:.3f}, mean pLDDT={trace.mean_plddt_whole:.1f}")
    if trace.tier == 2:
        print(f"Non-FSR RMSD: {trace.nonfsr_rmsd:.2f} Å, FSR RMSD: {trace.fsr_rmsd:.2f} Å")
    
    print(f"\nWhole protein (n={trace.n_residues}):")
    n = trace.n_residues
    print(f"  FalseVerify:      {trace.false_verify_count:3d} ({100*trace.false_verify_count/n:5.1f}%)")
    print(f"  SelfCatch:        {trace.self_catch_count:3d} ({100*trace.self_catch_count/n:5.1f}%)")
    print(f"  OverRefusal:      {trace.over_refusal_count:3d} ({100*trace.over_refusal_count/n:5.1f}%)")
    print(f"  CorrectConfident: {trace.correct_confident_count:3d} ({100*trace.correct_confident_count/n:5.1f}%)")
    print(f"  IR: {trace.ir_whole:.4f}")
    
    nf = trace.n_fsr_residues
    if nf > 0:
        print(f"\nFSR only (n={nf}, residues {trace.fsr_range}):")
        print(f"  FalseVerify:      {trace.fsr_false_verify:3d} ({100*trace.fsr_false_verify/nf:5.1f}%)")
        print(f"  SelfCatch:        {trace.fsr_self_catch:3d} ({100*trace.fsr_self_catch/nf:5.1f}%)")
        print(f"  OverRefusal:      {trace.fsr_over_refusal:3d} ({100*trace.fsr_over_refusal/nf:5.1f}%)")
        print(f"  CorrectConfident: {trace.fsr_correct_confident:3d} ({100*trace.fsr_correct_confident/nf:5.1f}%)")
        print(f"  IR: {trace.ir_fsr:.4f}")
    
    if verbose:
        print(f"\nPer-residue detail (FSR):")
        header = f"{'Res':>5s}  {'Name':>4s}  {'pLDDT':>6s}"
        if trace.tier == 2:
            header += f"  {'Dist':>7s}"
        header += f"  {'CrossPAE':>8s}  {'Type'}"
        print(header)
        
        for r in trace.residues:
            if r.in_fsr:
                line = f"{r.residue_num:5d}  {r.residue_name:>4s}  {r.plddt:6.1f}"
                if trace.tier == 2:
                    line += f"  {r.ca_distance:7.2f}"
                pae_str = f"{r.cross_pae:8.2f}" if not np.isnan(r.cross_pae) else f"{'N/A':>8s}"
                line += f"  {pae_str}  {r.sdi_type}"
                print(line)


def write_csv(trace: ProteinSDITrace, output_path: str):
    """Write per-residue results to CSV"""
    with open(output_path, 'w', newline='') as f:
        writer = csv.writer(f)
        writer.writerow([
            'residue_num', 'residue_name', 'plddt', 'ca_distance',
            'cross_pae', 'local_pae', 'sdi_type', 'in_fsr',
            'pdb1', 'pdb2', 'tier'
        ])
        for r in trace.residues:
            writer.writerow([
                r.residue_num, r.residue_name, f"{r.plddt:.2f}",
                f"{r.ca_distance:.2f}" if not np.isnan(r.ca_distance) else '',
                f"{r.cross_pae:.2f}" if not np.isnan(r.cross_pae) else '',
                f"{r.local_pae:.2f}" if not np.isnan(r.local_pae) else '',
                r.sdi_type, r.in_fsr,
                trace.pdb1, trace.pdb2, trace.tier
            ])


def write_summary_csv(traces: List[ProteinSDITrace], output_path: str):
    """Write aggregate summary across all proteins"""
    with open(output_path, 'w', newline='') as f:
        writer = csv.writer(f)
        writer.writerow([
            'pdb1', 'pdb2', 'tier', 'fsr_range', 'n_residues', 'n_fsr',
            'ptm', 'mean_plddt', 'mean_plddt_fsr', 'nonfsr_rmsd', 'fsr_rmsd',
            'fv_whole', 'sc_whole', 'or_whole', 'cc_whole', 'ir_whole',
            'fv_fsr', 'sc_fsr', 'or_fsr', 'cc_fsr', 'ir_fsr'
        ])
        for t in traces:
            writer.writerow([
                t.pdb1, t.pdb2, t.tier, t.fsr_range, t.n_residues, t.n_fsr_residues,
                f"{t.ptm:.3f}", f"{t.mean_plddt_whole:.1f}", f"{t.mean_plddt_fsr:.1f}",
                f"{t.nonfsr_rmsd:.2f}" if not np.isnan(t.nonfsr_rmsd) else '',
                f"{t.fsr_rmsd:.2f}" if not np.isnan(t.fsr_rmsd) else '',
                t.false_verify_count, t.self_catch_count,
                t.over_refusal_count, t.correct_confident_count,
                f"{t.ir_whole:.4f}" if not np.isnan(t.ir_whole) else '',
                t.fsr_false_verify, t.fsr_self_catch,
                t.fsr_over_refusal, t.fsr_correct_confident,
                f"{t.ir_fsr:.4f}" if not np.isnan(t.ir_fsr) else ''
            ])


# ============================================================================
# BATCH RUNNER
# ============================================================================

def run_benchmark(
    data_dir: str,
    range_file: str,
    output_dir: str,
    plddt_threshold: float = 70.0,
    dist_threshold: float = 3.8,
    pae_threshold: float = 10.0,
    verbose: bool = False
) -> List[ProteinSDITrace]:
    """
    Run SDI analysis across the full Porter benchmark.
    
    Attempts Tier 2 for each protein where files exist,
    falls back to Tier 1 (PAE-only) otherwise.
    
    Expected directory structure under data_dir:
        porter_AF2_benchmark/
            AF2Rank/final2_and_debug2_inputs/{pdb_id}.pdb  (experimental)
            data/{pdb_id}/*/...relaxed_rank_001_*.pdb       (prediction)
            data/{pdb_id}/*_scores_rank_001_*.json          (scores)
    """
    fsr_pairs = parse_fsr_ranges(range_file)
    porter_dir = os.path.join(data_dir, 'porter_AF2_benchmark')
    
    os.makedirs(output_dir, exist_ok=True)
    traces_dir = os.path.join(output_dir, 'deviation_traces')
    os.makedirs(traces_dir, exist_ok=True)
    
    traces = []
    
    for (pdb1, pdb2), ranges in fsr_pairs.items():
        pred_start, pred_end = parse_range_string(ranges['pred1'])
        
        # Look for experimental reference
        ref_pdb = os.path.join(porter_dir, 'AF2Rank', 'final2_and_debug2_inputs', f'{pdb1}.pdb')
        
        # Look for ColabFold prediction and scores
        data_subdir = os.path.join(porter_dir, 'data', pdb1)
        pred_pdb = None
        score_json = None
        
        if os.path.isdir(data_subdir):
            for subdir in os.listdir(data_subdir):
                subpath = os.path.join(data_subdir, subdir)
                if os.path.isdir(subpath):
                    for f in os.listdir(subpath):
                        if 'relaxed_rank_001' in f and f.endswith('.pdb'):
                            pred_pdb = os.path.join(subpath, f)
                        if 'scores_rank_001' in f and f.endswith('.json'):
                            score_json = os.path.join(subpath, f)
        
        # Decide tier
        if pred_pdb and ref_pdb and os.path.exists(ref_pdb):
            try:
                trace = run_tier2(
                    pred_pdb, ref_pdb, score_json,
                    pred_start, pred_end, pdb1, pdb2,
                    plddt_threshold, dist_threshold
                )
                if verbose:
                    print_trace(trace, verbose=False)
                traces.append(trace)
                write_csv(trace, os.path.join(traces_dir, f'{pdb1}_{pdb2}_tier2.csv'))
                continue
            except Exception as e:
                print(f"Tier 2 failed for {pdb1}/{pdb2}: {e}")
        
        # Fall back to Tier 1
        if score_json and os.path.exists(score_json):
            try:
                trace = run_tier1(
                    score_json, pred_start, pred_end,
                    pdb1, pdb2, plddt_threshold, pae_threshold
                )
                if verbose:
                    print_trace(trace, verbose=False)
                traces.append(trace)
                write_csv(trace, os.path.join(traces_dir, f'{pdb1}_{pdb2}_tier1.csv'))
                continue
            except Exception as e:
                print(f"Tier 1 failed for {pdb1}/{pdb2}: {e}")
        
        print(f"SKIP: No data found for {pdb1}/{pdb2}")
    
    # Write summary
    if traces:
        write_summary_csv(traces, os.path.join(output_dir, 'summary', 'sdi_benchmark_summary.csv'))
    
    return traces


# ============================================================================
# SENSITIVITY ANALYSIS
# ============================================================================

def threshold_sensitivity(
    trace: ProteinSDITrace,
    plddt_thresholds: List[float] = [50, 60, 70, 80, 90],
    dist_thresholds: List[float] = [2.0, 3.0, 3.8, 5.0, 8.0]
) -> List[dict]:
    """
    Re-classify residues across threshold combinations.
    Shows how sensitive results are to threshold choice.
    """
    results = []
    for pt in plddt_thresholds:
        for dt in dist_thresholds:
            fsr_res = [r for r in trace.residues if r.in_fsr]
            fv = sum(1 for r in fsr_res 
                     if r.plddt >= pt and r.ca_distance > dt)
            sc = sum(1 for r in fsr_res 
                     if r.plddt < pt and r.ca_distance > dt)
            results.append({
                'plddt_thresh': pt,
                'dist_thresh': dt,
                'fv': fv,
                'sc': sc,
                'ir': compute_ir(sc, fv),
                'n_fsr': len(fsr_res)
            })
    return results


# ============================================================================
# MAIN
# ============================================================================

if __name__ == '__main__':
    import argparse
    
    parser = argparse.ArgumentParser(description='SDI Deviation Tracer for protein structure predictions')
    parser.add_argument('--data-dir', required=True, help='Path to MAGIC-Protein-Fold/data/')
    parser.add_argument('--range-file', required=True, help='Path to range_fs_pairs_all.txt')
    parser.add_argument('--output-dir', required=True, help='Path to results/ directory')
    parser.add_argument('--plddt-threshold', type=float, default=70.0)
    parser.add_argument('--dist-threshold', type=float, default=3.8)
    parser.add_argument('--pae-threshold', type=float, default=10.0)
    parser.add_argument('--verbose', action='store_true')
    parser.add_argument('--single', nargs=2, metavar=('PDB1', 'PDB2'),
                        help='Run on single protein pair')
    
    args = parser.parse_args()
    
    if args.single:
        # Single protein mode (for testing)
        pdb1, pdb2 = args.single
        fsr_pairs = parse_fsr_ranges(args.range_file)
        key = (pdb1, pdb2)
        if key not in fsr_pairs:
            key = (pdb2, pdb1)
        if key not in fsr_pairs:
            print(f"ERROR: pair {pdb1}/{pdb2} not found in range file")
            sys.exit(1)
        
        ranges = fsr_pairs[key]
        pred_start, pred_end = parse_range_string(ranges['pred1'])
        
        # Find files (simplified for single mode)
        porter_dir = os.path.join(args.data_dir, 'porter_AF2_benchmark')
        ref_pdb = os.path.join(porter_dir, 'AF2Rank', 'final2_and_debug2_inputs', f'{key[0]}.pdb')
        
        # Search for prediction files
        data_subdir = os.path.join(porter_dir, 'data', key[0])
        pred_pdb = score_json = None
        if os.path.isdir(data_subdir):
            for subdir in os.listdir(data_subdir):
                subpath = os.path.join(data_subdir, subdir)
                if os.path.isdir(subpath):
                    for f in os.listdir(subpath):
                        if 'relaxed_rank_001' in f and f.endswith('.pdb'):
                            pred_pdb = os.path.join(subpath, f)
                        if 'scores_rank_001' in f and f.endswith('.json'):
                            score_json = os.path.join(subpath, f)
        
        if pred_pdb and os.path.exists(ref_pdb):
            trace = run_tier2(pred_pdb, ref_pdb, score_json,
                            pred_start, pred_end, key[0], key[1],
                            args.plddt_threshold, args.dist_threshold)
        elif score_json:
            trace = run_tier1(score_json, pred_start, pred_end,
                            key[0], key[1], args.plddt_threshold, args.pae_threshold)
        else:
            print(f"No data files found for {key[0]}")
            sys.exit(1)
        
        print_trace(trace, verbose=True)
        
        # Sensitivity analysis
        if trace.tier == 2:
            print(f"\n{'='*70}")
            print("Threshold sensitivity (FSR only):")
            print(f"{'pLDDT':>6s}  {'Dist':>5s}  {'FV':>4s}  {'SC':>4s}  {'IR':>6s}")
            for r in threshold_sensitivity(trace):
                ir_str = f"{r['ir']:.3f}" if not np.isnan(r['ir']) else 'N/A'
                print(f"{r['plddt_thresh']:6.0f}  {r['dist_thresh']:5.1f}  {r['fv']:4d}  {r['sc']:4d}  {ir_str:>6s}")
    
    else:
        # Full benchmark mode
        traces = run_benchmark(
            args.data_dir, args.range_file, args.output_dir,
            args.plddt_threshold, args.dist_threshold,
            args.pae_threshold, args.verbose
        )
        
        print(f"\n{'='*70}")
        print(f"BENCHMARK COMPLETE: {len(traces)} proteins analyzed")
        
        tier2 = [t for t in traces if t.tier == 2]
        tier1 = [t for t in traces if t.tier == 1]
        print(f"  Tier 2 (coordinate-based): {len(tier2)}")
        print(f"  Tier 1 (PAE-only): {len(tier1)}")
        
        if tier2:
            all_fv = sum(t.fsr_false_verify for t in tier2)
            all_sc = sum(t.fsr_self_catch for t in tier2)
            all_n = sum(t.n_fsr_residues for t in tier2)
            ir = compute_ir(all_sc, all_fv)
            print(f"\n  Tier 2 aggregate FSR results:")
            print(f"    Total FSR residues: {all_n}")
            print(f"    FalseVerify: {all_fv} ({100*all_fv/all_n:.1f}%)")
            print(f"    SelfCatch: {all_sc} ({100*all_sc/all_n:.1f}%)")
            print(f"    IR: {ir:.4f}")
```
