## supplement_v6 for "Confidence Without Verification: Screening pLDDT Unreliability in AlphaFold2 Fold-Switching Predictions": tier1_analysis.py

```
"""
Tier 1 Analysis: Internal Metrics Only

Classifies FSR residues as confident or uncertain using ONLY
AlphaFold's internal quality metrics (pLDDT, PAE).

This is the "self-verification" layer — what AF2 thinks about
its own predictions. Tier 1 CANNOT detect FalseVerify because
it has no external reference. Its purpose is to establish the
baseline: what would you catch using only internal metrics?

The gap between Tier 1 and Tier 2 is the measurement.
"""

import numpy as np
import json
from audit_trail import AuditTrail


def parse_fsr_range(range_str: str) -> list:
    """Parse '115-162' into list of 1-indexed residue numbers."""
    start, end = map(int, range_str.split('-'))
    return list(range(start, end + 1))


def run_tier1(
    scores_json_path: str,
    fsr_residues: list,
    pae_threshold: float = 10.0,
    plddt_threshold: float = 70.0,
    pae_scope: str = "fsr_internal",
    audit: AuditTrail = None
) -> dict:
    """
    Tier 1: Classify FSR residues using internal metrics only.

    Args:
        scores_json_path: Path to ColabFold scores JSON
        fsr_residues: List of 1-indexed residue numbers in FSR
        pae_threshold: Mean PAE below this = confident (default 10.0 Å)
        plddt_threshold: pLDDT above this = confident (default 70.0)
        pae_scope: How to compute per-residue PAE:
            "fsr_internal" — mean PAE within FSR block only
            "all_columns"  — mean PAE against all residues
        audit: AuditTrail instance

    Returns:
        dict with per-residue metrics and classification
    """
    if audit is None:
        audit = AuditTrail()

    # Load scores
    with open(scores_json_path) as f:
        scores = json.load(f)

    plddt = np.array(scores['plddt'])
    pae_matrix = np.array(scores['pae'])
    ptm = scores.get('ptm', None)

    audit.verified(
        f"Score JSON loaded: {len(plddt)} residues, pTM={ptm}",
        f"json.load({scores_json_path})"
    )
    audit.verified(
        f"PAE matrix shape: {pae_matrix.shape}",
        "scores['pae']"
    )

    # Convert 1-indexed residues to 0-indexed
    fsr_0idx = [r - 1 for r in fsr_residues]

    # Check which FSR residues exist in the score arrays
    valid_fsr = [r for r in fsr_0idx if r < len(plddt)]
    if len(valid_fsr) < len(fsr_0idx):
        missing = len(fsr_0idx) - len(valid_fsr)
        audit.flagged(
            f"{missing} FSR residues out of range of score array (len={len(plddt)})",
            "Index bounds check",
            "Some FSR residues not covered by ColabFold prediction"
        )

    # Compute per-residue PAE
    if pae_scope == "fsr_internal":
        pae_block = pae_matrix[np.ix_(valid_fsr, valid_fsr)]
        pae_per_res = np.mean(pae_block, axis=1)
        audit.derived(
            f"Per-residue PAE computed: scope=FSR-internal ({len(valid_fsr)}x{len(valid_fsr)} block)",
            f"mean(PAE[fsr, fsr], axis=1)"
        )
    elif pae_scope == "all_columns":
        pae_per_res = np.mean(pae_matrix[valid_fsr, :], axis=1)
        audit.derived(
            f"Per-residue PAE computed: scope=all-columns ({len(valid_fsr)}x{pae_matrix.shape[1]})",
            f"mean(PAE[fsr, :], axis=1)"
        )
    else:
        raise ValueError(f"Unknown pae_scope: {pae_scope}")

    # Classification
    residue_data = []
    n_confident = 0
    n_uncertain = 0

    for i, idx_0 in enumerate(valid_fsr):
        resnum = idx_0 + 1  # back to 1-indexed
        p = float(plddt[idx_0])
        pae_val = float(pae_per_res[i])

        confident = (p >= plddt_threshold) and (pae_val < pae_threshold)
        if confident:
            n_confident += 1
        else:
            n_uncertain += 1

        residue_data.append({
            'resnum': resnum,
            'plddt': p,
            'pae': pae_val,
            'confident': confident,
            'tier1_class': 'CONFIDENT' if confident else 'uncertain'
        })

    n_total = len(valid_fsr)
    audit.derived(
        f"Tier 1 classification: {n_confident} confident, {n_uncertain} uncertain "
        f"(of {n_total} FSR residues)",
        f"pLDDT >= {plddt_threshold} AND PAE < {pae_threshold} (scope={pae_scope})"
    )

    # Tier 1 cannot compute IR — it has no external truth
    audit.derived(
        "Tier 1 IR: NOT COMPUTABLE (no external reference)",
        "Tier 1 classifies confidence, not correctness. "
        "FalseVerify detection requires Tier 2."
    )

    return {
        'n_residues': n_total,
        'n_confident': n_confident,
        'n_uncertain': n_uncertain,
        'ptm': ptm,
        'plddt_threshold': plddt_threshold,
        'pae_threshold': pae_threshold,
        'pae_scope': pae_scope,
        'residues': residue_data,
        'audit': audit
    }


def print_results(results: dict):
    """Print Tier 1 results."""
    n = results['n_residues']
    nc = results['n_confident']
    nu = results['n_uncertain']

    print()
    print("=" * 70)
    print("TIER 1: INTERNAL METRICS ONLY")
    print(f"  pLDDT threshold: ≥ {results['plddt_threshold']}")
    print(f"  PAE threshold:   < {results['pae_threshold']} Å")
    print(f"  PAE scope:       {results['pae_scope']}")
    print(f"  pTM:             {results['ptm']}")
    print(f"  FSR residues:    {n}")
    print("=" * 70)
    print()
    print(f"  CONFIDENT: {nc:>3}  ({nc/n*100:.1f}%)")
    print(f"  uncertain: {nu:>3}  ({nu/n*100:.1f}%)")
    print()
    print(f"{'Res':>5} {'pLDDT':>6} {'PAE':>6} {'Class':>10}")
    print("-" * 30)
    for r in results['residues']:
        print(f"{r['resnum']:>5} {r['plddt']:>6.1f} {r['pae']:>6.1f} {r['tier1_class']:>10}")
    print()
    print("⚠ Tier 1 cannot detect FalseVerify. Confident ≠ correct.")
    print("  Tier 2 (structural comparison) required for SDI typing.")
    print()
```
