## supplement_v6 for "Confidence Without Verification: Screening pLDDT Unreliability in AlphaFold2 Fold-Switching Predictions": tier2_analysis.py

```
"""
Tier 2 Analysis: Structural Comparison

Compares predicted structure against experimental references
to produce full SDI typing at per-residue level.

This is where FalseVerify becomes detectable: a residue that
Tier 1 calls "confident" but is structurally far from the
alternate experimental fold.

The Tier 1 → Tier 2 gap is the core measurement:
  - What fraction of "confident" residues are actually wrong?
  - Do internal metrics provide ANY detection of these failures?
"""

import numpy as np
import json
from audit_trail import AuditTrail


def parse_pdb_ca(filename: str) -> dict:
    """
    Extract CA atom coordinates from PDB file.

    Returns:
        dict mapping 1-indexed residue number → numpy array [x, y, z]
    """
    cas = {}
    with open(filename) as f:
        for line in f:
            if line.startswith('ATOM') and line[12:16].strip() == 'CA':
                resnum = int(line[22:26].strip())
                x = float(line[30:38])
                y = float(line[38:46])
                z = float(line[46:54])
                cas[resnum] = np.array([x, y, z])
    return cas


def kabsch_align(P: np.ndarray, Q: np.ndarray):
    """
    Kabsch algorithm: optimal rotation to minimize RMSD.

    Args:
        P: Nx3 mobile coordinates
        Q: Nx3 reference coordinates

    Returns:
        (P_aligned, rmsd, per_residue_distances)
        All in the centered-on-Q frame.
    """
    p_center = P.mean(axis=0)
    q_center = Q.mean(axis=0)
    P_c = P - p_center
    Q_c = Q - q_center

    H = P_c.T @ Q_c
    U, S, Vt = np.linalg.svd(H)

    d = np.linalg.det(Vt.T @ U.T)
    sign_matrix = np.eye(3)
    sign_matrix[2, 2] = np.sign(d)

    R = Vt.T @ sign_matrix @ U.T
    P_aligned = P_c @ R.T

    dists = np.sqrt(np.sum((P_aligned - Q_c) ** 2, axis=1))
    rmsd = np.sqrt(np.mean(dists ** 2))

    return P_aligned, rmsd, dists


def run_tier2(
    pred_pdb_path: str,
    ref_fold1_pdb_path: str,
    ref_fold2_pdb_path: str,
    scores_json_path: str,
    fsr_residues: list,
    fold1_label: str = "fold1",
    fold2_label: str = "fold2",
    dist_threshold: float = 3.8,
    pae_threshold: float = 10.0,
    plddt_threshold: float = 70.0,
    pae_scope: str = "fsr_internal",
    audit: AuditTrail = None
) -> dict:
    """
    Tier 2: Full SDI typing with structural comparison.

    The prediction is compared to BOTH experimental folds.
    The fold it's closer to is the "predicted fold".
    The OTHER fold is the "alternate fold".
    Distance to the alternate fold determines "wrong" classification.

    Args:
        pred_pdb_path: ColabFold prediction PDB
        ref_fold1_pdb_path: Experimental reference, fold 1
        ref_fold2_pdb_path: Experimental reference, fold 2 (alternate)
        scores_json_path: ColabFold scores JSON
        fsr_residues: List of 1-indexed residue numbers in FSR
        fold1_label: Name for fold 1 (e.g., "2oug_C alpha")
        fold2_label: Name for fold 2 (e.g., "6c6s_D beta")
        dist_threshold: CA distance above this = wrong fold (default 3.8 Å)
        pae_threshold: PAE below this = confident (default 10.0 Å)
        plddt_threshold: pLDDT above this = confident (default 70.0)
        pae_scope: "fsr_internal" or "all_columns"
        audit: AuditTrail instance

    Returns:
        dict with per-residue SDI typing and audit trail
    """
    if audit is None:
        audit = AuditTrail()

    # Parse PDBs
    pred_ca = parse_pdb_ca(pred_pdb_path)
    ref1_ca = parse_pdb_ca(ref_fold1_pdb_path)
    ref2_ca = parse_pdb_ca(ref_fold2_pdb_path)

    audit.verified(
        f"Prediction PDB: {len(pred_ca)} CA atoms, "
        f"residues {min(pred_ca.keys())}-{max(pred_ca.keys())}",
        f"parse_pdb_ca({pred_pdb_path})"
    )
    audit.verified(
        f"{fold1_label} PDB: {len(ref1_ca)} CA atoms, "
        f"residues {min(ref1_ca.keys())}-{max(ref1_ca.keys())}",
        f"parse_pdb_ca({ref_fold1_pdb_path})"
    )
    audit.verified(
        f"{fold2_label} PDB: {len(ref2_ca)} CA atoms, "
        f"residues {min(ref2_ca.keys())}-{max(ref2_ca.keys())}",
        f"parse_pdb_ca({ref_fold2_pdb_path})"
    )

    # Find common FSR residues across all three structures
    common_fsr = sorted(
        set(fsr_residues) &
        set(pred_ca.keys()) &
        set(ref1_ca.keys()) &
        set(ref2_ca.keys())
    )

    if len(common_fsr) < len(fsr_residues):
        missing = len(fsr_residues) - len(common_fsr)
        audit.assumed(
            f"FSR coverage: {len(common_fsr)}/{len(fsr_residues)} residues resolved in all 3 structures",
            f"Intersection of residue sets",
            f"{missing} FSR residues not present in all structures. "
            f"Analysis limited to common residues {min(common_fsr)}-{max(common_fsr)}."
        )
    else:
        audit.verified(
            f"FSR coverage: all {len(common_fsr)} residues present in all 3 structures",
            "Intersection of residue sets"
        )

    # Build coordinate arrays
    pred_coords = np.array([pred_ca[r] for r in common_fsr])
    ref1_coords = np.array([ref1_ca[r] for r in common_fsr])
    ref2_coords = np.array([ref2_ca[r] for r in common_fsr])

    # Kabsch alignment on FSR residues
    _, rmsd1, dists1 = kabsch_align(pred_coords, ref1_coords)
    _, rmsd2, dists2 = kabsch_align(pred_coords, ref2_coords)

    audit.derived(
        f"FSR RMSD vs {fold1_label}: {rmsd1:.2f} Å",
        f"Kabsch alignment on {len(common_fsr)} FSR CA atoms"
    )
    audit.derived(
        f"FSR RMSD vs {fold2_label}: {rmsd2:.2f} Å",
        f"Kabsch alignment on {len(common_fsr)} FSR CA atoms"
    )

    # Determine which fold prediction resembles
    if rmsd1 < rmsd2:
        predicted_fold = fold1_label
        alternate_fold = fold2_label
        dists_predicted = dists1
        dists_alternate = dists2
    else:
        predicted_fold = fold2_label
        alternate_fold = fold1_label
        dists_predicted = dists2
        dists_alternate = dists1

    audit.derived(
        f"Prediction resembles {predicted_fold} (RMSD {min(rmsd1, rmsd2):.2f} Å); "
        f"alternate fold is {alternate_fold} (RMSD {max(rmsd1, rmsd2):.2f} Å)",
        "Lower RMSD = closer structural match"
    )

    # Load internal metrics for Tier 1/2 cross-reference
    with open(scores_json_path) as f:
        scores = json.load(f)
    plddt = np.array(scores['plddt'])
    pae_matrix = np.array(scores['pae'])
    ptm = scores.get('ptm', None)

    # Compute per-residue PAE
    fsr_0idx = [r - 1 for r in common_fsr]
    if pae_scope == "fsr_internal":
        pae_block = pae_matrix[np.ix_(fsr_0idx, fsr_0idx)]
        pae_per_res = np.mean(pae_block, axis=1)
    else:
        pae_per_res = np.mean(pae_matrix[fsr_0idx, :], axis=1)

    # SDI Typing per residue
    residue_data = []
    counts = {'FalseVerify': 0, 'SelfCatch': 0, 'Correct': 0, 'OverRefusal': 0}

    for i, resnum in enumerate(common_fsr):
        idx_0 = resnum - 1
        p = float(plddt[idx_0])
        pae_val = float(pae_per_res[i])
        d_pred = float(dists_predicted[i])
        d_alt = float(dists_alternate[i])

        confident = (p >= plddt_threshold) and (pae_val < pae_threshold)
        wrong_fold = d_alt > dist_threshold

        if confident and wrong_fold:
            sdi_type = "FalseVerify"
        elif confident and not wrong_fold:
            sdi_type = "Correct"
        elif not confident and wrong_fold:
            sdi_type = "SelfCatch"
        else:
            sdi_type = "OverRefusal"

        counts[sdi_type] += 1
        residue_data.append({
            'resnum': resnum,
            'plddt': p,
            'pae': pae_val,
            'confident': confident,
            'd_predicted_fold': d_pred,
            'd_alternate_fold': d_alt,
            'wrong_fold': wrong_fold,
            'sdi_type': sdi_type
        })

    n = len(common_fsr)
    fv = counts['FalseVerify']
    sc = counts['SelfCatch']
    cc = counts['Correct']
    orr = counts['OverRefusal']

    # IR
    ir = sc / (sc + fv) if (sc + fv) > 0 else 1.0

    audit.derived(
        f"Tier 2 SDI typing (n={n}): FV={fv} ({fv/n*100:.1f}%), "
        f"SC={sc} ({sc/n*100:.1f}%), Correct={cc}, OverRefusal={orr}",
        f"confident=(pLDDT≥{plddt_threshold}, PAE<{pae_threshold}), "
        f"wrong_fold=(d_alternate>{dist_threshold} Å)"
    )
    audit.derived(
        f"Tier 2 IR = SC/(SC+FV) = {sc}/({sc}+{fv}) = {ir:.3f}",
        "SDI Internalization Rate"
    )

    # The key finding: how many FV residues does Tier 1 detect?
    tier1_catches_fv = sum(
        1 for r in residue_data
        if r['sdi_type'] == 'FalseVerify' and not r['confident']
    )
    tier1_misses_fv = fv - tier1_catches_fv

    audit.derived(
        f"Tier 1 detection of Tier 2 FalseVerify: {tier1_catches_fv}/{fv} "
        f"({tier1_catches_fv/fv*100:.1f}% detected)" if fv > 0 else
        "No FalseVerify residues to detect",
        "Cross-reference: FV residues that Tier 1 classifies as uncertain"
    )

    if tier1_misses_fv > 0 and fv > 0:
        fv_plddt = [r['plddt'] for r in residue_data if r['sdi_type'] == 'FalseVerify']
        fv_pae = [r['pae'] for r in residue_data if r['sdi_type'] == 'FalseVerify']
        fv_dalt = [r['d_alternate_fold'] for r in residue_data if r['sdi_type'] == 'FalseVerify']
        audit.derived(
            f"FalseVerify zone: pLDDT {min(fv_plddt):.1f}-{max(fv_plddt):.1f}, "
            f"PAE {min(fv_pae):.1f}-{max(fv_pae):.1f}, "
            f"d_alternate {min(fv_dalt):.1f}-{max(fv_dalt):.1f} Å",
            "Range of metrics for FV residues"
        )

    # Threshold sensitivity for distance
    for alt_d in [dist_threshold - 1.0, dist_threshold + 1.0]:
        if alt_d <= 0:
            continue
        alt_fv = sum(
            1 for r in residue_data
            if r['confident'] and r['d_alternate_fold'] > alt_d
        )
        alt_sc = sum(
            1 for r in residue_data
            if not r['confident'] and r['d_alternate_fold'] > alt_d
        )
        alt_ir = alt_sc / (alt_sc + alt_fv) if (alt_sc + alt_fv) > 0 else 1.0
        audit.derived(
            f"Sensitivity (dist_thresh={alt_d:.1f} Å): FV={alt_fv}, SC={alt_sc}, IR={alt_ir:.3f}",
            f"Same classification with dist_threshold={alt_d:.1f}"
        )

    return {
        'n_residues': n,
        'common_fsr': common_fsr,
        'predicted_fold': predicted_fold,
        'alternate_fold': alternate_fold,
        'rmsd_fold1': rmsd1,
        'rmsd_fold2': rmsd2,
        'counts': counts,
        'fv_rate': fv / n if n > 0 else 0,
        'ir': ir,
        'tier1_detects_fv': tier1_catches_fv,
        'tier1_misses_fv': tier1_misses_fv,
        'ptm': ptm,
        'dist_threshold': dist_threshold,
        'plddt_threshold': plddt_threshold,
        'pae_threshold': pae_threshold,
        'pae_scope': pae_scope,
        'residues': residue_data,
        'audit': audit
    }


def print_results(results: dict):
    """Print Tier 2 results with full per-residue table."""
    n = results['n_residues']
    c = results['counts']

    print()
    print("=" * 80)
    print("TIER 2: STRUCTURAL SDI TYPING")
    print(f"  Framework: λ[C ≥ P ≥ E]")
    print(f"  Predicted fold:  {results['predicted_fold']} (RMSD {min(results['rmsd_fold1'], results['rmsd_fold2']):.2f} Å)")
    print(f"  Alternate fold:  {results['alternate_fold']} (RMSD {max(results['rmsd_fold1'], results['rmsd_fold2']):.2f} Å)")
    print(f"  pTM: {results['ptm']}")
    print(f"  Distance threshold: {results['dist_threshold']} Å")
    print(f"  Confidence: pLDDT ≥ {results['plddt_threshold']}, PAE < {results['pae_threshold']} ({results['pae_scope']})")
    print(f"  FSR residues: {n}")
    print("=" * 80)
    print()

    # Per-residue table
    print(f"{'Res':>5} {'pLDDT':>6} {'PAE':>6} {'Tier1':>8} {'d_pred':>7} {'d_alt':>7} {'SDI Type':>14}")
    print("-" * 60)
    for r in results['residues']:
        conf = "CONF" if r['confident'] else "unc"
        print(f"{r['resnum']:>5} {r['plddt']:>6.1f} {r['pae']:>6.1f} {conf:>8} "
              f"{r['d_predicted_fold']:>7.2f} {r['d_alternate_fold']:>7.2f} {r['sdi_type']:>14}")

    print()
    print(f"  FalseVerify:  {c['FalseVerify']:>3}  ({c['FalseVerify']/n*100:.1f}%)")
    print(f"  SelfCatch:    {c['SelfCatch']:>3}  ({c['SelfCatch']/n*100:.1f}%)")
    print(f"  Correct:      {c['Correct']:>3}  ({c['Correct']/n*100:.1f}%)")
    print(f"  OverRefusal:  {c['OverRefusal']:>3}  ({c['OverRefusal']/n*100:.1f}%)")
    print()
    print(f"  IR = SC/(SC+FV) = {results['ir']:.3f}")
    print()
    print(f"  Tier 1 detects {results['tier1_detects_fv']}/{c['FalseVerify']} FalseVerify residues")
    if results['tier1_misses_fv'] > 0:
        print(f"  ⚠ Internal metrics MISS {results['tier1_misses_fv']} FalseVerify residues")
    print()
```
