## supplement_v6 for "Confidence Without Verification: Screening pLDDT Unreliability in AlphaFold2 Fold-Switching Predictions": analyze_npy.py

```
#!/usr/bin/env python3
"""
Analyze aggregated numpy TM-score files from Porter benchmark
"""

import numpy as np
import os

def main():
    data_dir = r'C:\Users\robin\Desktop\substrate_agnostic_experiments\BlindFold\AF2_benchmark-AF2-benchmark\porterll-AF2_benchmark-40a57d7\data'
    
    npy_files = [
        'tmscores_all.npy',
        'tmscores_fs_all.npy', 
        'all_tmscores_af231_not.npy',
        'all_fstmscores_af231_not.npy',
    ]
    
    print("=" * 60)
    print("Analyzing Aggregated TM-Score Files")
    print("=" * 60)
    
    for npy_file in npy_files:
        filepath = os.path.join(data_dir, npy_file)
        if not os.path.exists(filepath):
            print(f"\n{npy_file}: NOT FOUND")
            continue
            
        data = np.load(filepath, allow_pickle=True)
        
        print(f"\n{npy_file}:")
        print(f"  Shape: {data.shape}")
        print(f"  Dtype: {data.dtype}")
        
        if data.dtype == np.float64 or data.dtype == np.float32:
            print(f"  Min: {np.min(data):.4f}")
            print(f"  Max: {np.max(data):.4f}")
            print(f"  Mean: {np.mean(data):.4f}")
            print(f"  Std: {np.std(data):.4f}")
            print(f"  First 20 values: {data.flatten()[:20]}")
        elif data.dtype == object:
            print(f"  Type of first element: {type(data.flat[0])}")
            if hasattr(data.flat[0], 'shape'):
                print(f"  Shape of first element: {data.flat[0].shape}")
            print(f"  First element: {data.flat[0]}")
        else:
            print(f"  First 10 values: {data.flatten()[:10]}")
    
    # Also check for any other npy files
    print("\n" + "=" * 60)
    print("All .npy files in data directory:")
    print("=" * 60)
    
    for f in os.listdir(data_dir):
        if f.endswith('.npy'):
            filepath = os.path.join(data_dir, f)
            data = np.load(filepath, allow_pickle=True)
            print(f"  {f}: shape={data.shape}, dtype={data.dtype}")
    
    # Check clean_chains and bfact_data directories
    print("\n" + "=" * 60)
    print("Checking subdirectories for additional data:")
    print("=" * 60)
    
    for subdir in ['clean_chains', 'bfact_data']:
        subpath = os.path.join(data_dir, subdir)
        if os.path.exists(subpath):
            contents = os.listdir(subpath)
            print(f"\n{subdir}/: {len(contents)} items")
            # Show first 10 items
            for item in sorted(contents)[:15]:
                print(f"  {item}")
            if len(contents) > 15:
                print(f"  ... and {len(contents) - 15} more")


if __name__ == '__main__':
    main()
```
