## supplement_v6 for "Confidence Without Verification: Screening pLDDT Unreliability in AlphaFold2 Fold-Switching Predictions": diagnose_data.py

```
#!/usr/bin/env python3
"""
Diagnostic script to find all tmscores.txt files and protein directories
"""

import os
import glob
from pathlib import Path

def find_all_files(root_dir, filename):
    """Recursively find all files matching filename"""
    matches = []
    for root, dirs, files in os.walk(root_dir):
        if filename in files:
            matches.append(os.path.join(root, filename))
    return matches

def main():
    # Base directory - adjust as needed
    base_dirs = [
        r'C:\Users\robin\Desktop\substrate_agnostic_experiments\BlindFold\AF2_benchmark-AF2-benchmark\porterll-AF2_benchmark-40a57d7',
        r'C:\Users\robin\Desktop\substrate_agnostic_experiments\BlindFold\AF2_benchmark-AF2-benchmark',
        r'C:\Users\robin\Desktop\substrate_agnostic_experiments\BlindFold',
    ]
    
    for base_dir in base_dirs:
        if not os.path.exists(base_dir):
            continue
            
        print(f"\n{'='*60}")
        print(f"Searching: {base_dir}")
        print('='*60)
        
        # Find all tmscores.txt files
        tmscores_files = find_all_files(base_dir, 'tmscores.txt')
        print(f"\nFound {len(tmscores_files)} tmscores.txt files:")
        for f in tmscores_files[:20]:
            # Get parent directory (protein ID)
            parent = os.path.basename(os.path.dirname(f))
            print(f"  {parent}: {f}")
        if len(tmscores_files) > 20:
            print(f"  ... and {len(tmscores_files) - 20} more")
        
        # Find all JSON score files
        json_files = []
        for root, dirs, files in os.walk(base_dir):
            for f in files:
                if '_scores_rank_' in f and f.endswith('.json'):
                    json_files.append(os.path.join(root, f))
        
        print(f"\nFound {len(json_files)} score JSON files")
        if json_files:
            # Group by protein
            proteins = set()
            for jf in json_files:
                # Go up directories to find protein
                parts = Path(jf).parts
                for i, p in enumerate(parts):
                    if p == 'data' and i + 1 < len(parts):
                        proteins.add(parts[i + 1])
                        break
            print(f"  Across {len(proteins)} proteins: {list(proteins)[:10]}...")
        
        # Check data directory structure
        data_dir = os.path.join(base_dir, 'data')
        if os.path.exists(data_dir):
            subdirs = [d for d in os.listdir(data_dir) if os.path.isdir(os.path.join(data_dir, d))]
            print(f"\nSubdirectories in data/: {len(subdirs)}")
            print(f"  First 10: {subdirs[:10]}")
            
            # Check what's in each subdir
            for sd in subdirs[:3]:
                sd_path = os.path.join(data_dir, sd)
                contents = os.listdir(sd_path)
                print(f"\n  {sd}/ contains:")
                for c in contents[:10]:
                    cp = os.path.join(sd_path, c)
                    if os.path.isdir(cp):
                        print(f"    {c}/ (dir)")
                    else:
                        print(f"    {c}")
        
        break  # Only check first existing directory


if __name__ == '__main__':
    main()
```
