## supplement_v6 for "Confidence Without Verification: Screening pLDDT Unreliability in AlphaFold2 Fold-Switching Predictions": extract_pse.py

```
"""Extract PDB structures from E. coli PSE files"""
import pymol
from pymol import cmd
import glob, os

pymol.finish_launching(['pymol', '-cq'])

base = r'C:\Users\robin\Desktop\MAGIC-Protein-Fold\Phase 2 - Porter\data\cf_random_repo\CF-random_software\Data\E_coli'
os.chdir(base)
os.makedirs('extracted_pdbs', exist_ok=True)

for pse in sorted(glob.glob('*.pse')):
    wp = pse.replace('.pse', '')
    cmd.load(pse)
    objects = cmd.get_object_list()
    n_states = cmd.count_states('all')
    print(f'{wp}: {len(objects)} objects, {n_states} total states')
    for obj in objects:
        ns = cmd.count_states(obj)
        for s in range(1, ns + 1):
            outfile = f'extracted_pdbs/{wp}_{obj}_s{s}.pdb'
            cmd.save(outfile, obj, state=s)
    cmd.delete('all')

print('Done.')
print(f'Total PDBs: {len(glob.glob("extracted_pdbs/*.pdb"))}')
```
