## supplement_v6 for "Confidence Without Verification: Screening pLDDT Unreliability in AlphaFold2 Fold-Switching Predictions": extract_pse_v2.py

```
"""Extract PDB structures from E. coli PSE files - NO PyMOL required.
PSE files are gzipped Python pickles containing molecular data."""
import gzip, pickle, glob, os, struct

base = r'C:\Users\robin\Desktop\MAGIC-Protein-Fold\Phase 2 - Porter\data\cf_random_repo\CF-random_software\Data\E_coli'
os.chdir(base)
os.makedirs('extracted_pdbs', exist_ok=True)

summary = []

for pse_file in sorted(glob.glob('*.pse')):
    wp = pse_file.replace('.pse', '')
    try:
        with gzip.open(pse_file, 'rb') as f:
            data = pickle.load(f, encoding='latin1')
    except Exception as e:
        print(f'{wp}: ERROR loading - {e}')
        summary.append((wp, 0, 0, 'ERROR'))
        continue

    # PSE structure: data is typically a list/dict with 'names' containing objects
    names = data.get('names', []) if isinstance(data, dict) else []
    
    obj_count = 0
    total_states = 0
    obj_names = []
    
    for entry in names:
        if entry is None:
            continue
        if not isinstance(entry, list) or len(entry) < 2:
            continue
        
        obj_name = entry[0] if isinstance(entry[0], str) else str(entry[0])
        obj_data = entry[1] if len(entry) > 1 else None
        
        if obj_data is None:
            continue
        
        # Check if it's a molecular object (type 1) or group/other
        if not isinstance(obj_data, list) or len(obj_data) < 1:
            continue
        
        obj_type = obj_data[0] if isinstance(obj_data[0], int) else -1
        
        # Type 1 = molecule
        if obj_type == 1 and len(obj_data) > 4:
            obj_count += 1
            obj_names.append(obj_name)
            
            # obj_data[4] contains coordinate sets (states)
            coord_sets = obj_data[4] if len(obj_data) > 4 else []
            if isinstance(coord_sets, list):
                n_states = len([cs for cs in coord_sets if cs is not None])
            else:
                n_states = 1
            total_states += n_states
            
            # Extract atom info and coordinates for each state
            # obj_data[2] = atom list, obj_data[4] = coordinate states
            atom_info = obj_data[2] if len(obj_data) > 2 else None
            
            if atom_info is None:
                continue
            
            for state_idx, cs in enumerate(coord_sets):
                if cs is None:
                    continue
                
                state_num = state_idx + 1
                outfile = f'extracted_pdbs/{wp}_{obj_name}_s{state_num}.pdb'
                
                try:
                    # cs is a coordinate set - try to get coordinates
                    # Format varies by PSE version
                    if isinstance(cs, list) and len(cs) > 0:
                        # Try to extract coords from the coordinate set
                        # Typically cs[0] contains packed coordinates
                        coords_raw = cs[0] if len(cs) > 0 else None
                        
                        if coords_raw is None:
                            continue
                        
                        # Coords may be bytes (packed floats) or list
                        if isinstance(coords_raw, bytes):
                            n_floats = len(coords_raw) // 4
                            coords = struct.unpack(f'{n_floats}f', coords_raw)
                        elif isinstance(coords_raw, (list, tuple)):
                            coords = coords_raw
                        else:
                            continue
                        
                        # Write PDB file
                        with open(outfile, 'w') as pdb:
                            pdb.write(f'REMARK  Extracted from {pse_file} object={obj_name} state={state_num}\n')
                            
                            n_atoms = len(coords) // 3
                            for i in range(min(n_atoms, len(atom_info))):
                                x = coords[i*3]
                                y = coords[i*3 + 1]
                                z = coords[i*3 + 2]
                                
                                ai = atom_info[i]
                                # atom_info fields vary, extract what we can
                                if isinstance(ai, (list, tuple)):
                                    # Try to get atom name, resn, chain, resi, b-factor
                                    resn = str(ai[0]) if len(ai) > 0 else 'UNK'
                                    chain = str(ai[1]) if len(ai) > 1 else 'A'
                                    name = str(ai[6]) if len(ai) > 6 else 'CA'
                                    resi = int(ai[3]) if len(ai) > 3 else i+1
                                    b_factor = float(ai[14]) if len(ai) > 14 else 0.0
                                    elem = str(ai[30]) if len(ai) > 30 else name[0]
                                    
                                    pdb.write(f'ATOM  {i+1:5d} {name:<4s} {resn:<3s} {chain:1s}{resi:4d}    '
                                             f'{x:8.3f}{y:8.3f}{z:8.3f}{1.0:6.2f}{b_factor:6.2f}'
                                             f'          {elem:>2s}\n')
                                else:
                                    pdb.write(f'ATOM  {i+1:5d}  CA  UNK A{i+1:4d}    '
                                             f'{x:8.3f}{y:8.3f}{z:8.3f}{1.0:6.2f}{0.0:6.2f}\n')
                            
                            pdb.write('END\n')
                    
                except Exception as e:
                    print(f'  Warning: {obj_name} state {state_num}: {e}')
                    continue
    
    print(f'{wp}: {obj_count} objects ({", ".join(obj_names)}), {total_states} total states')
    summary.append((wp, obj_count, total_states, ','.join(obj_names)))

# Write summary
print(f'\nDone. Processed {len(summary)} PSE files.')
total_pdbs = len(glob.glob('extracted_pdbs/*.pdb'))
print(f'Total PDB files extracted: {total_pdbs}')

# Save summary CSV
with open('extraction_summary.csv', 'w') as f:
    f.write('wp_id,n_objects,n_states,object_names\n')
    for wp, no, ns, names in summary:
        f.write(f'{wp},{no},{ns},{names}\n')
print('Summary saved to extraction_summary.csv')
```
