## supplement_v6 for "Confidence Without Verification: Screening pLDDT Unreliability in AlphaFold2 Fold-Switching Predictions": find_mapping.py

```
#!/usr/bin/env python3
"""
Find protein-to-index mapping in Porter data
Analyze the af231_not subset (AF2 failures)
"""

import numpy as np
import os
import glob

def main():
    base_dir = r'C:\Users\robin\Desktop\substrate_agnostic_experiments\BlindFold\AF2_benchmark-AF2-benchmark\porterll-AF2_benchmark-40a57d7'
    data_dir = os.path.join(base_dir, 'data')
    
    print("=" * 60)
    print("Searching for protein index mapping")
    print("=" * 60)
    
    # Check for any txt/csv files with protein lists
    for root, dirs, files in os.walk(base_dir):
        depth = root.replace(base_dir, '').count(os.sep)
        if depth > 2:
            continue
        for f in files:
            if f.endswith(('.txt', '.csv', '.tsv')) and 'readme' not in f.lower():
                filepath = os.path.join(root, f)
                try:
                    with open(filepath, 'r') as fh:
                        lines = fh.readlines()[:5]
                    if lines:
                        print(f"\n{f} (in {os.path.basename(root)}):")
                        for line in lines:
                            print(f"  {line.strip()[:80]}")
                except:
                    pass
    
    # Check for Python files that might define the ordering
    print("\n" + "=" * 60)
    print("Looking for Python scripts with protein lists")
    print("=" * 60)
    
    for root, dirs, files in os.walk(base_dir):
        depth = root.replace(base_dir, '').count(os.sep)
        if depth > 2:
            continue
        for f in files:
            if f.endswith('.py'):
                filepath = os.path.join(root, f)
                print(f"\n{f}:")
                try:
                    with open(filepath, 'r') as fh:
                        content = fh.read()
                    # Look for protein lists or dictionaries
                    if 'pdb' in content.lower() or 'protein' in content.lower():
                        # Show relevant excerpts
                        lines = content.split('\n')
                        for i, line in enumerate(lines):
                            if any(x in line.lower() for x in ['pdb', 'protein', 'chain', '2oug', '6c6s']):
                                print(f"  L{i+1}: {line[:100]}")
                except:
                    pass
    
    # Analyze the "not" arrays - these are AF2 failures
    print("\n" + "=" * 60)
    print("Analyzing AF2.3.1 failure subset")
    print("=" * 60)
    
    # Load the failure subset
    tmscores_fail = np.load(os.path.join(data_dir, 'all_tmscores_af231_not.npy'))
    fstmscores_fail = np.load(os.path.join(data_dir, 'all_fstmscores_af231_not.npy'))
    
    print(f"\nall_tmscores_af231_not: {len(tmscores_fail)} values")
    print(f"all_fstmscores_af231_not: {len(fstmscores_fail)} values")
    
    # The pattern appears to be pairs (TM to Fold1, TM to Fold2)
    # Let's reshape and analyze
    n_pairs = len(fstmscores_fail) // 2
    print(f"\nIf paired (fold1, fold2): {n_pairs} prediction pairs")
    
    # Reshape to pairs
    fs_pairs = fstmscores_fail[:n_pairs*2].reshape(-1, 2)
    print(f"Reshaped: {fs_pairs.shape}")
    print(f"\nFirst 10 pairs (TM_fold1, TM_fold2):")
    for i, (tm1, tm2) in enumerate(fs_pairs[:10]):
        dominant = "Fold1" if tm1 > tm2 else "Fold2"
        margin = abs(tm1 - tm2)
        print(f"  {i}: ({tm1:.2f}, {tm2:.2f}) -> {dominant} (margin={margin:.2f})")
    
    # From success spreadsheet: AF2.3.1 succeeded on 11/93 proteins
    # So failures = 82 proteins
    # 1880/82 ≈ 23 predictions per protein (if multiple MSA clusters × 5 models)
    print(f"\n1880 values / 82 failing proteins ≈ {1880/82:.1f} predictions each")
    print(f"If 5 models × ~4-5 clusters = ~20-25 predictions per protein ✓")
    
    # Compute disagreement statistics
    print("\n" + "=" * 60)
    print("Disagreement analysis (proxy for V_PC)")
    print("=" * 60)
    
    # For each "protein" (assuming 20 predictions each)
    preds_per_protein = 20  # Approximate
    n_proteins = len(fs_pairs) // preds_per_protein
    
    print(f"Estimated {n_proteins} proteins with ~{preds_per_protein} predictions each")
    
    for p in range(min(5, n_proteins)):
        start = p * preds_per_protein
        end = start + preds_per_protein
        protein_preds = fs_pairs[start:end]
        
        # Dominant fold for each prediction
        dominant_folds = [1 if tm1 > tm2 else 2 for tm1, tm2 in protein_preds]
        
        # Agreement rate
        fold1_count = dominant_folds.count(1)
        fold2_count = dominant_folds.count(2)
        agreement = max(fold1_count, fold2_count) / len(dominant_folds)
        
        # Mean "confidence" (max TM-score per prediction)
        mean_max_tm = np.mean([max(tm1, tm2) for tm1, tm2 in protein_preds])
        
        # V_PC proxy
        vpc = mean_max_tm * (1 - agreement)
        
        print(f"\nProtein {p}:")
        print(f"  Fold1 predictions: {fold1_count}, Fold2: {fold2_count}")
        print(f"  Agreement: {agreement:.2f}")
        print(f"  Mean max TM: {mean_max_tm:.2f}")
        print(f"  V_PC proxy: {vpc:.3f}")


if __name__ == '__main__':
    main()
```
