## supplement_v6 for "Confidence Without Verification: Screening pLDDT Unreliability in AlphaFold2 Fold-Switching Predictions": probe_pse.py

```
"""Probe first PSE file to understand its internal structure."""
import pickle, glob, os

base = r'C:\Users\robin\Desktop\MAGIC-Protein-Fold\Phase 2 - Porter\data\cf_random_repo\CF-random_software\Data\E_coli'
os.chdir(base)

pse_file = sorted(glob.glob('*.pse'))[0]
print(f'Probing: {pse_file} ({os.path.getsize(pse_file)} bytes)')

with open(pse_file, 'rb') as f:
    data = pickle.load(f, encoding='latin1')

print(f'\nTop-level type: {type(data).__name__}')

if isinstance(data, dict):
    print(f'Keys: {list(data.keys())}')
    for k, v in data.items():
        if isinstance(v, list):
            print(f'  {k}: list of {len(v)} items')
            if len(v) > 0 and v[0] is not None:
                print(f'    first item type: {type(v[0]).__name__}')
                if isinstance(v[0], list) and len(v[0]) > 0:
                    print(f'    first item[0]: {v[0][0]!r}')
                    if len(v[0]) > 1 and isinstance(v[0][1], list):
                        print(f'    first item[1] type: list of {len(v[0][1])} items')
                        print(f'    first item[1][0]: {v[0][1][0]!r}')
        elif isinstance(v, (str, int, float, bytes)):
            print(f'  {k}: {v!r}')
        else:
            print(f'  {k}: {type(v).__name__}')
    
    # Look at 'names' entries more carefully
    if 'names' in data:
        print(f'\n--- NAMES entries ---')
        for i, entry in enumerate(data['names']):
            if entry is None:
                continue
            if isinstance(entry, list) and len(entry) >= 2:
                obj_name = entry[0]
                obj_data = entry[1]
                if isinstance(obj_data, list) and len(obj_data) > 0:
                    obj_type = obj_data[0]
                    print(f'  [{i}] name={obj_name!r}, type={obj_type}, data_len={len(obj_data)}')
                    
                    if obj_type == 1 and len(obj_data) > 4:
                        # Molecule - check coord sets
                        atom_info = obj_data[2]
                        coord_sets = obj_data[4]
                        print(f'       atoms: {len(atom_info) if isinstance(atom_info, list) else "?"}'
                              f', coord_sets: {len(coord_sets) if isinstance(coord_sets, list) else "?"}')
                        
                        # Look at first atom info
                        if isinstance(atom_info, list) and len(atom_info) > 0:
                            ai = atom_info[0]
                            print(f'       first atom info ({len(ai)} fields): {ai[:20]}...' if len(ai) > 20 else f'       first atom info: {ai}')
                        
                        # Look at first coord set
                        if isinstance(coord_sets, list) and len(coord_sets) > 0:
                            cs = coord_sets[0]
                            if cs is not None:
                                print(f'       first coord set type: {type(cs).__name__}, len={len(cs) if hasattr(cs, "__len__") else "?"}')
                                if isinstance(cs, list) and len(cs) > 0:
                                    print(f'       cs[0] type: {type(cs[0]).__name__}, len={len(cs[0]) if hasattr(cs[0], "__len__") else "?"}')
                                    if isinstance(cs[0], bytes):
                                        print(f'       cs[0] first 24 bytes: {cs[0][:24].hex()}')
                else:
                    print(f'  [{i}] name={obj_name!r}, data type={type(obj_data).__name__}')

print('\nDone probing.')
```
