## supplement_v6 for "Confidence Without Verification: Screening pLDDT Unreliability in AlphaFold2 Fold-Switching Predictions": probe_pse_v2.py

```
"""Probe PSE file with mock PyMOL module to bypass import errors."""
import sys, types, pickle, glob, os

# Create a fake pymol module hierarchy so pickle.load doesn't crash
class MockModule(types.ModuleType):
    def __getattr__(self, name):
        # Return a dummy class for any attribute access
        mock = types.ModuleType(f'{self.__name__}.{name}')
        mock.__class__ = type(self)
        sys.modules[f'{self.__name__}.{name}'] = mock
        return mock

# Register mock pymol and all submodules pickle might reference
for mod_name in ['pymol', 'pymol.wizarding', 'pymol.viewing', 'pymol.setting',
                 'pymol.querying', 'pymol.selecting', 'pymol.editing',
                 'pymol.commanding', 'pymol.controlling', 'pymol.completing',
                 'pymol.moving', 'pymol.fitting', 'pymol.exporting',
                 'pymol.importing', 'pymol.creating', 'pymol.helping',
                 'pymol.measuring', 'pymol.experimenting', 'pymol.morphing',
                 'pymol.computing', 'pymol.sensing', 'pymol.wizard',
                 'pymol._symmetry', 'pymol.cgo', 'pymol.vfont',
                 'pymol.preset', 'pymol.menu', 'pymol.cmd',
                 'pymol._cmd', 'pymol.CmdException', 'pymol.constants',
                 'pymol.colorprinting', 'pymol.colorramping',
                 'chempy', 'chempy.models', 'chempy.cpv', 'chempy.place',
                 'chempy.champ', 'chempy.bmin', 'chempy.io',
                 'chempy.io.pdb', 'chempy.io.mol', 'chempy.io.mol2',
                 'chempy.io.mmd', 'chempy.io.sdf', 'chempy.io.xyz',
                 'chempy.models.Indexed']:
    if mod_name not in sys.modules:
        mock = MockModule(mod_name)
        sys.modules[mod_name] = mock

# Also need a generic unpickler that handles unknown classes
class GenericUnpickler(pickle.Unpickler):
    def find_class(self, module, name):
        # Try normal resolution first
        try:
            return super().find_class(module, name)
        except (ImportError, AttributeError):
            # Return a generic class that accepts any init args
            def make_generic(*args, **kwargs):
                obj = object.__new__(GenericObj)
                obj._args = args
                obj._kwargs = kwargs
                obj._class_name = f'{module}.{name}'
                return obj
            
            class GenericObj:
                def __init__(self, *args, **kwargs):
                    self._args = args
                    self._kwargs = kwargs
                def __reduce__(self):
                    return (make_generic, self._args)
                def __setstate__(self, state):
                    if isinstance(state, dict):
                        self.__dict__.update(state)
                    else:
                        self._state = state
                def __repr__(self):
                    return f'<{self._class_name}>'
            
            GenericObj.__name__ = name
            GenericObj.__qualname__ = f'{module}.{name}'
            return GenericObj

# Now try loading
base = r'C:\Users\robin\Desktop\MAGIC-Protein-Fold\Phase 2 - Porter\data\cf_random_repo\CF-random_software\Data\E_coli'
os.chdir(base)

pse_file = sorted(glob.glob('*.pse'))[0]
print(f'Probing: {pse_file} ({os.path.getsize(pse_file)} bytes)')

with open(pse_file, 'rb') as f:
    unpickler = GenericUnpickler(f, encoding='latin1')
    data = unpickler.load()

print(f'\nTop-level type: {type(data).__name__}')

if isinstance(data, dict):
    print(f'Keys: {list(data.keys())}')
    for k, v in data.items():
        if isinstance(v, list):
            non_none = [x for x in v if x is not None]
            print(f'  {k}: list of {len(v)} items ({len(non_none)} non-None)')
            if non_none:
                first = non_none[0]
                print(f'    first non-None type: {type(first).__name__}')
                if isinstance(first, list) and len(first) >= 2:
                    print(f'    first[0]: {first[0]!r}')
                    if isinstance(first[1], list):
                        print(f'    first[1]: list of {len(first[1])} items, first[1][0]={first[1][0]!r}')
        elif isinstance(v, (str, int, float)):
            vr = repr(v)
            print(f'  {k}: {vr[:100]}')
        else:
            print(f'  {k}: {type(v).__name__}')

    if 'names' in data:
        print(f'\n--- NAMES entries (non-None) ---')
        for i, entry in enumerate(data['names']):
            if entry is None:
                continue
            if isinstance(entry, list) and len(entry) >= 2:
                obj_name = entry[0]
                obj_data = entry[1]
                if isinstance(obj_data, list) and len(obj_data) > 0:
                    obj_type = obj_data[0]
                    print(f'\n  [{i}] name={obj_name!r}, type_id={obj_type}, data fields={len(obj_data)}')
                    
                    if obj_type == 1 and len(obj_data) > 4:
                        atom_info = obj_data[2]
                        coord_sets = obj_data[4]
                        n_atoms = len(atom_info) if isinstance(atom_info, list) else '?'
                        n_cs = len(coord_sets) if isinstance(coord_sets, list) else '?'
                        print(f'       MOLECULE: {n_atoms} atoms, {n_cs} coord sets')
                        
                        if isinstance(atom_info, list) and len(atom_info) > 0:
                            ai = atom_info[0]
                            if isinstance(ai, (list, tuple)):
                                print(f'       atom[0] ({len(ai)} fields):')
                                for j, field in enumerate(ai):
                                    print(f'         [{j}] = {field!r}')
                                    if j > 35:
                                        print(f'         ... ({len(ai)} total fields)')
                                        break
                            else:
                                print(f'       atom[0] type: {type(ai).__name__}')
                                if hasattr(ai, '__dict__'):
                                    print(f'       atom[0] attrs: {list(ai.__dict__.keys())[:20]}')
                        
                        if isinstance(coord_sets, list) and len(coord_sets) > 0:
                            cs = coord_sets[0]
                            if cs is not None:
                                print(f'       cs[0] type: {type(cs).__name__}')
                                if isinstance(cs, list):
                                    print(f'       cs[0] len: {len(cs)}')
                                    if len(cs) > 0:
                                        print(f'       cs[0][0] type: {type(cs[0]).__name__}, len={len(cs[0]) if hasattr(cs[0], "__len__") else "?"}')
                                elif hasattr(cs, '__dict__'):
                                    print(f'       cs[0] attrs: {list(cs.__dict__.keys())[:20]}')
else:
    print(f'Unexpected top-level type. Checking attributes...')
    if hasattr(data, '__dict__'):
        print(f'Attributes: {list(data.__dict__.keys())[:30]}')

print('\nDone probing.')
```
