## supplement_v6 for "Confidence Without Verification: Screening pLDDT Unreliability in AlphaFold2 Fold-Switching Predictions": probe_pse_v3.py

```
"""Probe PSE file - v3 with fully callable mocks."""
import sys, types, pickle, glob, os

# Create callable mock that handles ANY attribute/call pattern
class MockCallable:
    def __init__(self, name='mock'):
        self._name = name
    def __call__(self, *args, **kwargs):
        return MockCallable(f'{self._name}()')
    def __getattr__(self, name):
        return MockCallable(f'{self._name}.{name}')
    def __repr__(self):
        return f'<Mock:{self._name}>'
    def __bool__(self):
        return False
    def __setstate__(self, state):
        if isinstance(state, dict):
            self.__dict__.update(state)
        else:
            self._state = state

# Override find_class in unpickler to handle everything
class SafeUnpickler(pickle.Unpickler):
    def find_class(self, module, name):
        # Let builtins and standard lib through
        safe_modules = {'builtins', 'collections', 'collections.abc', 
                        'copy_reg', 'copyreg', '_codecs', 'codecs',
                        'numpy', 'numpy.core', 'numpy.core.multiarray'}
        
        for safe in safe_modules:
            if module == safe or module.startswith(safe + '.'):
                try:
                    return super().find_class(module, name)
                except:
                    pass
        
        # For pymol/chempy/anything else: return a class that 
        # accepts any constructor args and stores state
        def make_obj(*args, **kwargs):
            obj = object.__new__(StorageObj)
            obj._mock_class = f'{module}.{name}'
            obj._args = args
            obj._kwargs = kwargs
            return obj
        
        class StorageObj:
            _mock_class = f'{module}.{name}'
            def __init__(self, *args, **kwargs):
                self._args = args
                self._kwargs = kwargs
            def __setstate__(self, state):
                if isinstance(state, dict):
                    self.__dict__.update(state)
                else:
                    self._state = state
            def __reduce__(self):
                return (make_obj, self._args)
            def __repr__(self):
                return f'<{self._mock_class}>'
            def __len__(self):
                return 0
            def __iter__(self):
                return iter([])
            def __getattr__(self, n):
                if n.startswith('_'):
                    raise AttributeError(n)
                return MockCallable(f'{self._mock_class}.{n}')
        
        StorageObj.__name__ = name
        return StorageObj

# Load the PSE
base = r'C:\Users\robin\Desktop\MAGIC-Protein-Fold\Phase 2 - Porter\data\cf_random_repo\CF-random_software\Data\E_coli'
os.chdir(base)

pse_file = sorted(glob.glob('*.pse'))[0]
print(f'Probing: {pse_file} ({os.path.getsize(pse_file)} bytes)')

with open(pse_file, 'rb') as f:
    data = SafeUnpickler(f, encoding='latin1').load()

print(f'Top-level type: {type(data).__name__}')

if isinstance(data, dict):
    print(f'Keys: {list(data.keys())}')
    for k, v in data.items():
        if isinstance(v, list):
            non_none = [x for x in v if x is not None]
            print(f'  {k}: list[{len(v)}] ({len(non_none)} non-None)')
            if non_none:
                first = non_none[0]
                print(f'    first type: {type(first).__name__}')
                if isinstance(first, list) and len(first) >= 2:
                    print(f'    [0]={first[0]!r}')
                    if isinstance(first[1], list):
                        print(f'    [1]=list[{len(first[1])}], [1][0]={first[1][0]!r}')
        elif isinstance(v, (str, int, float)):
            print(f'  {k}: {repr(v)[:100]}')
        else:
            t = type(v).__name__
            mc = getattr(v, '_mock_class', '')
            print(f'  {k}: {t} {mc}')
            if hasattr(v, '__dict__'):
                attrs = {k2:type(v2).__name__ for k2,v2 in v.__dict__.items() if not k2.startswith('_')}
                if attrs:
                    print(f'    attrs: {attrs}')

    if 'names' in data:
        print(f'\n--- NAMES ---')
        for i, entry in enumerate(data['names']):
            if entry is None:
                continue
            if isinstance(entry, list) and len(entry) >= 2:
                nm = entry[0]
                od = entry[1]
                if isinstance(od, list) and len(od) > 0:
                    ot = od[0]
                    print(f'\n  [{i}] name={nm!r} type_id={ot} fields={len(od)}')
                    if ot == 1 and len(od) > 4:
                        ai = od[2]
                        cs = od[4]
                        na = len(ai) if isinstance(ai, list) else '?'
                        nc = len(cs) if isinstance(cs, list) else '?'
                        print(f'       atoms={na} coord_sets={nc}')
                        # First atom
                        if isinstance(ai, list) and ai:
                            a0 = ai[0]
                            if isinstance(a0, (list, tuple)):
                                print(f'       atom[0] ({len(a0)} fields):')
                                for j, fld in enumerate(a0[:40]):
                                    print(f'         [{j:2d}] {fld!r}')
                            elif hasattr(a0, '__dict__'):
                                d = {k2:repr(v2)[:60] for k2,v2 in a0.__dict__.items() if not k2.startswith('_mock')}
                                print(f'       atom[0] is object with attrs:')
                                for k2,v2 in sorted(d.items()):
                                    print(f'         {k2} = {v2}')
                        # First coord set
                        if isinstance(cs, list) and cs:
                            c0 = cs[0]
                            if c0 is not None:
                                print(f'       cs[0] type={type(c0).__name__}')
                                if isinstance(c0, list):
                                    print(f'       cs[0] len={len(c0)}')
                                    if c0 and isinstance(c0[0], bytes):
                                        print(f'       cs[0][0] bytes len={len(c0[0])}')
                                elif hasattr(c0, '__dict__'):
                                    d = {k2:repr(v2)[:60] for k2,v2 in c0.__dict__.items() if not k2.startswith('_mock')}
                                    print(f'       cs[0] attrs:')
                                    for k2,v2 in sorted(d.items()):
                                        print(f'         {k2} = {v2}')
                elif hasattr(od, '__dict__'):
                    d = {k2:type(v2).__name__ for k2,v2 in od.__dict__.items() if not k2.startswith('_')}
                    print(f'  [{i}] name={nm!r} obj_attrs={d}')
else:
    print('Not a dict.')
    if hasattr(data, '__dict__'):
        for k,v in data.__dict__.items():
            if not k.startswith('_'):
                print(f'  {k}: {type(v).__name__}')

print('\nDone.')
```
