## supplement_v6 for "Confidence Without Verification: Screening pLDDT Unreliability in AlphaFold2 Fold-Switching Predictions": probe_pse_v4.py

```
"""Deep probe of PSE names structure."""
import sys, types, pickle, glob, os

class SafeUnpickler(pickle.Unpickler):
    def find_class(self, module, name):
        safe = {'builtins','collections','collections.abc','copy_reg','copyreg','_codecs','codecs'}
        for s in safe:
            if module == s or module.startswith(s + '.'):
                try: return super().find_class(module, name)
                except: pass
        def make(*a, **kw):
            o = object.__new__(S)
            o._mc = f'{module}.{name}'
            o._a = a
            return o
        class S:
            _mc = f'{module}.{name}'
            def __init__(s, *a, **kw): s._a = a
            def __setstate__(s, st):
                if isinstance(st, dict): s.__dict__.update(st)
                else: s._state = st
            def __reduce__(s): return (make, s._a)
            def __repr__(s): return f'<{s._mc}>'
        S.__name__ = name
        return S

base = r'C:\Users\robin\Desktop\MAGIC-Protein-Fold\Phase 2 - Porter\data\cf_random_repo\CF-random_software\Data\E_coli'
os.chdir(base)
pse_file = sorted(glob.glob('*.pse'))[0]

with open(pse_file, 'rb') as f:
    data = SafeUnpickler(f, encoding='latin1').load()

names = data['names']
print(f'names has {len(names)} entries')

for i, entry in enumerate(names):
    if entry is None:
        print(f'  [{i}] None')
        continue
    
    print(f'\n  [{i}] type={type(entry).__name__}, len={len(entry) if hasattr(entry,"__len__") else "?"}')
    
    if isinstance(entry, list):
        for j, item in enumerate(entry):
            if item is None:
                print(f'    [{j}] None')
            elif isinstance(item, (str, int, float)):
                print(f'    [{j}] {item!r}')
            elif isinstance(item, list):
                print(f'    [{j}] list[{len(item)}]')
                if len(item) > 0:
                    print(f'      [0] = {type(item[0]).__name__}: {repr(item[0])[:100]}')
                if len(item) > 1:
                    print(f'      [1] = {type(item[1]).__name__}: {repr(item[1])[:100]}')
                if len(item) > 2:
                    t2 = type(item[2]).__name__
                    if isinstance(item[2], list):
                        print(f'      [2] = list[{len(item[2])}]')
                        if item[2] and isinstance(item[2][0], (list,tuple)):
                            print(f'        [2][0] = {type(item[2][0]).__name__}[{len(item[2][0])}]')
                    else:
                        print(f'      [2] = {t2}: {repr(item[2])[:100]}')
                if len(item) > 3:
                    print(f'      [3] = {type(item[3]).__name__}: {repr(item[3])[:80]}')
                if len(item) > 4:
                    t4 = type(item[4]).__name__
                    if isinstance(item[4], list):
                        print(f'      [4] = list[{len(item[4])}]')
                        for k, cs in enumerate(item[4][:3]):
                            if cs is not None:
                                print(f'        [4][{k}] = {type(cs).__name__}')
                                if hasattr(cs, '__dict__'):
                                    for ak, av in sorted(cs.__dict__.items()):
                                        if ak.startswith('_'): continue
                                        avr = repr(av)[:80]
                                        print(f'          .{ak} = {avr}')
                    else:
                        print(f'      [4] = {t4}: {repr(item[4])[:80]}')
                for idx in range(5, min(len(item), 8)):
                    print(f'      [{idx}] = {type(item[idx]).__name__}: {repr(item[idx])[:80]}')
            elif hasattr(item, '__dict__'):
                mc = getattr(item, '_mc', type(item).__name__)
                print(f'    [{j}] {mc}')
                for ak, av in sorted(item.__dict__.items()):
                    if ak.startswith('_'): continue
                    avr = repr(av)[:100]
                    print(f'      .{ak} = {avr}')
            else:
                print(f'    [{j}] {type(item).__name__}: {repr(item)[:100]}')
    elif hasattr(entry, '__dict__'):
        mc = getattr(entry, '_mc', type(entry).__name__)
        print(f'  [{i}] object: {mc}')
        for ak, av in sorted(entry.__dict__.items()):
            if ak.startswith('_'): continue
            avr = repr(av)[:100]
            print(f'    .{ak} = {avr}')

print('\nDone.')
```
