## supplement_v6 for "Confidence Without Verification: Screening pLDDT Unreliability in AlphaFold2 Fold-Switching Predictions": search_broader.py

```
#!/usr/bin/env python3
"""
Broader search for AF2 prediction data
"""

import os
from pathlib import Path
from collections import defaultdict

def main():
    base = r'C:\Users\robin\Desktop\substrate_agnostic_experiments\BlindFold'
    
    print("Searching entire BlindFold directory for prediction data...")
    print(f"Base: {base}\n")
    
    # Track what we find
    json_scores = []
    pdb_files = []
    tmscores = []
    npy_files = []
    
    # Walk entire tree
    for root, dirs, files in os.walk(base):
        # Skip very deep paths
        depth = root.replace(base, '').count(os.sep)
        if depth > 8:
            continue
            
        for f in files:
            fp = os.path.join(root, f)
            
            if '_scores_' in f and f.endswith('.json'):
                json_scores.append(fp)
            elif f.endswith('.pdb'):
                pdb_files.append(fp)
            elif 'tmscore' in f.lower() and f.endswith('.txt'):
                tmscores.append(fp)
            elif f.endswith('.npy') and 'tmscore' in f.lower():
                npy_files.append(fp)
    
    print(f"Found {len(json_scores)} score JSON files")
    print(f"Found {len(pdb_files)} PDB files")
    print(f"Found {len(tmscores)} tmscore txt files")
    print(f"Found {len(npy_files)} tmscore npy files")
    
    # Show unique directories containing score JSONs
    if json_scores:
        print("\n--- Score JSON locations ---")
        dirs = set(os.path.dirname(f) for f in json_scores)
        for d in list(dirs)[:10]:
            count = sum(1 for f in json_scores if os.path.dirname(f) == d)
            print(f"  {d}: {count} files")
    
    # Show tmscore files
    if tmscores:
        print("\n--- TM-score txt files ---")
        for f in tmscores[:20]:
            print(f"  {f}")
    
    # Show npy files
    if npy_files:
        print("\n--- TM-score npy files ---")
        for f in npy_files[:10]:
            print(f"  {f}")
    
    # Check what's in the main data directories
    print("\n--- Top-level directories in BlindFold ---")
    for item in sorted(os.listdir(base)):
        item_path = os.path.join(base, item)
        if os.path.isdir(item_path):
            subcount = len(os.listdir(item_path))
            print(f"  {item}/ ({subcount} items)")
    
    # Look specifically for protein ID patterns (4-char PDB codes)
    print("\n--- Looking for PDB-code directories ---")
    pdb_pattern_dirs = []
    for root, dirs, files in os.walk(base):
        depth = root.replace(base, '').count(os.sep)
        if depth > 4:
            continue
        for d in dirs:
            # PDB codes are 4 chars, often with chain suffix like 2oug_C
            if len(d) >= 4 and d[0].isdigit() and '_' in d:
                pdb_pattern_dirs.append(os.path.join(root, d))
    
    print(f"Found {len(pdb_pattern_dirs)} potential protein directories")
    for d in pdb_pattern_dirs[:20]:
        print(f"  {d}")


if __name__ == '__main__':
    main()
```
