## supplement_v6 for "Confidence Without Verification: Screening pLDDT Unreliability in AlphaFold2 Fold-Switching Predictions": verify_bccipa.py

```
import re

# Read the BCCIPa files
with open('/mnt/user-data/uploads/tmscores_a_unrelaxed.txt', 'r') as f:
    lines = [l.strip() for l in f if l.strip()]

print("=== BCCIP-α VERIFICATION ===\n")

print(f"Number of runs: {len(lines)}")

all_tm = []
run_data = {}

for line in lines:
    match = re.match(r'(\S+)\s*:\s*(.+)', line)
    if match:
        run_id = match.group(1)
        values_str = match.group(2)
        # BCCIPa format appears to be single TM values, not pairs
        values = [v.strip() for v in values_str.split(',') if v.strip()]
        run_data[run_id] = []
        for i, v in enumerate(values):
            try:
                tm = float(v)
                all_tm.append({'run': run_id, 'model': i, 'tm': tm})
                run_data[run_id].append(tm)
            except:
                pass

print(f"Total predictions parsed: {len(all_tm)}")
if run_data:
    first_run = list(run_data.keys())[0]
    print(f"Models per run (first run): {len(run_data[first_run])}")

# Verify claim: 3750 predictions
print(f"\nCLAIM C-BCCIPa-001: 3750 predictions")
print(f"  Actual: {len(all_tm)}")

# Check max
if all_tm:
    max_pred = max(all_tm, key=lambda x: x['tm'])
    print(f"\nCLAIM: max TM ≈ 0.41")
    print(f"  Actual max TM: {max_pred['tm']}")
    print(f"  Run: {max_pred['run']}, Model: {max_pred['model']}")

# Check hits >= 0.5
hits_05 = [p for p in all_tm if p['tm'] >= 0.5]
print(f"\nCLAIM: 0 hits >= 0.5 (100% failure)")
print(f"  Hits >= 0.5: {len(hits_05)}")
print(f"  STATUS: {'VERIFIED ✓' if len(hits_05) == 0 else 'MISMATCH ✗'}")

# Distribution summary
if all_tm:
    tms = [p['tm'] for p in all_tm]
    print(f"\n=== DISTRIBUTION ===")
    print(f"Min: {min(tms)}")
    print(f"Max: {max(tms)}")
    print(f"Mean: {sum(tms)/len(tms):.3f}")
    
    # Count by ranges
    ranges = [(0.0, 0.2), (0.2, 0.25), (0.25, 0.3), (0.3, 0.35), (0.35, 0.4), (0.4, 0.5)]
    print(f"\nDistribution by range:")
    for low, high in ranges:
        count = sum(1 for t in tms if low <= t < high)
        print(f"  [{low:.2f}, {high:.2f}): {count} ({100*count/len(tms):.1f}%)")
```
