## supplement_v6 for "Confidence Without Verification: Screening pLDDT Unreliability in AlphaFold2 Fold-Switching Predictions": verify_sa1.py

```
import re

# Read the unrelaxed file (all 50 models per run)
with open('/mnt/user-data/uploads/tmscores_unrelaxed.txt', 'r') as f:
    lines = [l.strip() for l in f if l.strip()]

print("=== SA1 VERIFICATION ===\n")

# Count runs and predictions
print(f"Number of runs: {len(lines)}")

all_predictions = []
run_data = {}

for line in lines:
    # Parse: run_id : tm1 tm2,tm1 tm2,...
    match = re.match(r'(\S+)\s*:\s*(.+)', line)
    if match:
        run_id = match.group(1)
        pairs_str = match.group(2)
        # Split by comma, parse each pair
        pairs = [p.strip() for p in pairs_str.split(',') if p.strip()]
        run_data[run_id] = []
        for i, pair in enumerate(pairs):
            parts = pair.split()
            if len(parts) == 2:
                tm1, tm2 = float(parts[0]), float(parts[1])
                all_predictions.append({
                    'run': run_id,
                    'model_idx': i,
                    'tm_fold1': tm1,
                    'tm_fold2': tm2
                })
                run_data[run_id].append((tm1, tm2))

print(f"Total predictions parsed: {len(all_predictions)}")
print(f"Models per run (first run): {len(run_data[list(run_data.keys())[0]])}")

# Verify claim: 51 runs × 50 models = 2550
expected = 51 * 50
actual = len(all_predictions)
print(f"\nCLAIM C-Sa1-001: 2550 predictions")
print(f"  Expected: {expected}, Actual: {actual}")
print(f"  STATUS: {'VERIFIED ✓' if actual == expected else 'MISMATCH ✗'}")

# Find global max TM(Fold2)
max_fold2 = max(all_predictions, key=lambda x: x['tm_fold2'])
print(f"\n=== MAX FOLD2 ANALYSIS ===")
print(f"Global max TM(Fold2): {max_fold2['tm_fold2']}")
print(f"  Run: {max_fold2['run']}, Model: {max_fold2['model_idx']}")
print(f"  TM(Fold1) at this point: {max_fold2['tm_fold1']}")

# Check run 0_011 specifically
print(f"\n=== RUN 0_011 ANALYSIS ===")
if '0_011' in run_data:
    run_011 = run_data['0_011']
    print(f"Models in run 0_011: {len(run_011)}")
    
    # Find model 42 (0-indexed)
    if len(run_011) > 42:
        m42 = run_011[42]
        print(f"\nModel 42 (0-indexed): TM(Fold1)={m42[0]}, TM(Fold2)={m42[1]}")
        print(f"\nCLAIM: TM(Fold1)=0.50, TM(Fold2)=0.37")
        print(f"ACTUAL: TM(Fold1)={m42[0]}, TM(Fold2)={m42[1]}")
    
    # Find max Fold2 in this run
    max_f2_in_011 = max(enumerate(run_011), key=lambda x: x[1][1])
    print(f"\nMax Fold2 in run 0_011: model {max_f2_in_011[0]}, TM2={max_f2_in_011[1][1]}, TM1={max_f2_in_011[1][0]}")
    
    # Find "hesitant" predictions (low Fold1)
    hesitant = [(i, tm1, tm2) for i, (tm1, tm2) in enumerate(run_011) if tm1 < 0.75]
    print(f"\n'Hesitant' predictions in 0_011 (TM1 < 0.75): {len(hesitant)}")
    for idx, tm1, tm2 in sorted(hesitant, key=lambda x: x[1]):
        print(f"  Model {idx}: TM1={tm1}, TM2={tm2}")

# Global analysis: hesitant vs confident
print(f"\n=== HESITANT SUBSET ANALYSIS ===")
hesitant_all = [p for p in all_predictions if p['tm_fold1'] < 0.75]
confident_all = [p for p in all_predictions if p['tm_fold1'] >= 0.75]
print(f"Hesitant (TM1 < 0.75): {len(hesitant_all)} predictions")
print(f"Confident (TM1 >= 0.75): {len(confident_all)} predictions")

if hesitant_all:
    max_f2_hesitant = max(hesitant_all, key=lambda x: x['tm_fold2'])
    print(f"\nMax Fold2 in hesitant subset: {max_f2_hesitant['tm_fold2']}")
    print(f"  Run: {max_f2_hesitant['run']}, Model: {max_f2_hesitant['model_idx']}")
    print(f"  TM(Fold1): {max_f2_hesitant['tm_fold1']}")

if confident_all:
    max_f2_confident = max(confident_all, key=lambda x: x['tm_fold2'])
    print(f"\nMax Fold2 in confident subset: {max_f2_confident['tm_fold2']}")
    print(f"  Run: {max_f2_confident['run']}, Model: {max_f2_confident['model_idx']}")
    
# Find any prediction with Fold2 >= 0.37
print(f"\n=== CHECKING FOR TM(Fold2) >= 0.37 ===")
high_f2 = [p for p in all_predictions if p['tm_fold2'] >= 0.37]
print(f"Predictions with TM(Fold2) >= 0.37: {len(high_f2)}")
for p in high_f2[:10]:
    print(f"  Run {p['run']}, Model {p['model_idx']}: TM1={p['tm_fold1']}, TM2={p['tm_fold2']}")
```
