## supplement_v6 for "Confidence Without Verification: Screening pLDDT Unreliability in AlphaFold2 Fold-Switching Predictions": vpc_porter_analysis.py

```
#!/usr/bin/env python3
"""
V_PC Analysis Pipeline for Porter Fold-Switching Benchmark

Computes V_PC (Preservation-Coherence violation) scores for AlphaFold2 predictions
on the 93 fold-switching proteins from Porter et al.

V_PC detects: High confidence predictions that disagree across MSA sampling
This signals epistemic overconfidence - the model is confident but wrong.

Author: Robin (CPE Framework)
Date: 2026-01-14
"""

import json
import os
import glob
import numpy as np
from pathlib import Path
from dataclasses import dataclass
from typing import List, Dict, Tuple, Optional
import re


@dataclass
class ClusterPrediction:
    """Predictions from a single MSA cluster"""
    cluster_name: str
    models: List[dict]  # List of {plddt, pae, ptm, max_pae} for each model
    tm_fold1: List[float]  # TM-scores to Fold1 for each model
    tm_fold2: List[float]  # TM-scores to Fold2 for each model
    
    @property
    def mean_ptm(self) -> float:
        return np.mean([m['ptm'] for m in self.models])
    
    @property
    def std_ptm(self) -> float:
        return np.std([m['ptm'] for m in self.models])
    
    @property
    def mean_plddt(self) -> float:
        return np.mean([np.mean(m['plddt']) for m in self.models])
    
    @property
    def dominant_fold(self) -> int:
        """Returns 1 or 2 based on which fold has higher mean TM-score"""
        mean_tm1 = np.mean(self.tm_fold1)
        mean_tm2 = np.mean(self.tm_fold2)
        return 1 if mean_tm1 > mean_tm2 else 2
    
    @property 
    def fold_confidence(self) -> float:
        """How strongly does this cluster favor one fold over other"""
        mean_tm1 = np.mean(self.tm_fold1)
        mean_tm2 = np.mean(self.tm_fold2)
        return abs(mean_tm1 - mean_tm2)


@dataclass
class ProteinAnalysis:
    """Complete V_PC analysis for one protein"""
    protein_id: str
    clusters: List[ClusterPrediction]
    is_af2_success: bool  # Did AF2 find Fold2?
    
    @property
    def mean_ptm_all(self) -> float:
        """Mean pTM across all clusters and models"""
        all_ptms = []
        for c in self.clusters:
            all_ptms.extend([m['ptm'] for m in c.models])
        return np.mean(all_ptms)
    
    @property
    def cross_cluster_agreement(self) -> float:
        """Fraction of clusters agreeing on dominant fold"""
        if len(self.clusters) == 0:
            return 1.0
        folds = [c.dominant_fold for c in self.clusters]
        mode_fold = max(set(folds), key=folds.count)
        return folds.count(mode_fold) / len(folds)
    
    @property
    def cross_cluster_ptm_std(self) -> float:
        """Std of mean pTM across clusters - high = inconsistent confidence"""
        if len(self.clusters) <= 1:
            return 0.0
        cluster_ptms = [c.mean_ptm for c in self.clusters]
        return np.std(cluster_ptms)
    
    @property
    def vpc_score(self) -> float:
        """
        V_PC Score: Detects high confidence + low coherence
        
        High score = Model is confident but clusters disagree
        This predicts AF2 failures (false positives)
        
        V_PC = mean_confidence × (1 - agreement) × fold_ambiguity
        """
        # Component 1: Mean confidence (pTM)
        confidence = self.mean_ptm_all
        
        # Component 2: Disagreement across clusters
        disagreement = 1.0 - self.cross_cluster_agreement
        
        # Component 3: Fold ambiguity - do predictions sit near decision boundary?
        fold_margins = [c.fold_confidence for c in self.clusters]
        mean_margin = np.mean(fold_margins) if fold_margins else 0.5
        ambiguity = 1.0 - min(mean_margin, 0.3) / 0.3  # Normalize: margin < 0.3 is ambiguous
        
        # V_PC: High when confident but incoherent
        return confidence * (0.5 * disagreement + 0.5 * ambiguity)
    
    @property
    def vpc_score_simple(self) -> float:
        """
        Simplified V_PC: Just confidence × disagreement
        """
        return self.mean_ptm_all * (1.0 - self.cross_cluster_agreement)


def parse_tmscore_line(line: str) -> Tuple[str, List[Tuple[float, float]]]:
    """
    Parse tmscores.txt line format:
    'cluster_name : tm1 tm2,tm1 tm2,...'
    
    Returns (cluster_name, [(tm_fold1, tm_fold2), ...])
    """
    parts = line.strip().split(' : ')
    if len(parts) != 2:
        return None, []
    
    cluster_name = parts[0].strip()
    scores_str = parts[1].strip()
    
    pairs = []
    for pair in scores_str.split(','):
        pair = pair.strip()
        if not pair:
            continue
        vals = pair.split()
        if len(vals) >= 2:
            try:
                pairs.append((float(vals[0]), float(vals[1])))
            except ValueError:
                continue
    
    return cluster_name, pairs


def load_protein_data(protein_dir: str) -> Optional[ProteinAnalysis]:
    """
    Load all prediction data for a single protein directory
    """
    protein_id = os.path.basename(protein_dir)
    
    # Load TM-scores
    tmscores_file = os.path.join(protein_dir, 'tmscores.txt')
    tmscores_fs_file = os.path.join(protein_dir, 'tmscores_fs.txt')
    
    if not os.path.exists(tmscores_file):
        print(f"  Warning: No tmscores.txt for {protein_id}")
        return None
    
    # Parse TM-scores by cluster
    cluster_tmscores = {}
    with open(tmscores_file, 'r') as f:
        for line in f:
            cluster_name, pairs = parse_tmscore_line(line)
            if cluster_name:
                cluster_tmscores[cluster_name] = pairs
    
    if not cluster_tmscores:
        print(f"  Warning: Empty tmscores for {protein_id}")
        return None
    
    # Load predictions for each cluster
    clusters = []
    for cluster_name, tm_pairs in cluster_tmscores.items():
        cluster_dir = os.path.join(protein_dir, cluster_name)
        
        if not os.path.exists(cluster_dir):
            # Try without leading zeros or other formats
            continue
        
        # Find score JSON files
        score_files = glob.glob(os.path.join(cluster_dir, '*_scores_rank_*.json'))
        
        if not score_files:
            continue
        
        # Load each model's scores
        models = []
        for sf in sorted(score_files)[:5]:  # Max 5 models per cluster
            try:
                with open(sf, 'r') as f:
                    data = json.load(f)
                    models.append({
                        'plddt': data.get('plddt', []),
                        'pae': data.get('pae', []),
                        'ptm': data.get('ptm', 0.0),
                        'max_pae': data.get('max_pae', 30.0)
                    })
            except (json.JSONDecodeError, IOError) as e:
                continue
        
        if models and len(tm_pairs) >= len(models):
            tm_fold1 = [p[0] for p in tm_pairs[:len(models)]]
            tm_fold2 = [p[1] for p in tm_pairs[:len(models)]]
            
            clusters.append(ClusterPrediction(
                cluster_name=cluster_name,
                models=models,
                tm_fold1=tm_fold1,
                tm_fold2=tm_fold2
            ))
    
    if not clusters:
        print(f"  Warning: No valid clusters loaded for {protein_id}")
        return None
    
    # Determine if AF2 succeeded (found Fold2)
    # Success = any model has TM-score > 0.5 to Fold2
    best_tm_fold2 = max(
        max(c.tm_fold2) for c in clusters
    )
    is_success = best_tm_fold2 > 0.5
    
    return ProteinAnalysis(
        protein_id=protein_id,
        clusters=clusters,
        is_af2_success=is_success
    )


def compute_enrichment(proteins: List[ProteinAnalysis], k: int = 15) -> dict:
    """
    Compute enrichment of failures in top-K by V_PC score
    
    Enrichment = (failures in top-K / K) / (total failures / total)
    """
    # Sort by V_PC score descending
    sorted_proteins = sorted(proteins, key=lambda p: p.vpc_score, reverse=True)
    
    total_failures = sum(1 for p in proteins if not p.is_af2_success)
    total = len(proteins)
    baseline_rate = total_failures / total if total > 0 else 0
    
    top_k = sorted_proteins[:k]
    failures_in_top_k = sum(1 for p in top_k if not p.is_af2_success)
    top_k_rate = failures_in_top_k / k if k > 0 else 0
    
    enrichment = top_k_rate / baseline_rate if baseline_rate > 0 else 0
    
    return {
        'k': k,
        'total_proteins': total,
        'total_failures': total_failures,
        'baseline_failure_rate': baseline_rate,
        'failures_in_top_k': failures_in_top_k,
        'top_k_failure_rate': top_k_rate,
        'enrichment': enrichment,
        'top_k_proteins': [(p.protein_id, p.vpc_score, p.is_af2_success) for p in top_k]
    }


def load_success_list(excel_path: str) -> set:
    """
    Load list of proteins where AF2 succeeded from Excel file
    Returns set of protein IDs that AF2 found Fold2 for
    """
    try:
        import pandas as pd
        df = pd.read_excel(excel_path)
        # The success file has columns for different AF2 versions
        # We use AF2.2.0 or All_AF column
        # For now, return empty - will determine from TM-scores directly
        return set()
    except Exception as e:
        print(f"  Could not load success list: {e}")
        return set()


def run_full_analysis(data_dir: str, output_path: str = None):
    """
    Run V_PC analysis on all proteins in data directory
    """
    print("=" * 60)
    print("V_PC Analysis: Porter Fold-Switching Benchmark")
    print("=" * 60)
    print(f"\nData directory: {data_dir}")
    
    # Find all protein directories
    protein_dirs = []
    for item in os.listdir(data_dir):
        item_path = os.path.join(data_dir, item)
        if os.path.isdir(item_path):
            # Check if it has tmscores.txt
            if os.path.exists(os.path.join(item_path, 'tmscores.txt')):
                protein_dirs.append(item_path)
    
    print(f"Found {len(protein_dirs)} protein directories with tmscores.txt")
    
    # Load all proteins
    proteins = []
    for pdir in sorted(protein_dirs):
        protein = load_protein_data(pdir)
        if protein:
            proteins.append(protein)
            print(f"  Loaded {protein.protein_id}: {len(protein.clusters)} clusters, "
                  f"pTM={protein.mean_ptm_all:.3f}, agreement={protein.cross_cluster_agreement:.2f}, "
                  f"V_PC={protein.vpc_score:.3f}")
    
    print(f"\nSuccessfully loaded {len(proteins)} proteins")
    
    if not proteins:
        print("No proteins loaded. Check data directory structure.")
        return
    
    # Summary statistics
    print("\n" + "=" * 60)
    print("Summary Statistics")
    print("=" * 60)
    
    successes = [p for p in proteins if p.is_af2_success]
    failures = [p for p in proteins if not p.is_af2_success]
    
    print(f"AF2 Successes (found Fold2): {len(successes)}")
    print(f"AF2 Failures (missed Fold2): {len(failures)}")
    print(f"Failure rate: {len(failures)/len(proteins)*100:.1f}%")
    
    # V_PC distribution
    vpc_scores = [p.vpc_score for p in proteins]
    print(f"\nV_PC Score Distribution:")
    print(f"  Mean: {np.mean(vpc_scores):.3f}")
    print(f"  Std:  {np.std(vpc_scores):.3f}")
    print(f"  Min:  {np.min(vpc_scores):.3f}")
    print(f"  Max:  {np.max(vpc_scores):.3f}")
    
    # Compare V_PC for successes vs failures
    if successes:
        vpc_success = [p.vpc_score for p in successes]
        print(f"\nV_PC for Successes: mean={np.mean(vpc_success):.3f}, std={np.std(vpc_success):.3f}")
    if failures:
        vpc_failure = [p.vpc_score for p in failures]
        print(f"V_PC for Failures:  mean={np.mean(vpc_failure):.3f}, std={np.std(vpc_failure):.3f}")
    
    # Enrichment analysis
    print("\n" + "=" * 60)
    print("Enrichment Analysis")
    print("=" * 60)
    
    for k in [5, 10, 15, 20, 25]:
        if k > len(proteins):
            continue
        result = compute_enrichment(proteins, k)
        print(f"\nK={k}:")
        print(f"  Failures in top-{k}: {result['failures_in_top_k']}/{k}")
        print(f"  Enrichment: {result['enrichment']:.2f}x")
        
        if k == 15:
            print(f"\n  Top-15 proteins by V_PC:")
            for pid, vpc, is_fail in result['top_k_proteins']:
                status = "FAILURE" if not is_fail else "success"
                print(f"    {pid}: V_PC={vpc:.3f} [{status}]")
    
    # Output results
    print("\n" + "=" * 60)
    print("Full Rankings (by V_PC descending)")
    print("=" * 60)
    
    sorted_proteins = sorted(proteins, key=lambda p: p.vpc_score, reverse=True)
    
    results_data = []
    for i, p in enumerate(sorted_proteins, 1):
        status = "FAILURE" if not p.is_af2_success else "success"
        results_data.append({
            'rank': i,
            'protein_id': p.protein_id,
            'vpc_score': p.vpc_score,
            'mean_ptm': p.mean_ptm_all,
            'cross_cluster_agreement': p.cross_cluster_agreement,
            'num_clusters': len(p.clusters),
            'af2_success': p.is_af2_success
        })
        if i <= 30:
            print(f"{i:3d}. {p.protein_id:12s} V_PC={p.vpc_score:.3f} "
                  f"pTM={p.mean_ptm_all:.2f} agree={p.cross_cluster_agreement:.2f} "
                  f"[{status}]")
    
    # Save to CSV if output path provided
    if output_path:
        try:
            import pandas as pd
            df = pd.DataFrame(results_data)
            df.to_csv(output_path, index=False)
            print(f"\nResults saved to: {output_path}")
        except ImportError:
            # Save as plain CSV
            with open(output_path, 'w') as f:
                f.write("rank,protein_id,vpc_score,mean_ptm,cross_cluster_agreement,num_clusters,af2_success\n")
                for r in results_data:
                    f.write(f"{r['rank']},{r['protein_id']},{r['vpc_score']:.4f},"
                            f"{r['mean_ptm']:.4f},{r['cross_cluster_agreement']:.4f},"
                            f"{r['num_clusters']},{r['af2_success']}\n")
            print(f"\nResults saved to: {output_path}")
    
    return proteins, results_data


def main():
    """
    Main analysis pipeline
    """
    import argparse
    
    parser = argparse.ArgumentParser(description='V_PC Analysis for Porter Fold-Switching Benchmark')
    parser.add_argument('--data', '-d', type=str, 
                        default=r'C:\Users\robin\Desktop\substrate_agnostic_experiments\BlindFold\AF2_benchmark-AF2-benchmark\porterll-AF2_benchmark-40a57d7\data',
                        help='Path to data directory containing protein folders')
    parser.add_argument('--output', '-o', type=str, default='vpc_results.csv',
                        help='Output CSV path')
    
    args = parser.parse_args()
    
    if os.path.exists(args.data):
        run_full_analysis(args.data, args.output)
    else:
        print(f"Data directory not found: {args.data}")
        print("\nUsage: python vpc_porter_analysis.py --data /path/to/data --output results.csv")
        print("\nExpected structure:")
        print("  data/")
        print("    protein_id/")
        print("      tmscores.txt")
        print("      cluster_name/")
        print("        *_scores_rank_*.json")


if __name__ == '__main__':
    main()
```
