## supplement_v6 for "Confidence Without Verification: Screening pLDDT Unreliability in AlphaFold2 Fold-Switching Predictions": vpc_porter_complete.py

```
#!/usr/bin/env python3
"""
V_PC Analysis for Porter Fold-Switching Benchmark

Uses:
- TableS1.xlsx: Full list of 93 fold-switching proteins  
- success_using_TMscore_metric_revised.xlsx: Success/failure labels per method
- tmscores_fs_all.npy: Aggregated fold-switching TM-scores

Strategy:
Since we don't have per-protein pTM confidence scores in the aggregated data,
we use TM-score VARIANCE as a proxy for V_PC:
- High variance in TM-scores across predictions = disagreement
- Combined with mean TM-score gives a V_PC-like signal
"""

import numpy as np
import pandas as pd
import os

def load_protein_list(tables1_path):
    """Load 93 proteins from TableS1.xlsx"""
    df = pd.read_excel(tables1_path)
    proteins = []
    for _, row in df.iterrows():
        # Use Fold2 as the protein ID (matches data directory naming like 2oug_C)
        fold1 = str(row['Fold1']).replace('_', '')  # 1h38D
        fold2 = str(row['Fold2']).replace('_', '')  # 1qlnA
        # Convert to directory format (e.g., 2oug_C)
        proteins.append({
            'fold1': fold1,
            'fold2': fold2,
            'fold1_fmt': f"{fold1[:-1]}_{fold1[-1]}",  # 1h38_D
            'fold2_fmt': f"{fold2[:-1]}_{fold2[-1]}",  # 1qln_A
        })
    return proteins

def load_successes(success_path):
    """Load success lists from all sheets"""
    xl = pd.ExcelFile(success_path)
    successes = {}
    
    for sheet in xl.sheet_names:
        if sheet == 'counts':
            continue
        df = pd.read_excel(xl, sheet_name=sheet)
        # Get PDB1 and PDB2 columns
        success_pairs = set()
        for _, row in df.iterrows():
            pdb1 = str(row.get('PDB1', '')).strip()
            pdb2 = str(row.get('PDB2', '')).strip()
            if pdb1 and pdb2 and pdb1 != 'nan':
                # Normalize format
                pdb1 = pdb1.replace('_', '')
                pdb2 = pdb2.replace('_', '')
                success_pairs.add((pdb1, pdb2))
                success_pairs.add((pdb2, pdb1))  # Either direction
        successes[sheet] = success_pairs
    
    return successes

def analyze_tmscore_variance(tmscores_fs, n_proteins=93, preds_per_protein=None):
    """
    Analyze TM-score variance as V_PC proxy
    
    The tmscores_fs array contains alternating (TM_fold1, TM_fold2) pairs
    """
    # Reshape to pairs
    n_pairs = len(tmscores_fs) // 2
    fs_pairs = tmscores_fs[:n_pairs*2].reshape(-1, 2)
    
    if preds_per_protein is None:
        # Estimate predictions per protein
        preds_per_protein = n_pairs // n_proteins
    
    print(f"Total pairs: {n_pairs}")
    print(f"Estimated predictions per protein: {preds_per_protein}")
    
    results = []
    for p in range(n_proteins):
        start = p * preds_per_protein
        end = min(start + preds_per_protein, len(fs_pairs))
        
        if start >= len(fs_pairs):
            break
            
        protein_preds = fs_pairs[start:end]
        
        if len(protein_preds) == 0:
            continue
        
        # Metrics
        tm_fold1 = protein_preds[:, 0]
        tm_fold2 = protein_preds[:, 1]
        
        # Dominant fold per prediction
        dominant_folds = [1 if t1 > t2 else 2 for t1, t2 in protein_preds]
        fold1_count = dominant_folds.count(1)
        fold2_count = dominant_folds.count(2)
        
        # Agreement (coherence proxy)
        agreement = max(fold1_count, fold2_count) / len(dominant_folds)
        
        # Mean confidence proxy (max TM per prediction)  
        max_tms = np.array([max(t1, t2) for t1, t2 in protein_preds])
        mean_confidence = np.mean(max_tms)
        
        # TM-score variance (disagreement signal)
        tm1_var = np.var(tm_fold1)
        tm2_var = np.var(tm_fold2)
        total_var = tm1_var + tm2_var
        
        # V_PC proxy: high confidence × low agreement
        vpc_proxy = mean_confidence * (1 - agreement)
        
        # Alternative V_PC: variance-weighted
        vpc_variance = mean_confidence * np.sqrt(total_var)
        
        results.append({
            'protein_idx': p,
            'n_predictions': len(protein_preds),
            'mean_tm_fold1': np.mean(tm_fold1),
            'mean_tm_fold2': np.mean(tm_fold2),
            'std_tm_fold1': np.std(tm_fold1),
            'std_tm_fold2': np.std(tm_fold2),
            'fold1_predictions': fold1_count,
            'fold2_predictions': fold2_count,
            'agreement': agreement,
            'mean_confidence': mean_confidence,
            'vpc_agreement': vpc_proxy,
            'vpc_variance': vpc_variance,
        })
    
    return pd.DataFrame(results)

def main():
    # Paths - adjust as needed
    tables1_path = r'C:\Users\robin\Desktop\substrate_agnostic_experiments\BlindFold\AF2_benchmark-AF2-benchmark\porterll-AF2_benchmark-40a57d7\TableS1.xlsx'
    success_path = r'C:\Users\robin\Desktop\substrate_agnostic_experiments\BlindFold\AF2_benchmark-AF2-benchmark\porterll-AF2_benchmark-40a57d7\success_using_TMscore_metric_revised.xlsx'
    data_dir = r'C:\Users\robin\Desktop\substrate_agnostic_experiments\BlindFold\AF2_benchmark-AF2-benchmark\porterll-AF2_benchmark-40a57d7\data'
    
    # Alternative paths (if files are in uploads)
    if not os.path.exists(tables1_path):
        tables1_path = 'TableS1.xlsx'  
    if not os.path.exists(success_path):
        success_path = 'success_using_TMscore_metric_revised.xlsx'
    
    print("=" * 70)
    print("V_PC Analysis: Porter Fold-Switching Benchmark")
    print("=" * 70)
    
    # Load protein list
    print("\n--- Loading protein list ---")
    proteins = load_protein_list(tables1_path)
    print(f"Loaded {len(proteins)} proteins from TableS1")
    print(f"First 5: {[p['fold2'] for p in proteins[:5]]}")
    
    # Load success data
    print("\n--- Loading success/failure labels ---")
    successes = load_successes(success_path)
    for method, pairs in successes.items():
        print(f"  {method}: {len(pairs)//2} successes")
    
    # Create success labels for each protein
    for p in proteins:
        p['af2_success'] = (p['fold1'], p['fold2']) in successes.get('AF2.3.1', set())
        p['all_af_success'] = (p['fold1'], p['fold2']) in successes.get('AF_cluster colabfold1.5.2', set())
        p['ace_success'] = (p['fold1'], p['fold2']) in successes.get('ACE', set())
    
    af2_failures = sum(1 for p in proteins if not p['af2_success'])
    print(f"\nAF2.3.1 failures: {af2_failures}/93 = {af2_failures/93*100:.1f}%")
    
    # Load TM-score data
    print("\n--- Loading TM-score data ---")
    tmscores_fs = np.load(os.path.join(data_dir, 'tmscores_fs_all.npy'))
    print(f"tmscores_fs_all.npy: {len(tmscores_fs)} values")
    
    # Analyze with variance proxy
    print("\n--- Computing V_PC proxy scores ---")
    # 212018 values / 2 = 106009 pairs / 93 proteins ≈ 1140 predictions each
    # That's a lot! Let's compute with that estimate
    n_pairs = len(tmscores_fs) // 2
    preds_per_protein = n_pairs // 93
    
    results_df = analyze_tmscore_variance(tmscores_fs, n_proteins=93, 
                                          preds_per_protein=preds_per_protein)
    
    # Merge with protein info
    for i, row in results_df.iterrows():
        if i < len(proteins):
            results_df.loc[i, 'fold1'] = proteins[i]['fold1']
            results_df.loc[i, 'fold2'] = proteins[i]['fold2']
            results_df.loc[i, 'af2_success'] = proteins[i]['af2_success']
    
    # Filter to valid entries
    results_df = results_df[results_df['fold1'].notna()].copy()
    
    print(f"\nAnalyzed {len(results_df)} proteins")
    
    # Summary statistics
    print("\n" + "=" * 70)
    print("V_PC Proxy Distribution")
    print("=" * 70)
    
    print(f"\nV_PC (agreement-based):")
    print(f"  Mean: {results_df['vpc_agreement'].mean():.4f}")
    print(f"  Std:  {results_df['vpc_agreement'].std():.4f}")
    print(f"  Max:  {results_df['vpc_agreement'].max():.4f}")
    
    print(f"\nV_PC (variance-based):")
    print(f"  Mean: {results_df['vpc_variance'].mean():.4f}")
    print(f"  Std:  {results_df['vpc_variance'].std():.4f}")
    print(f"  Max:  {results_df['vpc_variance'].max():.4f}")
    
    # Compare successes vs failures
    successes_df = results_df[results_df['af2_success'] == True]
    failures_df = results_df[results_df['af2_success'] == False]
    
    if len(successes_df) > 0 and len(failures_df) > 0:
        print("\n" + "=" * 70)
        print("V_PC: Successes vs Failures")
        print("=" * 70)
        print(f"\nAF2 Successes (n={len(successes_df)}):")
        print(f"  V_PC agreement: {successes_df['vpc_agreement'].mean():.4f} ± {successes_df['vpc_agreement'].std():.4f}")
        print(f"  V_PC variance:  {successes_df['vpc_variance'].mean():.4f} ± {successes_df['vpc_variance'].std():.4f}")
        
        print(f"\nAF2 Failures (n={len(failures_df)}):")
        print(f"  V_PC agreement: {failures_df['vpc_agreement'].mean():.4f} ± {failures_df['vpc_agreement'].std():.4f}")
        print(f"  V_PC variance:  {failures_df['vpc_variance'].mean():.4f} ± {failures_df['vpc_variance'].std():.4f}")
    
    # Enrichment analysis
    print("\n" + "=" * 70)
    print("Enrichment Analysis")
    print("=" * 70)
    
    total = len(results_df)
    total_failures = len(failures_df)
    baseline_rate = total_failures / total if total > 0 else 0
    
    for vpc_col in ['vpc_agreement', 'vpc_variance']:
        print(f"\n--- Using {vpc_col} ---")
        sorted_df = results_df.sort_values(vpc_col, ascending=False)
        
        for k in [5, 10, 15, 20, 25]:
            if k > total:
                continue
            top_k = sorted_df.head(k)
            failures_in_k = len(top_k[top_k['af2_success'] == False])
            rate_k = failures_in_k / k
            enrichment = rate_k / baseline_rate if baseline_rate > 0 else 0
            
            print(f"  K={k:2d}: {failures_in_k}/{k} failures, enrichment = {enrichment:.2f}x")
    
    # Top 15 by V_PC
    print("\n" + "=" * 70)
    print("Top 15 proteins by V_PC (agreement-based)")
    print("=" * 70)
    
    sorted_df = results_df.sort_values('vpc_agreement', ascending=False)
    for i, row in sorted_df.head(15).iterrows():
        status = "FAILURE" if not row['af2_success'] else "success"
        print(f"  {row['fold1']}-{row['fold2']}: V_PC={row['vpc_agreement']:.4f} "
              f"agree={row['agreement']:.2f} conf={row['mean_confidence']:.2f} [{status}]")
    
    # Save results
    output_path = 'vpc_porter_results.csv'
    results_df.to_csv(output_path, index=False)
    print(f"\nResults saved to: {output_path}")


if __name__ == '__main__':
    main()
```
